# Instructions and experiential learning have similar impacts on pain and pain-related brain responses but produce dissociations in value-based reversal learning

**DOI:** 10.1101/2021.08.25.457682

**Authors:** Lauren Y. Atlas, Troy C. Dildine, Esther E. Palacios-Barrios, Qingbao Yu, Richard C. Reynolds, Lauren A. Banker, Shara S. Grant, Daniel S. Pine

**Affiliations:** National Center for Complementary and Integrative Health, National Institutes of Health, Bethesda, MD; National Institute on Drug Abuse, National Institutes of Health, Baltimore, MD; National Institute of Mental Health, National Institutes of Health, Bethesda, MD; Department of Psychology, University of Pittsburgh, Pittsburgh, PA; Department of Clinical Neuroscience, Karolinska Institutet, Solna, Sweden; Department of Psychology, University of California – Los Angeles, Los Angeles, CA

**Keywords:** pain, reversal learning, expectancy, conditioning, fMRI, computational modeling, prediction

## Abstract

Recent data suggest that interactions between systems involved in higher order knowledge and associative learning drive responses during appetitive and aversive learning. However, it is unknown how these systems impact subjective responses, such as pain. We tested how instructions and reversal learning influence pain and pain-evoked brain activation. Healthy volunteers (n = 40) were either instructed about contingencies between cues and aversive outcomes or learned through experience in a paradigm where contingencies reversed three times. We measured predictive cue effects on pain and heat-evoked brain responses using functional magnetic resonance imaging. Predictive cues dynamically modulated pain perception as contingencies changed, regardless of whether participants received contingency instructions. Heat-evoked responses in the insula, anterior cingulate, and putamen updated as contingencies changed, whereas the periaqueductal gray and thalamus responded to initial contingencies throughout the task. Quantitative modeling revealed that expected value was shaped purely by instructions in the Instructed Group, whereas expected value updated dynamically in the Uninstructed Group as a function of error-based learning. These differences were accompanied by dissociations in the neural correlates of value-based learning in the rostral anterior cingulate, medial prefrontal cortex, and orbitofrontal cortex. These results show how predictions impact subjective pain. Moreover, imaging data delineate three types of networks involved in pain generation and value-based learning: those that respond to initial contingencies, those that update dynamically during feedback-driven learning as contingencies change, and those that are sensitive to instruction. Together, these findings provide multiple points of entry for therapies designs to impact pain.

---

Predictions and expectations shape perception across many domains, through processes such as predictive coding. This is particularly apparent in the context of pain as evidenced by data on placebo analgesia and expectancy-based pain modulation (Büchel et al., 2014; Ongaro and Kaptchuk, 2018; Kaptchuk et al., 2020). While most studies of predictive coding examine probabilistic error-driven learning, humans also use verbal instructions to shape predictions, with instructions acting either alone or through effects on learning (for reviews, see (Koban et al., 2017; Mertens et al., 2018; Atlas, 2019)). However, it is unknown how instructions and learning combine dynamically to shape pain and pain-related brain responses. We introduced a novel pain reversal learning task to measure the dynamic effects of predictive cues on subjective pain and brain responses to noxious heat. We used this task to study how responding is affected by task instructions and experiential learning.

Placebo effects are thought to depend on expectations formed through conditioning or associative learning (e.g. prior treatment experiences) as well as verbal instruction and explicit knowledge (e.g. the doctor’s instruction). Most studies of placebo mechanisms combine suggestion and conditioning to maximize expectations and measure downstream responses.

These experiments indicate that placebos reliably reduce acute pain (Forsberg et al., 2017; Zunhammer et al., 2018) and alter stimulus-evoked responses in multiple brain regions, including the insula, dorsal anterior cingulate, and thalamus, as well as pain modulatory regions including the opioid-rich periaqueductal gray (PAG), the dorsolateral prefrontal cortex (DLPFC), and the rostral anterior cingulate cortex (rACC) (Atlas and Wager, 2014). While placebo effects are influenced by both instructions and experiential learning, many studies have sought to dissociate their effects (Montgomery and Kirsch, 1996; Benedetti et al., 2003; Colloca et al., 2008a, 2008b). For example, in one study (Benedetti et al., 2003) participants underwent several days of conditioning with active treatments for pain, motor performance in Parkinson’s disease, or drugs that affect hormonal responses (cortisol or growth hormone). Participants subsequently received verbal instructions that they would receive a drug that leads to the opposite effect of conditioning. All participants actually received placebo. Placebo effects on outcomes that could be consciously monitored (pain and motor responding) reversed with instruction, while hormonal responses continued to mimic conditioning. Other studies indicate that instructions only reverse placebo analgesia after brief conditioning (Scott M Schafer et al., 2015). Thus, placebo effects on specific outcomes manifest unique sensitivities to instructed knowledge alone or through effects on learning.

These studies illustrate that instructed reversals can distinguish between purely associative processes and those that are sensitive to higher-order knowledge. This connects placebo with an established literature on how instructions influence appetitive and aversive learning (Grings, 1973; McNally, 1981; Costa et al., 2015; Mertens and De Houwer, 2016; Atlas, 2019). However, despite a large body of work on the neurobiology of placebo analgesia, we lack a mechanistic understanding of how instructions and learning modify pain. This at least partly reflects the practice of combining these factors to maximize expectations. Yet cognitive neuroscience indicates that distinct processes may support instructions and learning. In traditional dual systems frameworks, frontal regions including the DLPFC maintain goals and higher order knowledge, while subcortical systems including the striatum, amygdala, and the orbitofrontal / ventromedial prefrontal cortex (OFC/VMPFC) support value-based associative learning. These dissociations are supported by lesion work (Bechara et al., 1995; Clark, 1998) and both systems have been implicated in neuroimaging studies of placebo (Atlas and Wager, 2014). However, reinforcement learning experiments indicate that instructions can shape reward learning, and that this occurs through interactions between the DLPFC and striatum (Doll et al., 2009, 2011; Li et al., 2011a). We previously showed that corticostriatal interactions also support the effect of instructions on aversive learning, but that the amygdala learned from aversive outcomes irrespective of instruction (Atlas et al., 2016; Atlas, 2019). This provides a potential mechanism by which some outcomes may continue to respond to associative learning in spite of instructions, while others may update with instruction, consistent with dissociations observed in previous work (Benedetti et al., 2003). Importantly, most previous work on how instructions shape learning has measured autonomic responses during classical conditioning or binary choices in instrumental learning tasks. Acute pain tasks provide a unique opportunity to measure how learning and instructions shape conscious, subjective decisions, which may be distinct from autonomic responses or instrumental choice.

We asked how instructions and learning combine to dynamically shape pain and pain-related brain responses. Participants underwent a pain reversal learning task and were assigned to an Instructed Group, who was informed about contingencies and reversals, or an Uninstructed Group, who learned purely through experience. We used multilevel mediation analysis to identify brain regions that are modulated by instructions or learning and modulate subjective pain. We also fit computational models of instructed learning (Atlas et al., 2016, 2019) to pain ratings to determine how instructions and associative learning dynamically shape pain, and to isolate brain regions that track expected value during pain reversal learning. We were most interested in understanding how instructions and learning affect brain responses within brain networks involved in pain and value-based learning. We hypothesized that instructions and learning would both dynamically shape pain, and that instructed reversals would lead to immediate reversals of pain reports and heat-evoked brain responses in the DLPFC and pain processing network.

## Methods

### Participants

49 participants (25 female, *M*age= 28.04 years, SDage = 7.04) were recruited and consented to participate in an fMRI study designed to measure “how pain and emotions are processed in the human brain and influenced by psychological factors.” Participants provided informed consent in accordance with the Declaration of Helsinki, and the protocol was approved by the NIH’s Combined Neuroscience Institutional Review Board (Protocol 15-AT-0132, PI: Atlas).

Participants were eligible to participate if they were between 18 and 50, fluent in English, healthy (i.e. had no medical conditions that affect pain or somatosensation, no psychiatric, neurological, autonomic, or cardiovascular disorders, no chronic systemic diseases, and no medication that can affect pain perception), right-handed, and had received a medical exam at NIH within the previous year. All participants underwent urine toxicology testing to ensure they had not used recreational drugs that alter pain. Participants were drawn from a pool of subjects who had completed an initial screening visit that tested whether participants reliably reported increased pain with increased temperatures (r^2^>0.4) and exhibited pain tolerance at or below 50℃ (the maximum temperature we applied during the study). 9 participants who provided consent did not complete the experiment due to ineligibility based on calibration (n = 4) technical failures (n = 1), compliance with procedures (n = 2), or anatomical abnormalities identified in a clinical scan (n = 2) and were not included in the current analyses. The final sample included 40 participants (22 female; *M*age= 27.00 years, SDage = 6.21).

## Materials and procedure

### Stimuli and apparatus

We delivered thermal stimulation to the left (non-dominant) volar forearm using a 16 x 16 ATS contact heat thermode controlled with a Pathway pain and sensory evaluation system (Medoc Ltd, Ramat Yisha, Israel). Each heat stimulus lasted 8 seconds and consisted of 3 phases: a 1.5 second on-ramp phase in which the temperature of the thermode rose from 32°C to the target temperature level, a 5-second plateau phase in which target temperature was maintained, and a 1.5 second off-ramp phase in which the temperature returned to 32°C. Thermode placement was adjusted between each block of trials (i.e. every 12 trials) to avoid sensitization, habituation, and skin damage. Temperatures ranged from 36°C to 50°C, in increments of 0.5°C, and were selected based on a thermal pain calibration conducted immediately prior to the experiment. Thermode temperature was maintained at 32°C between trials.

Experiment Builder (SR-Research, Ontario, Canada) was used to deliver visual and auditory stimuli, to trigger noxious stimulation on the Pathways computer, and to synchronize task timing with physiological recording. Physiological data, including electrodermal activity (EDA), respiration, electrocardiography, and peripheral pulse, were recorded from the left hand using Biopac recording equipment and accompanying AcqKnowledge software (Goleta, CA). Participants recorded pain ratings using a trackball with their right hand while EDA was recorded from the left hand and heat was applied to the left arm. Pupillometry and gaze position was recorded with Eyelink 1000-Plus (SR-Research, Ontario, Canada).

Participants also completed questionnaires prior to the experiment, including the State-Trait Anxiety Inventory (STAI Form X (Gaudry et al., 1975), the Positive and Negative Affect Scale (Watson et al., 1988), and Behavioral Inhibition/Activation Scale (Carver and White, 1994). For the present manuscript, we focused on pain reports and brain responses evoked by painful heat. Physiological data and associations with individual difference measures may be analyzed and reported separately in future work.

### Procedure

Participants underwent an adaptive staircase pain calibration prior to the experimental task (see Figure 1A). Participants were only eligible to continue if they reported reliable increases in pain as a function of temperature (r^2^ > 0.4) and reported pain tolerance at 50 degrees or less. 4 participants were deemed ineligible based on calibration. The calibration procedure also allowed us to identify four sites that responded most similarly across temperatures and to individually calibrate temperatures associated with ratings of low pain (2 on 10-point scale), medium pain (5 on 10-point scale), and high pain (maximum tolerable pain; 8 on 10-point scale). These temperatures and skin sites were used during the main experiment, as described below. The adaptive staircase procedure has been described in depth in previous work (Atlas et al., 2010, 2012; Mischkowski et al., 2019; Dildine et al., 2020).

**Figure 1.**
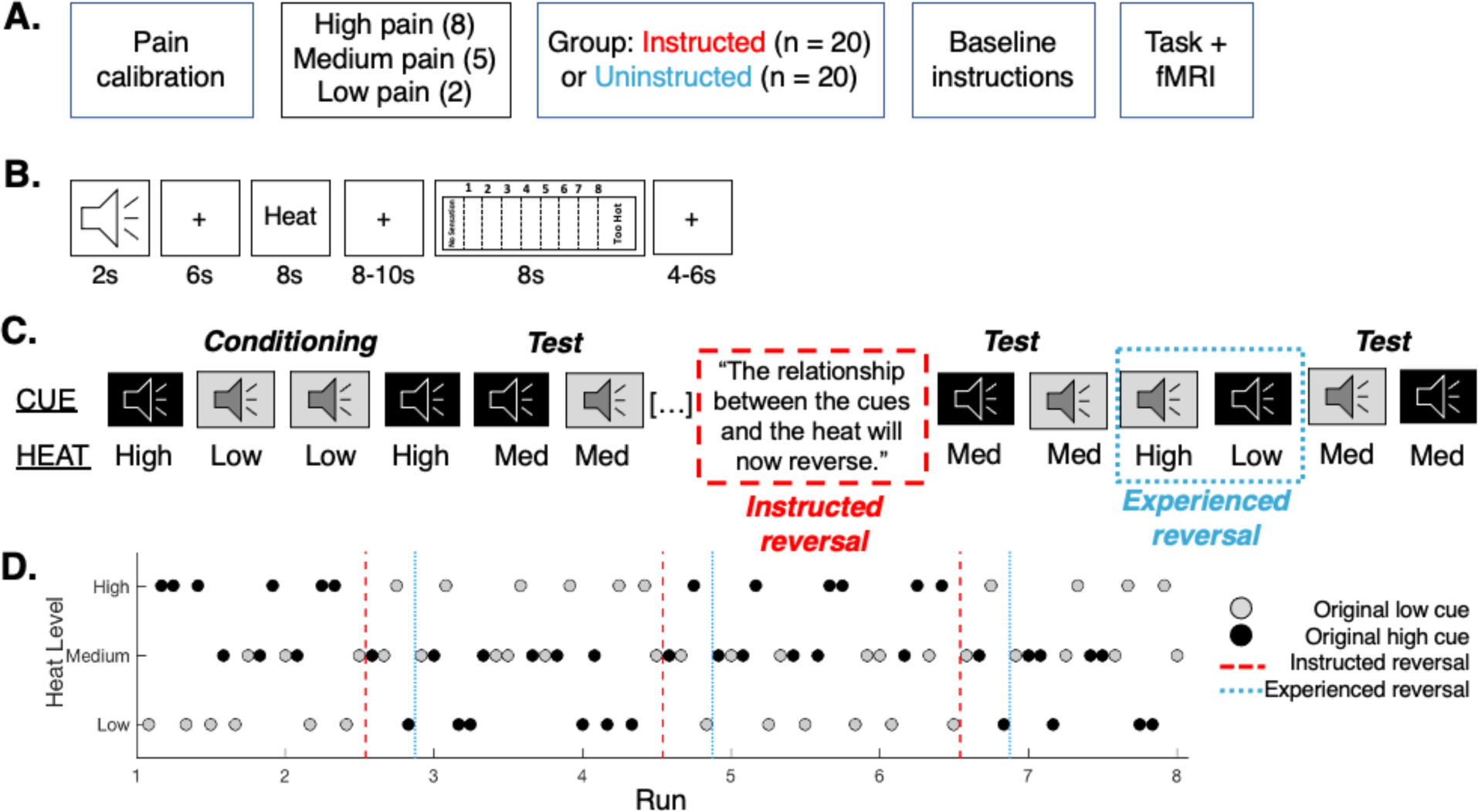
Experimental design. A) Experimental design. Participants underwent a pain calibration that identified temperatures corresponding to maximum tolerable pain (high pain; 8), pain threshold (low pain; 2), or medium pain (5). They were then positioned in the fMRI scanner and randomly assigned to group. Participants in the Instructed Group were informed about contingencies, while participants in the Uninstructed Group were told to pay attention to the associations between auditory cues and heat but were not informed about the specific cue-outcome contingencies. B) Trial structure. On each trial, a 2-second auditory cue preceded heat delivered to the participants left forearm. Participants rated perceived pain following offset using an 8-point continuous visual analogue scale. Trials were 48 seconds long. C) Instructed and experience-based reversals. Participants first underwent a conditioning phase in which Original Low Cues (gray) were followed by heat calibrated to elicit low pain (level 2) and Original High Cues (black) were followed by heat calibrated to elicit high pain (level 8). During the test phase, we delivered medium heat following each cue, which tests the effects of predictive cues on perceived pain. Following the test phase, participants in the Instructed Group were informed about reversals and we delivered medium stimuli to test the effects of instructions. We then paired high heat with the Original Low cue and low heat with the Original High cue, which should act as an experiential reversal, and again administered medium heat to test whether pain reverses upon experience. D) Example trial order. There were three reversals across the entire task. We used two trial orders that were counterbalanced across participants.

Following the pain calibration, each eligible participant was positioned in the fMRI scanner. They were then randomized (n = 20 per group) to either the Instructed Group (12 female; *M*age= 27.05 years, SDage = 6.64) or the Uninstructed Group (10 female; *M*age= 26.95 years, SDage = 5.91) and given instructions about the experiment (see Figure 1A). Both groups were told “In this task you will hear two sounds, followed by heat from the thermode. The sound will last a few seconds and will be followed by a short variable delay period, and then heat from the thermode. You do not need to respond when you hear the cue.” Instructed Group participants were then told, “It is there just to let you know what level of heat will be next.” They then heard both cues and were informed which cue would be followed by high heat and which would be followed by low heat stimulation. Uninstructed Group participants were instead told “Your job is to pay attention to the cues and try to figure out the relationship between the sounds that you hear and the heat that you feel.” They then heard both cues but were simply told that one was the first cue and the other was the second cue. We used two auditory cues (a cymbal and an accordion), which were counterbalanced across participants. Following instructions and between each block of the task, participants provided expectancy ratings in response to each cue and an experimenter moved the thermode to the next inner arm location.

During the experiment, each trial began with a 2-second auditory cue, followed by a 6- second anticipation interval, and then heat from the thermode (see Figure 1B). After an 8-10 second temporal jitter, participants provided pain ratings using an 8-point visual analogue scale. Participants viewed the scale for 3 seconds then had 5 seconds to record their ratings. There was a 4-6 second temporal jitter before the next cue was presented. The two jitters always combined to 14s within a single trial, for a total trial duration of 48s. There were 7 blocks of trials with 12 trials per block (i.e. 84 trials total). We used two trial orders (counterbalanced across participants within Group) that each included 1) a conditioning phase, 2) a test of cue-based expectancy effects, and 3) three contingency reversals (Figures 1C and 1D). During the conditioning phase, Original Low cues were followed by stimulation calibrated to elicit ratings of low pain and Original High cues were followed by stimulation calibrated to elicit high pain (see Figure 1C and 1D). The conditioning phase included three Original High cue + high heat pairings and three Original Low cue + low heat pairings. Following conditioning, each cue was paired with stimulation calibrated to elicit ratings of medium pain. This provides a test of cue-based expectancy effects, consistent with our previous work (Atlas et al., 2010, 2013; Johnston et al., 2012; Michalska et al., 2018; Abend et al., 2021). Intermittent reinforcement continued at a 50% reinforcement rate until contingencies reversed.

Halfway through runs 2, 4, and 6, the screen displayed instructions to the Instructed Group indicating that contingencies had reversed (see Figure 1C). Immediately following instructions, participants experienced at least one medium heat trial paired with each cue, which provides an immediate test of instructed reversals in the Instructed Group (Atlas et al., 2016; Atlas and Phelps, 2018; Abend et al., 2021). Following the medium heat trials, the new contingencies were reinforced (i.e. the high pain temperature was paired with the previous low pain cue and the low pain temperature was paired with the previous high pain cue), leading to an experience-based reversal. Medium trials were then delivered, and learning continued with the same reinforcement rates until the next reversal (see Figure 1D). Uninstructed Group participants experienced the same trials, but a fixation cross was displayed instead of instructions.

We used two pseudorandom trial orders with three contingency reversals, which were counterbalanced across participants. Each trial order ensured that no condition was repeated three times in a row. When we visualized responses as a function of trial order, we noticed that one trial order presented the same cue-heat condition as the first trial on 6 out of the 7 blocks (i.e. medium heat paired with the original high pain cue). Because the first trial of each block is applied to a new skin site, the novel stimulus was rated as much higher than all other trials. We therefore omitted the first trial from all analyses in the main manuscript, because this novelty response contaminated the otherwise strong reversal behavior. Furthermore, due to a programming error, Instructed Group participants in one of the two trial orders received two incompatible trials following the third instructed reversal (i.e. they received one low stimulus with the previous low cue and one high stimulus with the previous high cue). Because this experience contradicted instructions and we are interested in measuring the effects of veridical instructions on reversal learning, we only analyzed the trials prior to these instructions (i.e. two reversals instead of three) in these participants (n = 10).

Following the fMRI scan, participants rated affect associated with each cue and provided retrospective expectancy ratings to report how much pain they expected in response to each cue at the beginning and the end of the task. Results are reported in Supplementary Materials.

### BOLD FMRI data acquisition and preprocessing

BOLD fMRI data were collected on a 3T Siemens Skyra scanner at the NIH’s MRI Research Facility / Functional Magnetic Resonance Imaging Facility. After positioning the participant in the scanner bore, we collected a localizer followed by a T1-MPRAGE collected in the sagittal plane (256 slices). We collected 7 runs of multi-echo data with a 2.5s TR and 3-mm isotropic voxels (191 volumes collected anterior to posterior; flip angle = 90*°*; acquisition matrix = 70 x 0 x 0 x 64; 1^st^ echo = 11ms; 2^nd^ echo = 22ms; 3^rd^ echo 33ms).

Multi-echo data were preprocessed and combined using the “afniproc.py” program in Analysis of Functional Images (AFNI; (Cox, 1996)). We removed the first 4 TRs of functional data to reach magnetization steady state, leaving a total of 187 TRs of fMRI data per run during subsequent processing and analysis steps. Details of each preprocessing step as well as the accompanying afniproc.py commands are provided in Supplementary Methods. Weights were applied and combined using AFNI’s 3dMean program to generate “optimally combined” (OC) data, which were used for subsequent analyses. Preliminary analyses indicated that combining across the echoes with optimal combination led to better heat-related activation in pain-related regions than analyses of a single echo or using other approaches to combine multi-echo data (e.g. TE-dependent analysis (Kundu et al., 2012; Lombardo et al., 2016)). In future analyses, we may formally compare optimal combination with other approaches for echo combination and denoising. Following optimal combination, data were smoothed using a 4mm full-width half max smoothing kernel and normalized to percent signal change. Data were then analyzed using single trial estimates in Matlab, as described below.

### Statistical analysis of expectations, pain, and heat-evoked neural signature pattern expression

We used the statistical software R (R Core Team, 1996) to analyze effects of our experimental manipulations on expectancy ratings, pain reports, and heat-evoked brain signature pattern expression (see below, “Brain-based classifier analyses”). We measured effects across the entire task, as well as before the first reversal. We used 2 x 2 ANOVAs implemented through the R function anova_test from the package rstatix (Kassambara, 2021) to analyze expectancy ratings prior to the task and after conditioning, as well as post-task ratings (see Supplementary Results). All other analyses were conducted using multilevel linear mixed effects models in R.

In each analysis, we modeled within-subjects effects of Cue (original high pain cue / CS+ versus original low pain cue / CS-), Phase (original versus reversed contingencies), and Cue x Phase interactions (i.e. current high pain cue / CS+ versus current low pain cue / CS-) at the first level, and Group was modeled at the second (i.e. between-subjects) level. Consistent with our previous work on aversive reversal learning (Atlas et al., 2016; Atlas and Phelps, 2018), reversals (i.e. Phase effects) were coded relative to instructed reversals in the Instructed Group, and relative to experienced reversals in the Uninstructed Group (i.e. the first time the previous high pain cue was paired with low heat, or vice versa; see Figure 1C). Analyses across temperatures also included a first-level factor for Heat Intensity, and analyses during acquisition omitted the effect of Phase. We also included an effect of Time (linear effect of trial) in model comparisons to evaluate whether cue effects varied over time, although Bayesian model comparison indicated that including Time did not improve any models.

We evaluated linear mixed models using three statistical packages to ensure results were robust to analysis approaches. We implemented Bayesian models using *brms* (Bürkner, 2017) and used “bayesfactor_models” from the *bayestestR* package to perform model comparison and to draw inferences about acceptance and rejection of null effects (see Supplementary Methods). In most cases, Bayesian model comparisons supported maximal models (Barr et al., 2013), i.e. including all fixed factors (except Time), all interactions, and random intercepts and slopes for each subject. We complemented Bayesian statistics with frequentist statistics using the *lmer* function in the R package lme4 (Bates et al., 2015) and confirmed findings using the *nlme* package (Pinheiro et al., 2021) with autoregression (AR(1)) to account for non-independence of sequential measurements. We acknowledge findings from all approaches in our Results, using guidelines for Bayesian modeling from Makowski et al. (Makowski et al., 2019b, 2019a), and report details of model specifications and comparisons in Supplementary Results. For additional details on Bayesian models and model comparison, please see Supplementary Methods.

### Computational modeling of pain reversal learning and modulation by instructions

We applied a computational model of instructed reversal learning (Atlas et al., 2016; Atlas and Phelps, 2018) to predict pain reports on medium heat trials. In brief, the model includes two free parameters: an instructed reversal parameter (⍴), which determines whether expected value of cues are exchanged in response to reversal instructions, and a learning rate (ɑ), which determines how swiftly value updates in response to prediction errors. See Atlas et al., (Atlas et al., 2016) and Supplementary Methods for additional details. We compared our instructed reversal learning model with an uninstructed model (i.e. a standard Rescorla-Wagner model), a model that included an additional free parameter for initial expected value, and an adaptive learning model modified to reverse expected value upon instruction (Li et al., 2011b; Atlas et al., 2019). Models were fit to each individual’s pain ratings on medium heat trials, since the stimulus temperature was constant and therefore the only factor likely to guide cue-based variation in pain would presumably be dynamic expected value based on learned and/or instructed value. Models were analyzed at the level of the individual, which provides estimates that can be tested using standard statistics, and through an iterative jack-knife procedure, which is less sensitive to noise based on individual estimates (Wu, 1986; Miller et al., 1998; Atlas and Phelps, 2018). Group analyses and model comparisons were conducted using individual estimates. Goodness-of-fit was evaluated based on Akaike’s Information Criterion (Akaike, 1974), which evaluates model fit with a penalty for additional parameters and is appropriate for nested models, and models were compared based on Bayesian Model Selection, as implemented in SPM’s SPM_bms.m (Stephan et al., 2009). As reported in Supplementary Results, our basic instructed learning model (Atlas et al., 2016; Atlas and Phelps, 2018) with a constant learning rate and fixed initial expected values provided the best fit for pain reports. We used t-tests to assess whether each parameter (ɑ and ⍴) varied as a function of group. The mean parameters based on jack-knife estimates were used to generate regressors for fMRI analyses (see “Neural correlates of expected value”, below and Supplementary Results).

### FMRI Analyses

#### Single trial analyses

Following preprocessing in AFNI, we used single trial analyses to estimate heat-evoked responses on a trial-by-trial basis and avoid assumptions about the fixed shape of the hemodynamic response. We used flexible basis functions optimized to capture heat- evoked BOLD responses, consistent with previous work on heat-evoked fMRI (Atlas et al., 2010, 2012, 2014; Wager et al., 2013; Woo et al., 2017). We applied principal components analysis-based spike detection (scn_session_spike_id.m, available at https://canlab.github.io/) to identify potential spikes and noise in the data which were modeled as nuisance covariates, along with the movement parameters from AFNI’s preprocessing pipeline. We used the function single_trial_analysis.m (https://canlab.github.io/) to generate trial-by-trial estimates of height, width, delay, and area-under-the-curve (AUC) for each heat period. Trials in which subjects failed to respond were omitted from analyses. We focused on AUC estimates that were smoothed with a 4mm Gaussian kernel in subsequent analyses, consistent with our previous work. We computed variance inflation factors (VIFs) using the single_trial_weights_vifthresh.m function to identify bad trials, i.e. those who coincided with spikes or motion and were therefore not reliable estimates. We excluded any trials with VIFs > 2 from subsequent analyses (*M* = 3.43, *SD =* 2.30), consistent with previous work (Atlas et al., 2010, 2012, 2014). Trial estimates were passed into voxel-wise second level analyses across trials and across participants using the general linear model (fit_gls_brain.m; https://canlab.github.io/) and robust regression (robfit.m; https://canlab.github.io/) to examine neural correlates of associative learning (see below, “*Neural correlates of expected value*”). Trial-level estimates were also employed in multilevel mediation analyses (see below, “*Multilevel mediation analyses”*).

#### Multilevel mediation analyses

We used multilevel mediation to examine whether brain activity mediated the effect of predictive cues on subjective pain on medium heat trials.

Mediation was implemented by the Matlab function *mediation.m* (https://canlab.github.io/). Cue was included as the input variable (i.e. X), Pain was included as the output variable (i.e. Y), and we searched for potential mediators. Voxelwise mediation, or mediation effect parametric mapping (Wager et al., 2009; Atlas et al., 2010) yields interpretable maps for each of the effects: Path *a* denotes the effect of the input variable on the potential mediator, thereby representing cue effects on brain responses to medium heat. Path *b* measures the association between the mediator and outcome, controlling for the input variable. Here, this represents brain regions that predict pain, controlling for cue type. Finally, the mediation effect (a*b) identifies regions whose activity contributes to variance in the effect of the independent variable on the dependent variable (Path *c*). In multilevel mediation, the difference between the total effect (Path *c*: the effect of cues on subjective pain) and the direct effect (Path c’: the effect of cues on subjective pain when controlling for the mediator) is equivalent to the sum of the product of Path *a* and Path *b* coefficients and their covariance (Shrout and Bolger, 2002; Kenny et al., 2003).

We ran two types of mediation analyses on medium heat trials: a) voxel-wise mediation analyses, which search for brain regions that mediate the effects of predictive cues on pain; b) statistical tests of whether brain responses within ROIs formally mediated cue effects on subjective pain. We evaluated mediation analyses irrespective of Group, and with Group as moderator. We used bootstrapping to estimate the significance of the mediation effect (Shrout and Bolger, 2002; Kenny et al., 2003) in analyses irrespective of Group, and used ordinary least squares to estimate moderated mediation when Group was included in the model. We omitted the first trial of each run from analyses to be consistent with behavioral results, and focus on mediation of current cue contingencies (i.e. Cue x Phase interactions), which identify responses that reverse as contingencies change. To isolate brain responses that maintain initial contingencies regardless of reversals, we conducted a second mediation analysis to examine effects of original cue contingencies, controlling for current contingencies.

#### Neural correlates of expected value

Whereas our mediation analyses tested effects of Phase, i.e. immediate changes in response to instruction or contingency reversal, we used quantitative models to test whether expected value dynamically shapes responses to noxious stimulation. We used parameters from the best-fitting models for each group, based on jack-knife estimation, to generate the timecourse of expected value for each subject based on their sequence of trials. We chose to use the mean of the group-level estimates to avoid noise that might come from individual-level model fits. We examined the neural correlates of expected value on medium heat trials only, which avoids confounds due to temperature. Noxious stimulation might also be accompanied by prediction errors; for example, if an individual expects high pain and receives medium heat, this should generate an appetitive PE if the deviation is noticed. However, expected value and prediction error are inversely correlated in the standard RL model we used.

We therefore only modeled expected value in our analyses. We focused on how expected value influenced responses to medium intensity heat, rather than responses to cues themselves, as we were most interested in how pain-related responses are influenced by learned expectations, and we did not optimize the anticipatory period to jointly estimate cue-evoked responses and responses to heat. We report three group-level analyses: 1) Analysis across all participants testing for differences by group; 2) Analyses within each group to isolate effects of instructed learning (Instructed Group) or feedback-driven learning (Uninstructed Group); 3) Comparisons of instructed and feedback-driven learning within the Instructed Group. Individual results were computed using the matlab function fit_gls_brain (https://canlab.github.io/) and group results were computed using robust regression (Wager et al., 2005) using the function robfit.m (https://canlab.github.io/).

#### Brain-based classifier analyses

To isolate effects on pain-related regions, we employed two recently developed brain-based classifiers that have been shown to be sensitive and specific to acute pain, the Neurologic Pain Signature (NPS; (Wager et al., 2013)) and the Stimulus Intensity Independent Pain Signature (SIIPS; (Woo et al., 2017)). Each consists of a pattern of weights across the brain, which can be combined with a brain activation map (e.g. a coefficient for a condition, a contrast, or a trial estimate) by computing the dot product of weights and the result map. The resulting values have been shown to accurately predict subjective pain, whether a stimulus is painful or not, and which of two conditions is more painful (Wager et al., 2013; Woo et al., 2017). Here, we computed the dot-product of each signature with trial-level images, which allowed us to use the brain response as an outcome in our multilevel mediation analyses. To validate the use of the brain-based classifiers, we tested the association between subjective pain and brain activity, both across temperatures and within medium heat trials (see Supplementary Results). We used the unthresholded NPS pattern for all analyses, and the function apply_mask.m (https://canlab.github.io/) to compute dot products. Resulting pattern expression values were analyzed using the same multilevel approach outlined above for behavioral outcomes. We also computed pattern expression across beta coefficients to evaluate associations between expected value and NPS and SIIPS expression.

#### Pain modulatory regions of interest

While the NPS and SIIPS capture activation related to acute pain, other modulatory regions are involved in regulating pain that may not be captured in the patterns, and previous work indicates that some forms of pain modulation, including placebo analgesia, may not elicit reliable changes in the NPS (Zunhammer et al., 2018). We therefore also tested for cue-based modulation of brain regions that have been previously implicated in studies of expectancy-based pain modulation by applying an *a priori* mask generated from our previous meta-analysis of fMRI studies of placebo analgesia and expectancy- based modulation (Atlas and Wager, 2014). We included regions that showed either expectancy- related increases or decreases in activation within the mask. Supplemental Figure S1A depicts the mask, which includes regions that show increased activation with expected pain relief (i.e. activation inversely related to subjective pain) such as the DLPFC, rACC, PAG, and VMPFC, and regions that show reduced activation with expected pain relief, including the insula, thalamus, cingulate, and secondary somatosensory cortex. We report results FDR-corrected within this mask to evaluate responses within pain modulatory regions.

#### Value-processing regions of interest

In addition to pain-related classifiers and pain modulatory networks, we were also interested in testing effects of predictive cues on brain regions involved in value-based learning. To this end, we examined responses within 5 *a priori* regions of interest (see Supplemental Figure S1B): the bilateral striatum, bilateral amygdala, and the ventromedial prefrontal cortex (VMPFC). We used the same ROI masks that were applied in our prior work on instructed reversal learning (Atlas et al., 2016). While the amygdala and striatum masks were defined based on Atlases in MNI space (amygdala ROI available at https://canlab.github.io/; striatum ROI based on combining putamen and caudate masks from the Automated Anatomical Labeling atlas for SPM8 (http://www.gin.cnrs.fr/AAL; (Tzourio-Mazoyer et al., 2002)), the VMPFC ROI was functionally defined in our previous work by analyzing deactivation in response to shock. We used the same ROI mask here since analyses as a function of heat intensity elicited significant decreases in this region. We averaged trial-level AUC estimates across each ROI to conduct mediation analyses and averaged across beta coefficients and contrast maps to analyze ROI-wise associations with expected value. Results are reported in Table 4.

#### Whole brain exploratory analyses

In addition to the analyses in *a priori* networks and regions of interest involved in pain, placebo, and value-based processing, we also conducted exploratory voxel-wise whole brain analyses. We report whole brain results at FDR-corrected p < .05 in the main manuscript, and present exploratory uncorrected results at p < .001 in Supplementary Results for completeness and for use in future meta-analyses. Anatomical labels were identified using the SPM Anatomy Toolbox (Eickhoff et al., 2005).

## Results

### Heat intensity effects on pain are similar across groups

Prior to the fMRI experiment, all participants underwent an adaptive pain calibration procedure (Atlas et al., 2010; Mischkowski et al., 2019; Dildine et al., 2020) to identify each participant’s pain threshold, tolerance, and the reliability of the temperature-pain association (i.e. r^2^). Consistent with our IRB protocol, four participants were dismissed prior to the fMRI portion of the experiment due to low reliability (n = 3) or pain tolerance above 50℃ (n = 1). For each participant who continued to the fMRI phase, we used linear regression to identify temperatures associated with ratings of low pain (*M* = 42.04℃, *SE* = 0.43), medium pain (*M* = 44.71℃, *SE* = 0.37), and high pain (*M* = 47.30℃, *SE* = 0.30). There were no differences between groups in the reliability of the association between temperature and pain, as measured by r^2^ (*M* = 0.803, *SE* = 0.022; p > 0.2), or in temperatures applied during the task (all p’s > 0.1).

We next examined pain as a function of heat intensity (i.e. temperature level: low, medium, or high) during the fMRI experiment (see Supplementary Figure S2). Bayesian model comparison indicated that the best model included fixed effects of Heat Intensity, Cue, Phase, and Group and all possible interactions, along with random intercepts and slopes for all factors (see Supplemental Results). All models revealed significant effects of Heat Intensity, Cue, Phase, Cue x Phase, and Heat Intensity x Cue x Phase interactions across participants (see Table 1). We also observed a significant Group x Cue x Phase interaction and a significant Group x Heat Intensity x Cue x Phase interaction, which were likely to be driven by the critical medium heat trials, as reported below. Bayesian posterior estimates indicated that the effects of Heat Intensity, Cue x Phase interactions, and Heat Intensity x Cue x Phase interactions were practically significant with enough evidence to reject the null (<1% in ROPE), while the main effect Phase supported the null (i.e. no effect of Phase; 99.8% in ROPE), despite being statistically significant. All other effects were of undecided significance (i.e. not enough evidence to accept or reject the null); complete results are reported in Table 1. We observed similar results when we restricted analyses to pain ratings from the 35 participants with useable fMRI data; see Supplementary Table S1.

**Table 1.**
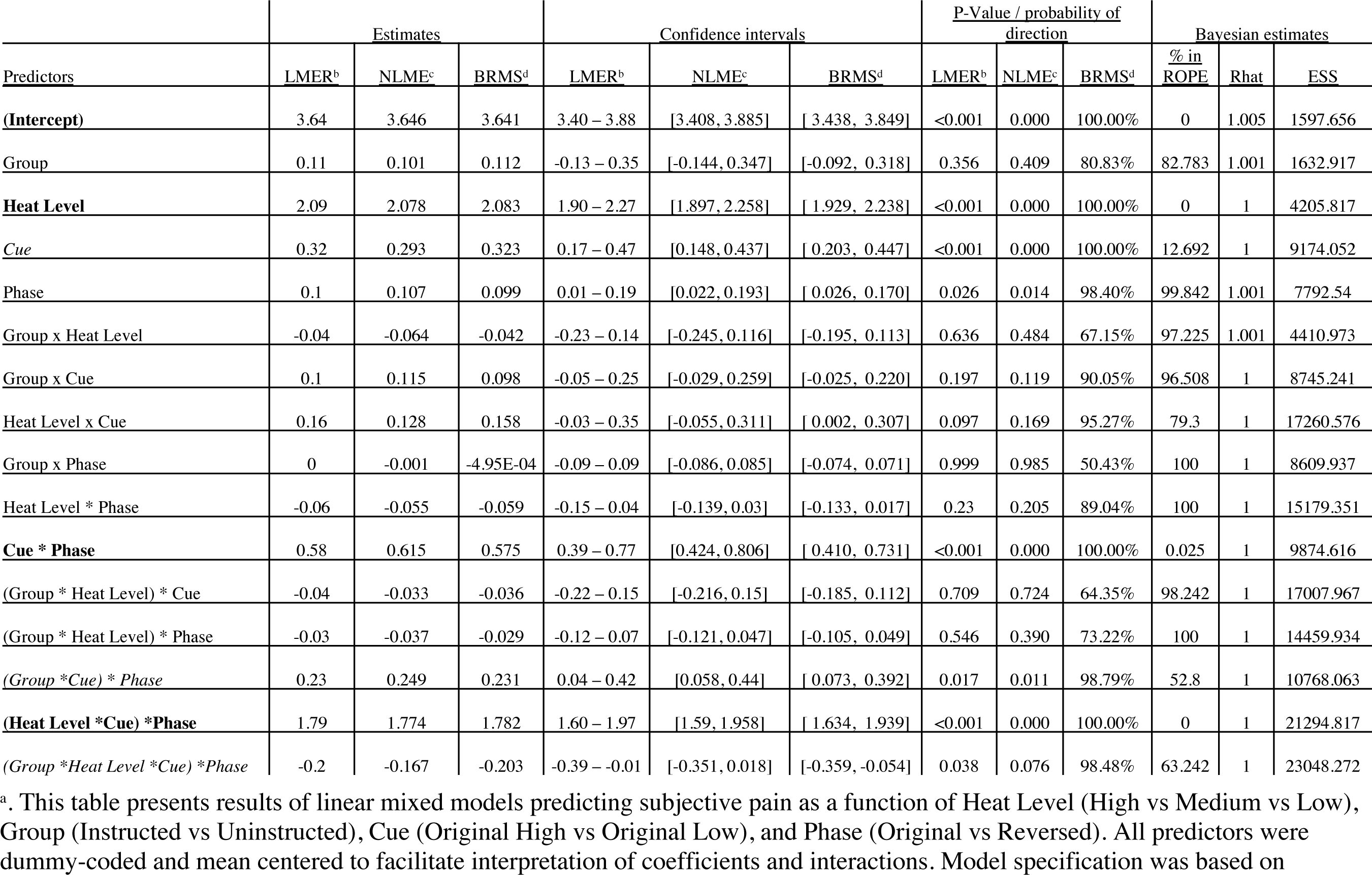
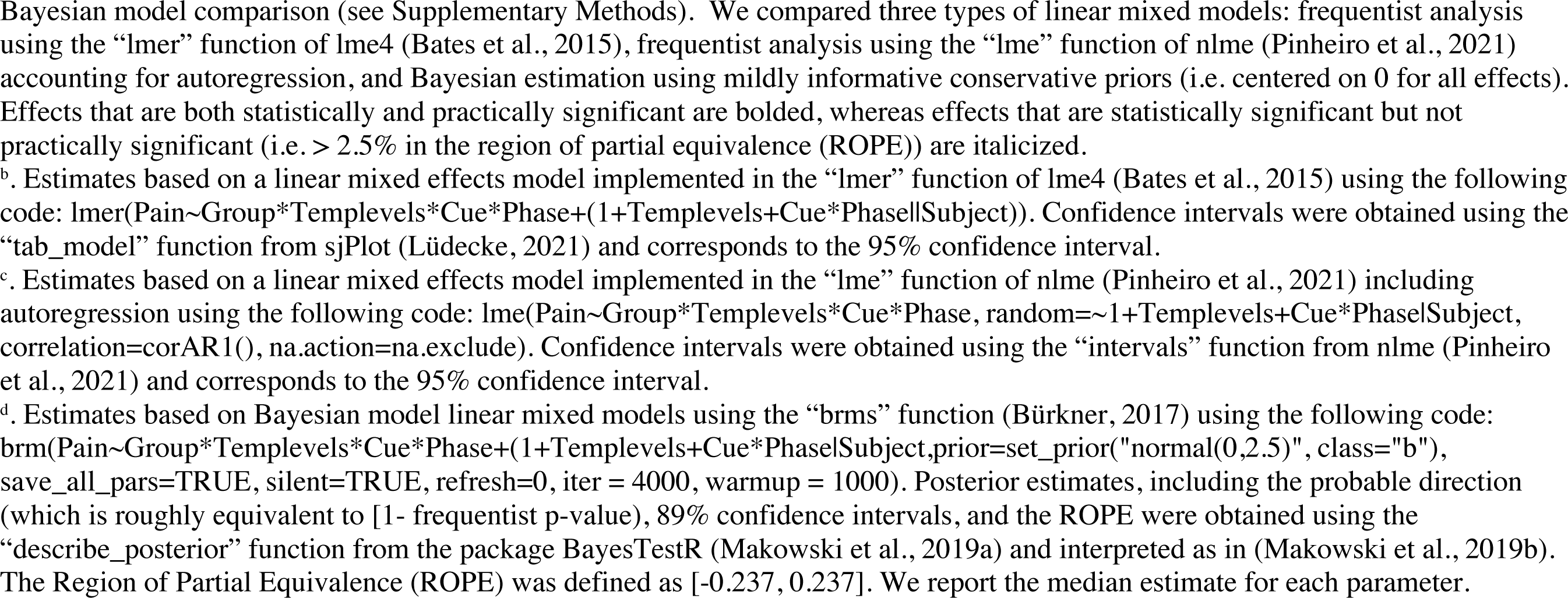
Heat intensity effects on pain across all participants (n = 40).^a^

| Predictors | Estimates |  |  | Confidence intervals |  |  | P-Value / probability of direction |  |  | Bayesian estimates |  |  |
| --- | --- | --- | --- | --- | --- | --- | --- | --- | --- | --- | --- | --- |
|  | LMER<sup>b</sup> | NLME<sup>c</sup> | BRMS<sup>d</sup> | LMER<sup>b</sup> | NLME<sup>c</sup> | BRMS<sup>d</sup> | LMER<sup>b</sup> | NLME<sup>c</sup> | BRMS<sup>d</sup> | % in ROPE | Rhat | ESS |
| **(Intercept)** | 3.64 | 3.646 | 3.641 | 3.40 – 3.88 | [3.408, 3.885] | [ 3.438, 3.849] | <0.001 | 0.000 | 100.00% | 0 | 1.005 | 1597.656 |
| Group | 0.11 | 0.101 | 0.112 | -0.13 – 0.35 | [-0.144, 0.347] | [-0.092, 0.318] | 0.356 | 0.409 | 80.83% | 82.783 | 1.001 | 1632.917 |
| **Heat Level** | 2.09 | 2.078 | 2.083 | 1.90 – 2.27 | [1.897, 2.258] | [ 1.929, 2.238] | <0.001 | 0.000 | 100.00% | 0 | 1 | 4205.817 |
| **Cue** | 0.32 | 0.293 | 0.323 | 0.17 – 0.47 | [0.148, 0.437] | [ 0.203, 0.447] | <0.001 | 0.000 | 100.00% | 12.692 | 1 | 9174.052 |
| Phase | 0.1 | 0.107 | 0.099 | 0.01 – 0.19 | [0.022, 0.193] | [ 0.026, 0.170] | 0.026 | 0.014 | 98.40% | 99.842 | 1.001 | 7792.54 |
| Group x Heat Level | -0.04 | -0.064 | -0.042 | -0.23 – 0.14 | [-0.245, 0.116] | [-0.195, 0.113] | 0.636 | 0.484 | 67.15% | 97.225 | 1.001 | 4410.973 |
| Group x Cue | 0.1 | 0.115 | 0.098 | -0.05 – 0.25 | [-0.029, 0.259] | [-0.025, 0.220] | 0.197 | 0.119 | 90.05% | 96.508 | 1 | 8745.241 |
| Heat Level x Cue | 0.16 | 0.128 | 0.158 | -0.03 – 0.35 | [-0.055, 0.311] | [ 0.002, 0.307] | 0.097 | 0.169 | 95.27% | 79.3 | 1 | 17260.576 |
| Group x Phase | 0 | -0.001 | -4.95E-04 | -0.09 – 0.09 | [-0.086, 0.085] | [-0.074, 0.071] | 0.999 | 0.985 | 50.43% | 100 | 1 | 8609.937 |
| Heat Level * Phase | -0.06 | -0.055 | -0.059 | -0.15 – 0.04 | [-0.139, 0.03] | [-0.133, 0.017] | 0.23 | 0.205 | 89.04% | 100 | 1 | 15179.351 |
| **Cue \* Phase** | 0.58 | 0.615 | 0.575 | 0.39 – 0.77 | [0.424, 0.806] | [ 0.410, 0.731] | <0.001 | 0.000 | 100.00% | 0.025 | 1 | 9874.616 |
| (Group * Heat Level) * Cue | -0.04 | -0.033 | -0.036 | -0.22 – 0.15 | [-0.216, 0.15] | [-0.185, 0.112] | 0.709 | 0.724 | 64.35% | 98.242 | 1 | 17007.967 |
| (Group * Heat Level) * Phase | -0.03 | -0.037 | -0.029 | -0.12 – 0.07 | [-0.121, 0.047] | [-0.105, 0.049] | 0.546 | 0.390 | 73.22% | 100 | 1 | 14459.934 |
| (Group * Cue) * Phase | 0.23 | 0.249 | 0.231 | 0.04 – 0.42 | [0.058, 0.44] | [ 0.073, 0.392] | 0.017 | 0.011 | 98.79% | 52.8 | 1 | 10768.063 |
| **(Heat Level \* Cue) \* Phase** | 1.79 | 1.774 | 1.782 | 1.60 – 1.97 | [1.59, 1.958] | [ 1.634, 1.939] | <0.001 | 0.000 | 100.00% | 0 | 1 | 21294.817 |
| (Group * Heat Level * Cue) * Phase | -0.2 | -0.167 | -0.203 | -0.39 – -0.01 | [-0.351, 0.018] | [-0.359, -0.054] | 0.038 | 0.076 | 98.48% | 63.242 | 1 | 23048.272 |

<sup>a</sup>. This table presents results of linear mixed models predicting subjective pain as a function of Heat Level (High vs Medium vs Low), Group (Instructed vs Uninstructed), Cue (Original High vs Original Low), and Phase (Original vs Reversed). All predictors were dummy-coded and mean centered to facilitate interpretation of coefficients and interactions. Model specification was based on
Bayesian model comparison (see Supplementary Methods). We compared three types of linear mixed models: frequentist analysis using the “lmer” function of lme4 (Bates et al., 2015), frequentist analysis using the “lme” function of nlme (Pinheiro et al., 2021) accounting for autoregression, and Bayesian estimation using mildly informative conservative priors (i.e. centered on 0 for all effects). Effects that are both statistically and practically significant are bolded, whereas effects that are statistically significant but not practically significant (i.e. > 2.5% in the region of partial equivalence (ROPE)) are italicized.
<sup>b</sup>. Estimates based on a linear mixed effects model implemented in the “lmer” function of lme4 (Bates et al., 2015) using the following code: `lmer(Pain~Group*Templevels*Cue*Phase+(1+Templevels+Cue*Phase|Subject))`. Confidence intervals were obtained using the “tab\_model” function from sjPlot (Lüdtke, 2021) and corresponds to the 95% confidence interval.
<sup>c</sup>. Estimates based on a linear mixed effects model implemented in the “lme” function of nlme (Pinheiro et al., 2021) including autoregression using the following code: `lme(Pain~Group*Templevels*Cue*Phase, random=~1+Templevels+Cue*Phase|Subject, correlation=corAR1(), na.action=na.exclude)`. Confidence intervals were obtained using the “intervals” function from nlme (Pinheiro et al., 2021) and corresponds to the 95% confidence interval.
<sup>d</sup>. Estimates based on Bayesian model linear mixed models using the “brms” function (Bürkner, 2017) using the following code: `brm(Pain~Group*Templevels*Cue*Phase+(1+Templevels+Cue*Phase|Subject,prior=set_prior("normal(0,2.5)", class="b"), save_all_pars=TRUE, silent=TRUE, refresh=0, iter = 4000, warmup = 1000)`. Posterior estimates, including the probable direction (which is roughly equivalent to [1- frequentist p-value), 89% confidence intervals, and the ROPE were obtained using the “describe\_posterior” function from the package BayesTestR (Makowski et al., 2019a) and interpreted as in (Makowski et al., 2019b). The Region of Partial Equivalence (ROPE) was defined as [-0.237, 0.237]. We report the median estimate for each parameter.

### Predictive cues modulate expectations and pain whether learned through instruction or experience

Analyses across all trials indicated potential influences of predictive cues and cue-based reversals on pain, as indicated by the Cue x Phase and Heat Intensity x Cue x Phase interactions. To measure cue-based expectancy effects more directly, we measured cue effects on 1) expectancy ratings and 2) pain reports on medium heat trials, which were crossed with predictive cues. We first examined expectations as a function of Cue prior to conditioning, i.e. immediately after instruction. Consistent with our manipulation, there was a significant Group x Cue interaction on expectancy at baseline (F(1,38) = 8.959, p = .005), driven by significant differences in the Instructed Group (p = .0027) but not the Uninstructed Group (p > 0.3), as shown in Figure 2A. There were no main effects of Group or Cue prior to conditioning (all p’s > 0.1). Following the first acquisition block, we collected a second set of expectancy ratings. We again observed a significant Group x Cue interaction (F(1,38) = 7.102, p = .011) as well as a main effect of Cue (F(1,38) = 31.195, p < .001). Post-hoc comparisons indicated that both groups reported higher expectancy with the high pain cue (see Figure 2A), but that differences were larger in the Instructed Group (p < .001), relative to the Uninstructed Group (p = .003). Thus, instructions and learning both modulated cue-based expectations about pain.

**Figure 2.**
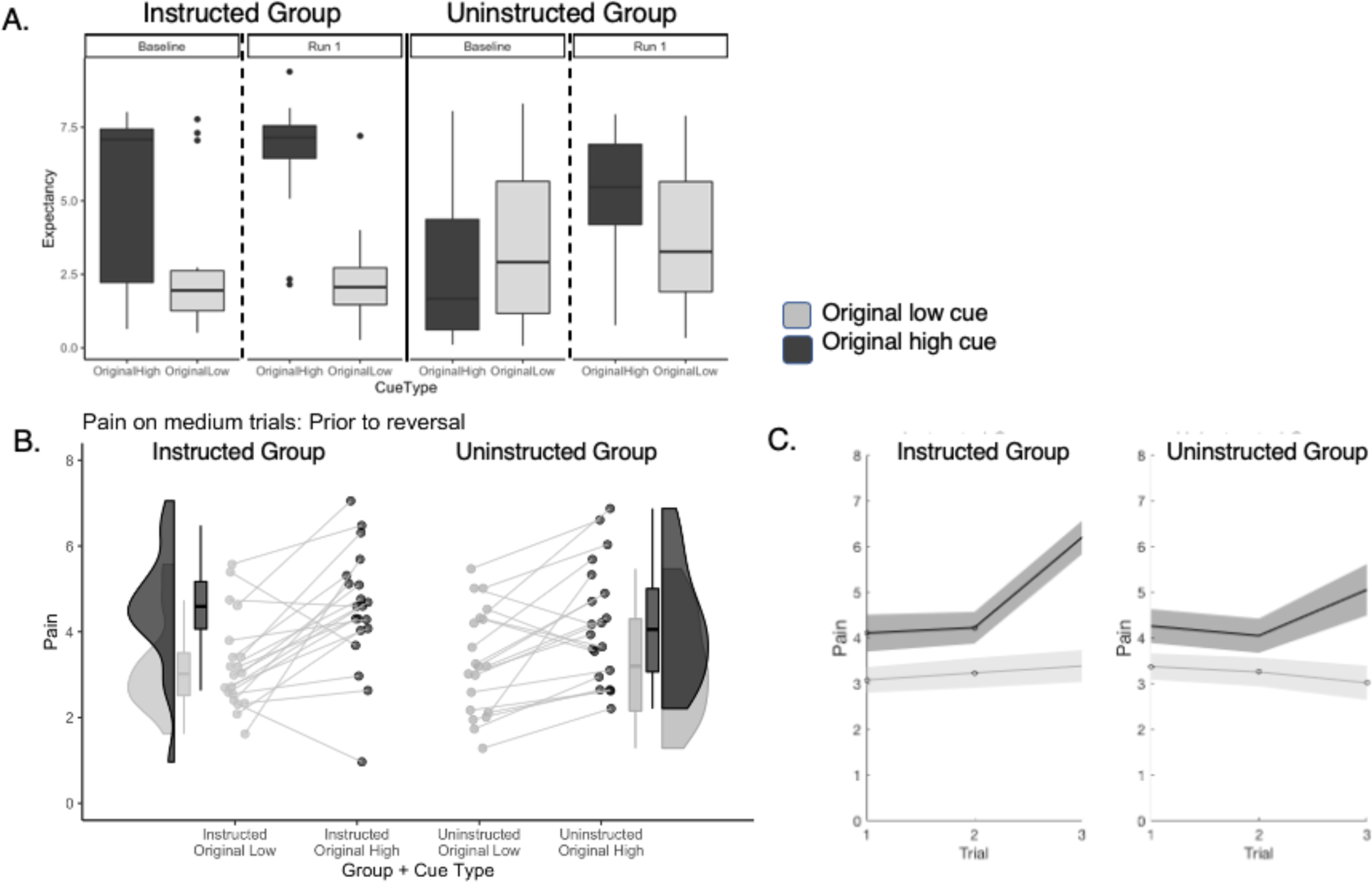
Effects of instructions and learning on expected pain and pain ratings prior to reversal. A) Expectancy ratings prior to reversal. Participants in the Instructed Group (Top Left) expected higher pain in response to the Original High Cue relative to the Original Low Cue at baseline (left) and differences in expectations grew larger following conditioning and the first test phase (right). Participants in the Uninstructed Group did not report differences prior to the task (left), consistent with the fact that they were not instructed about specific cue-outcome contingencies. Following conditioning and the first test phase, Uninstructed Group participants expected higher pain in response to the Original High Cue, relative to the Original Low Cue. Cue-based differences in expectancy ratings were larger in the Instructed Group. B) Predictive cue effects on pain prior to reversal. We measured the effects of predictive cues on perceived pain prior to the first reversal. Both groups reported higher pain when medium heat was preceded by the high pain cue (black) relative to the low pain cue (gray) and this effect was present in nearly all participants. C) Cue effects increase over time. Both groups show larger cue-based differences in perceived pain on medium heat trials as a function of experience prior to the first reversal, but effects of time were larger in the Instructed Group. Data were visualized using the R toolboxes ggplot2 (Wickham, 2016) and Raincloud plots (Allen et al., 2021).

We next asked whether cue-based expectations in turn modulate subjective pain on medium heat trials. We first measured effects during the acquisition phase, i.e. prior to the first reversal, and asked whether effects vary based on whether learning is paired with verbal instruction. Bayesian model comparison indicated that the best model included fixed effects of Group, Cue, and Trial, with random intercepts and random slopes for Cue and Trial (see Supplemental Results). Consistent with other studies of expectancy-based pain modulation (Atlas et al., 2010; Wiech et al., 2014; Reicherts et al., 2016; Fazeli and Büchel, 2018; Michalska et al., 2018; Abend et al., 2021), all models indicated that participants reported higher pain when medium heat was preceded by high pain cues than low pain cues (main effect of Cue: see Figure 2B and Table 2). Bayesian modeling indicated that this effect had a 100% probability of being positive (Median = 1.261, 89% CI [0.82, 1.338]), and can be considered practically significant (0% in ROPE). There was a significant Group x Cue interaction (see Table 2) which was of undecided significance (8% in ROPE). Importantly, post-hoc analyses within groups indicated that both groups reported practically significant effects of Cue on pain prior to the first reversal (see Figure 2B and Table 2), although effects were larger in the Instructed Group. We also observed a statistically significant Group x Cue x Trial interaction (see Table 2), although there was not enough evidence to accept the null of no difference, as 35.45% of the posterior estimate was within the ROPE. Post-hoc analyses within groups indicated that Cue effects increased over time in the Instructed Group (see Figure 2C and Table 2), as did pain reports overall (although neither of these effects were practically significant), whereas there were no interactions with time in Uninstructed Group participants. Together, these results indicate that instructions and learning both shape pain prior to reversal, that effects are somewhat larger in Instructed Group participants, and that the dynamics of expectancy effects on pain may differ as a function of whether individuals learn from experience or instruction. For complete results, please see Table 2.

**Table 2.**
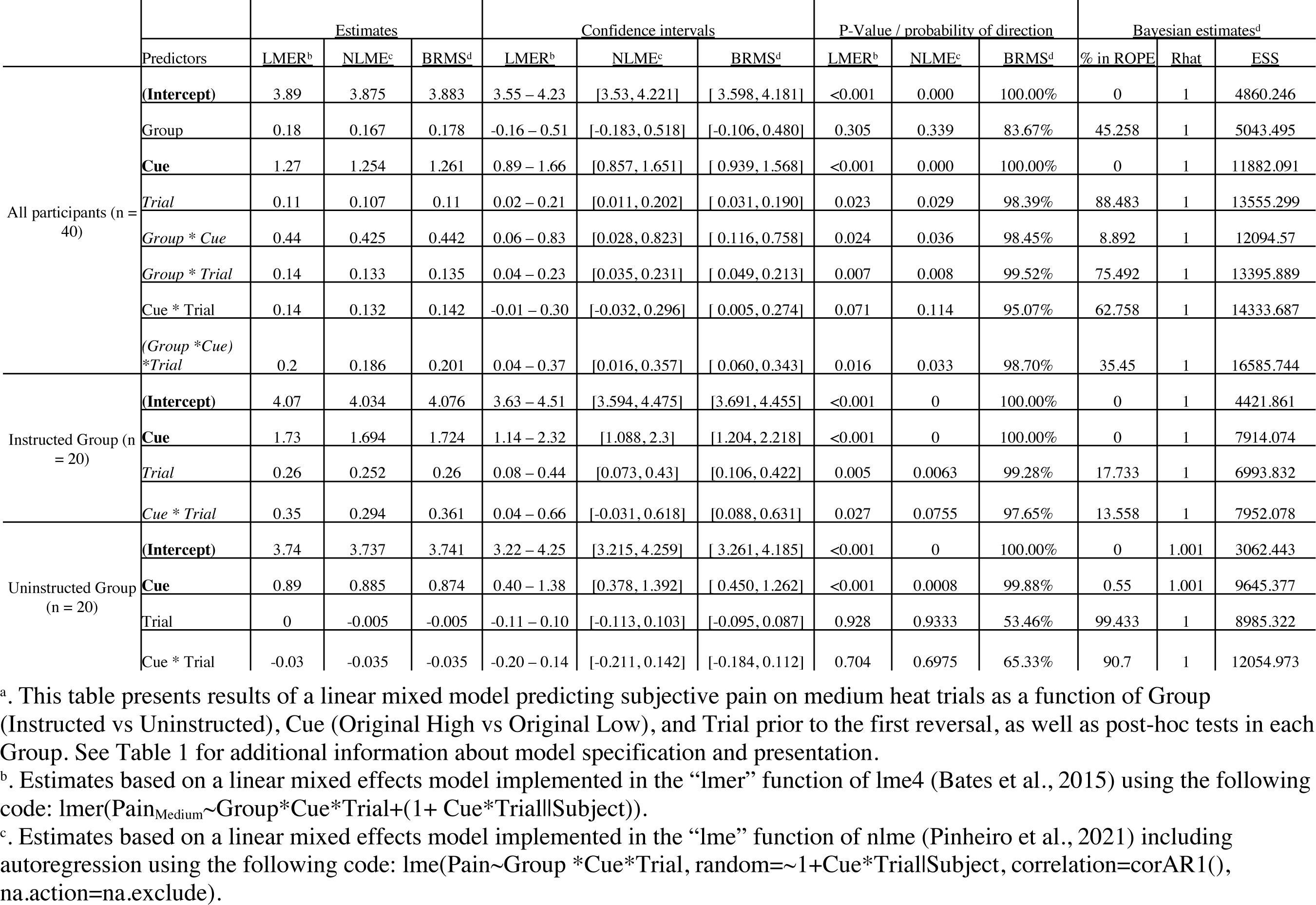
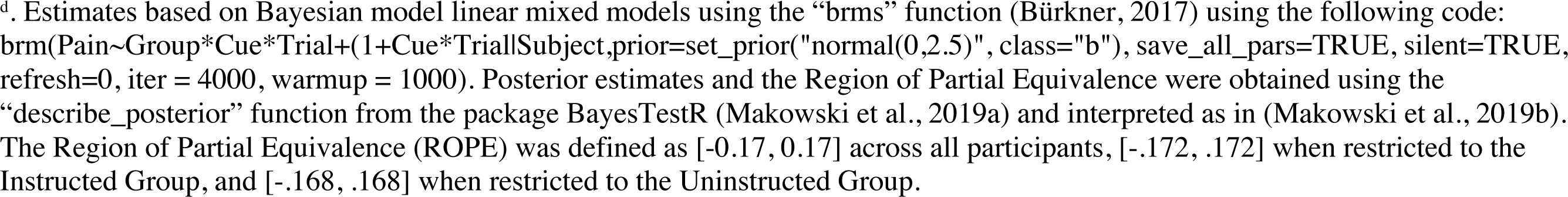
Multilevel model evaluating effects of Group, Cue, and Trial on medium heat pain prior to reversal.^a^

|  | Predictors | Estimates |  |  | Confidence intervals |  |  | P-Value / probability of direction |  |  | Bayesian estimates<sup>d</sup> |  |  |
| --- | --- | --- | --- | --- | --- | --- | --- | --- | --- | --- | --- | --- | --- |
|  |  | LMER<sup>b</sup> | NLME<sup>c</sup> | BRMS<sup>d</sup> | LMER<sup>b</sup> | NLME<sup>c</sup> | BRMS<sup>d</sup> | LMER<sup>b</sup> | NLME<sup>c</sup> | BRMS<sup>d</sup> | % in ROPE | Rhat | ESS |
| All participants (n = 40) | **(Intercept)** | 3.89 | 3.875 | 3.883 | 3.55 – 4.23 | [3.53, 4.221] | [3.598, 4.181] | <0.001 | 0.000 | 100.00% | 0 | 1 | 4860.246 |
|  | Group | 0.18 | 0.167 | 0.178 | -0.16 – 0.51 | [-0.183, 0.518] | [-0.106, 0.480] | 0.305 | 0.339 | 83.67% | 45.258 | 1 | 5043.495 |
|  | **Cue** | 1.27 | 1.254 | 1.261 | 0.89 – 1.66 | [0.857, 1.651] | [0.939, 1.568] | <0.001 | 0.000 | 100.00% | 0 | 1 | 11882.091 |
|  | *Trial* | 0.11 | 0.107 | 0.11 | 0.02 – 0.21 | [0.011, 0.202] | [0.031, 0.190] | 0.023 | 0.029 | 98.39% | 88.483 | 1 | 13555.299 |
|  | *Group \* Cue* | 0.44 | 0.425 | 0.442 | 0.06 – 0.83 | [0.028, 0.823] | [0.116, 0.758] | 0.024 | 0.036 | 98.45% | 8.892 | 1 | 12094.57 |
|  | *Group \* Trial* | 0.14 | 0.133 | 0.135 | 0.04 – 0.23 | [0.035, 0.231] | [0.049, 0.213] | 0.007 | 0.008 | 99.52% | 75.492 | 1 | 13395.889 |
|  | *Cue \* Trial* | 0.14 | 0.132 | 0.142 | -0.01 – 0.30 | [-0.032, 0.296] | [0.005, 0.274] | 0.071 | 0.114 | 95.07% | 62.758 | 1 | 14333.687 |
|  | *(Group \* Cue) \* Trial* | 0.2 | 0.186 | 0.201 | 0.04 – 0.37 | [0.016, 0.357] | [0.060, 0.343] | 0.016 | 0.033 | 98.70% | 35.45 | 1 | 16585.744 |
| Instructed Group (n = 20) | **(Intercept)** | 4.07 | 4.034 | 4.076 | 3.63 – 4.51 | [3.594, 4.475] | [3.691, 4.455] | <0.001 | 0 | 100.00% | 0 | 1 | 4421.861 |
|  | **Cue** | 1.73 | 1.694 | 1.724 | 1.14 – 2.32 | [1.088, 2.3] | [1.204, 2.218] | <0.001 | 0 | 100.00% | 0 | 1 | 7914.074 |
|  | *Trial* | 0.26 | 0.252 | 0.26 | 0.08 – 0.44 | [0.073, 0.43] | [0.106, 0.422] | 0.005 | 0.0063 | 99.28% | 17.733 | 1 | 6993.832 |
|  | *Cue \* Trial* | 0.35 | 0.294 | 0.361 | 0.04 – 0.66 | [-0.031, 0.618] | [0.088, 0.631] | 0.027 | 0.0755 | 97.65% | 13.558 | 1 | 7952.078 |
| Uninstructed Group (n = 20) | **(Intercept)** | 3.74 | 3.737 | 3.741 | 3.22 – 4.25 | [3.215, 4.259] | [3.261, 4.185] | <0.001 | 0 | 100.00% | 0 | 1.001 | 3062.443 |
|  | **Cue** | 0.89 | 0.885 | 0.874 | 0.40 – 1.38 | [0.378, 1.392] | [0.450, 1.262] | <0.001 | 0.0008 | 99.88% | 0.55 | 1.001 | 9645.377 |
|  | *Trial* | 0 | -0.005 | -0.005 | -0.11 – 0.10 | [-0.113, 0.103] | [-0.095, 0.087] | 0.928 | 0.9333 | 53.46% | 99.433 | 1 | 8985.322 |
|  | *Cue \* Trial* | -0.03 | -0.035 | -0.035 | -0.20 – 0.14 | [-0.211, 0.142] | [-0.184, 0.112] | 0.704 | 0.6975 | 65.33% | 90.7 | 1 | 12054.973 |

<sup>a</sup>. This table presents results of a linear mixed model predicting subjective pain on medium heat trials as a function of Group (Instructed vs Uninstructed), Cue (Original High vs Original Low), and Trial prior to the first reversal, as well as post-hoc tests in each Group. See Table 1 for additional information about model specification and presentation.
<sup>b</sup>. Estimates based on a linear mixed effects model implemented in the “lmer” function of lme4 (Bates et al., 2015) using the following code: `lmer(PainMedium~Group*Cue*Trial+(1+Cue*Trial|Subject))`.
<sup>c</sup>. Estimates based on a linear mixed effects model implemented in the “lme” function of nlme (Pinheiro et al., 2021) including autoregression using the following code: `lme(Pain~Group *Cue*Trial, random=~1+Cue*Trial|Subject, correlation=corAR1(), na.action=na.exclude)`.
<sup>d</sup>. Estimates based on Bayesian model linear mixed models using the “brms” function (Bürkner, 2017) using the following code: `brm(Pain~Group*Cue*Trial+(1+Cue*Trial|Subject,prior=set_prior("normal(0,2.5)", class="b"), save_all_pars=TRUE, silent=TRUE, refresh=0, iter = 4000, warmup = 1000)`. Posterior estimates and the Region of Partial Equivalence were obtained using the “describe\_posterior” function from the package BayesTestR (Makowski et al., 2019a) and interpreted as in (Makowski et al., 2019b). The Region of Partial Equivalence (ROPE) was defined as [-0.17, 0.17] across all participants, [-.172, .172] when restricted to the Instructed Group, and [-.168, .168] when restricted to the Uninstructed Group.

### Cue-based expectations and cue effects on pain update as contingencies reverse

We next tested whether expectations and cue effects on pain updated as contingencies reversed, and whether they did so differently as a function of instruction. We computed an expectancy rating difference score (Original High Pain expectancy – Original Low Pain expectancy; see Figure 3B) for each pre-block rating and measured effects across the entire task as a function of Group and Phase (i.e. Original vs. Reversed Contingencies; see vertical dashed lines in Figure 1D and 3A). We observed a main effect of Phase (B = -2.03, p < .001), indicating that differential expectations varied as contingencies reversed, and significant Group x Phase interaction (B = 4.12, p < .001). Post-hoc analyses indicated that only the Instructed Group reported differences in expectation that varied significantly as a function of Phase, whereas the Uninstructed Group showed weaker variations in expectations as contingencies reversed (see Figures 3A and 3B).

**Figure 3.**
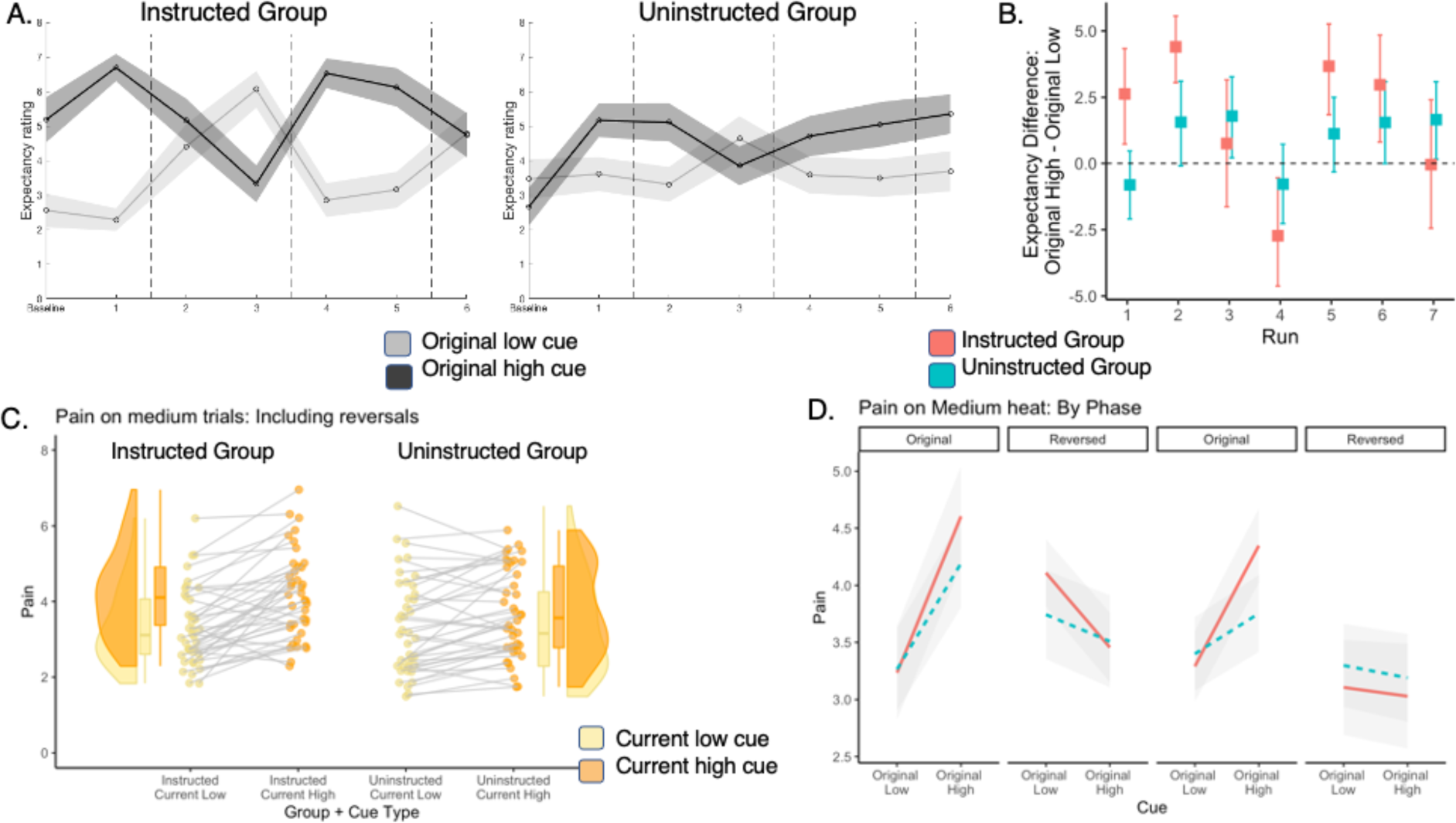
Expectations and pain ratings update as contingencies change. We analyzed cue-based expectations and the effects of cues on pain ratings in response to medium heat across the entire task, including reversals. Reversals were coded relative to instructions in the Instructed Group and relative to experience in the Uninstructed Group (see Figure 1C). A) Expectancy ratings across the entire task. Both groups updated expectations as contingencies reversed. B) Cue-based differences in expectancy. The Instructed Group (Red) shows larger differences in expectancy as a function of phase, although both groups show significant Cue x Phase interactions across the task, indicating that both instructions and experiential learning dynamically shape expectations. C) Effects of current cue contingencies on subjective pain. We analyzed Cue x Phase interactions on pain to evaluate whether individuals report higher pain with the cue that is currently paired with high heat (Original High Cue on original contingency blocks, Original Low Cue on reversed blocks). Both groups reported higher pain when medium heat was paired with the current high cue relative to the current low cue. D) Pain reversals are larger in Instructed Group participants. As with expectancy ratings, both groups showed significant reversals of cue effects on subjective pain as contingencies changed, but reversals were larger in Instructed Group participants.

We next examined pain reports in response to medium heat across all trials, including reversals (see Figures 3C, 3D, and Supplemental Figure S3). Bayesian model comparison using a normal distribution indicated the most likely model included fixed effects of Group, Cue, Phase, and Trial, with random intercepts and slopes (see Supplemental Results). All models revealed significant Cue x Phase interactions on pain, indicating that cue effects on pain varied as contingencies reversed (see Figures 3C and 3D and Table 3). This effect was sufficient to reject the null hypothesis of no interaction, as fewer than 1% of estimates fell within the ROPE (see Table 3). All models also revealed main effects of Cue, such that individuals reported higher pain in response to the original high pain cue than the original low pain cue (see Table 3), and main effects of Phase, such that pain was higher on original contingencies relative to reversals (see Table 3). Main effects of Cue and Phase were significant in all frequentist analyses, but evidence was not sufficient to reject the null (i.e. not practically significant) based on Bayesian models (see Table 3). Finally, frequentist analysis approachs revealed significant Group x Cue x Phase interactions, driven by stronger reversals of Cue effects in the Instructed Group (see Figure 3D), although Bayesian models indicated that group differences were not practically significant. Post hoc analyses conducted separately by Group indicated nearly 100% probability of positive Cue x Phase interactions in each group, although evidence was only sufficient to reject the null hypothesis in the Instructed Group (see Table 3). We observed similar results when we restricted analyses to pain ratings from the 35 participants with useable fMRI data, although the Group x Cue x Phase interaction was no longer significant in any model; see Supplementary Table S2 for complete details. We also observed consistent findings when we tested the model with a beta distribution, which was found to provide better fits based on posterior prediction (see Supplement Results and Supplementary Table S3). Thus predictive cues shape pain perception even as contingencies change, whether or not participants are instructed about contingencies. In addition, reversals may be slightly larger in participants who are explicitly instructed about contingencies and reversals, however group differences were not practically meaningful based on Bayesian statistics.

**Table 3.**
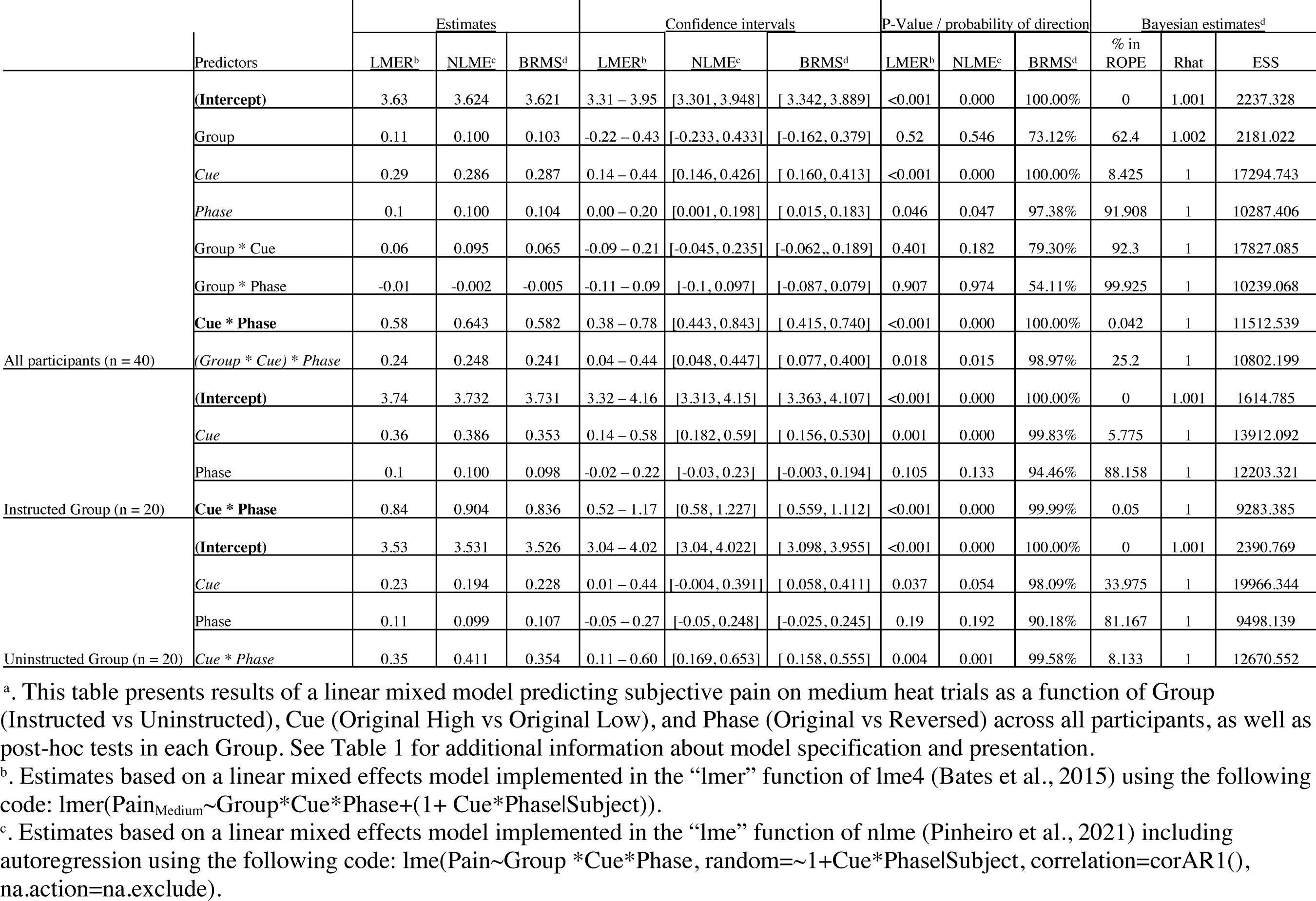
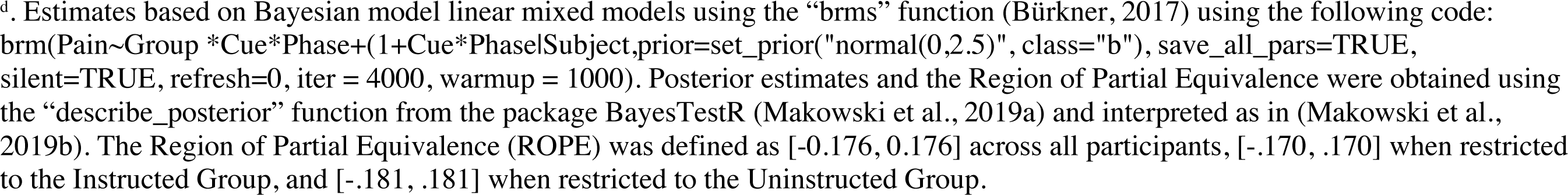
Multilevel model evaluating effects of Group, Cue, and Phase on medium heat pain across the entire task.^a^

### Brain mediators of dynamic cue effects on subjective pain during pain reversal learning

Our behavioral analyses indicated that predictive cues modulated expectations and subjective pain, and that cue effects on both outcomes updated as contingencies reversed. We next asked which brain regions mediated cue effects on pain on the critical medium heat trials, consistent with our previous work (Atlas et al., 2010). We performed two voxel-wise multilevel mediation analyses: 1) a search for brain mediators of current cue effects on pain, i.e. the Cue x Phase interaction (see Figure 4A, Supplementary Figures S4-S6, and Supplementary Tables S4-S5), and 2) a search for mediators of original cue effects on pain, controlling for current contingencies (see Figure 5A and Supplementary Figures S7-S10, and Supplementary Tables S6- S7). For both models, Path *a* evaluates the effect of predictive cue on brain response to medium heat, Path *b* captures the association between brain response and pain, controlling for cue, and the mediation effect (Path a*b) tests whether brain responses contribute to variance between cues and subjective pain on medium heat trials. We evaluated mediation overall irrespective of group, and with Group as a potential moderator of all paths.

**Figure 4.**
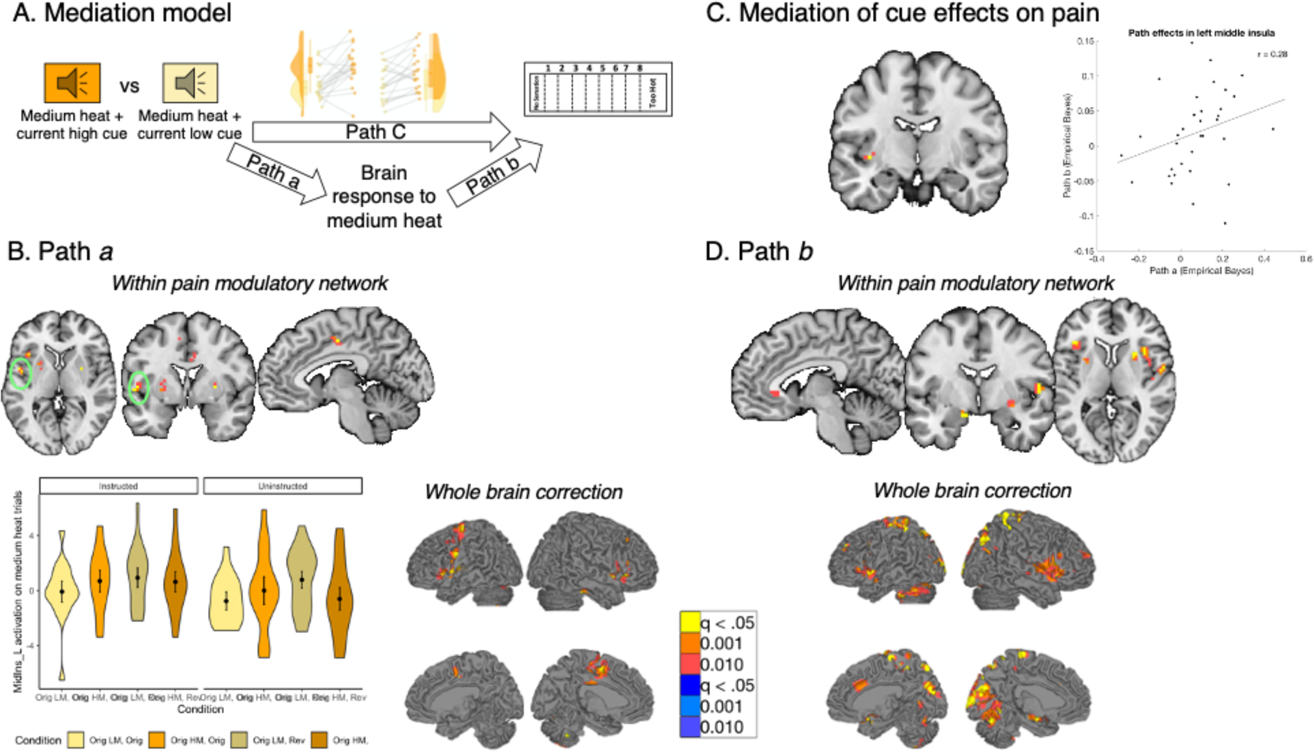
Mediation of current cue effects on medium heat pain. We examined brain mediators of current contingency effects on perceived pain on medium heat trials. Results are FDR-corrected within pain modulatory regions and across the whole brain. A) Mediation model. We tested for brain regions that mediate the effects of current cue contingencies on subjective pain, corresponding to the reversals we observed (Figure 3). B) Path A: effects of current contingencies. Path a identifies brain regions that show greater activation with the current high pain cue (e.g. Original High Cue during original contingencies, Original Low Cue during reversed contingencies), relative to the current low pain cue. Within pain modulatory regions, we observed positive Path a effects in the left middle insula, dorsal anterior cingulate, bilateral putamen, and left anterior insula. We extracted trial-by-trial estimates from the left middle insula and visualized average responses as a function of Group, Cue, and Phase (bottom left). Both groups showed greater left insula activation when medium heat was preceded by the Current High Cue, and cue effects did not differ by group. Whole brain FDR-correction (bottom right) additionally identified positive Path a effects in the bilateral DLPFC and lateral PFC. C) Mediation of current cue effects on pain. We observed significant mediation by the left anterior insula. Extracting responses within this region indicated that individuals who showed larger cue effects on insula (i.e. Path a effects) showed marginally stronger associations between insula activation and subjective pain (i.e. Path b; r = 0.28, p = .098). D) Path b: associations with pain controlling for cue. Path b regions are positively associated with pain, controlling for cue (and temperature, since we tested only medium heat trials). We observed positive Path b effects in the subgenual ACC, bilateral amygdala, bilateral anterior insula, and other regions within the pain modulatory network (top). For additional regions identified in whole brain search and uncorrected results, see Supplementary Figures S4-S6 and Supplementary Tables S4-S5.

**Figure 5.**
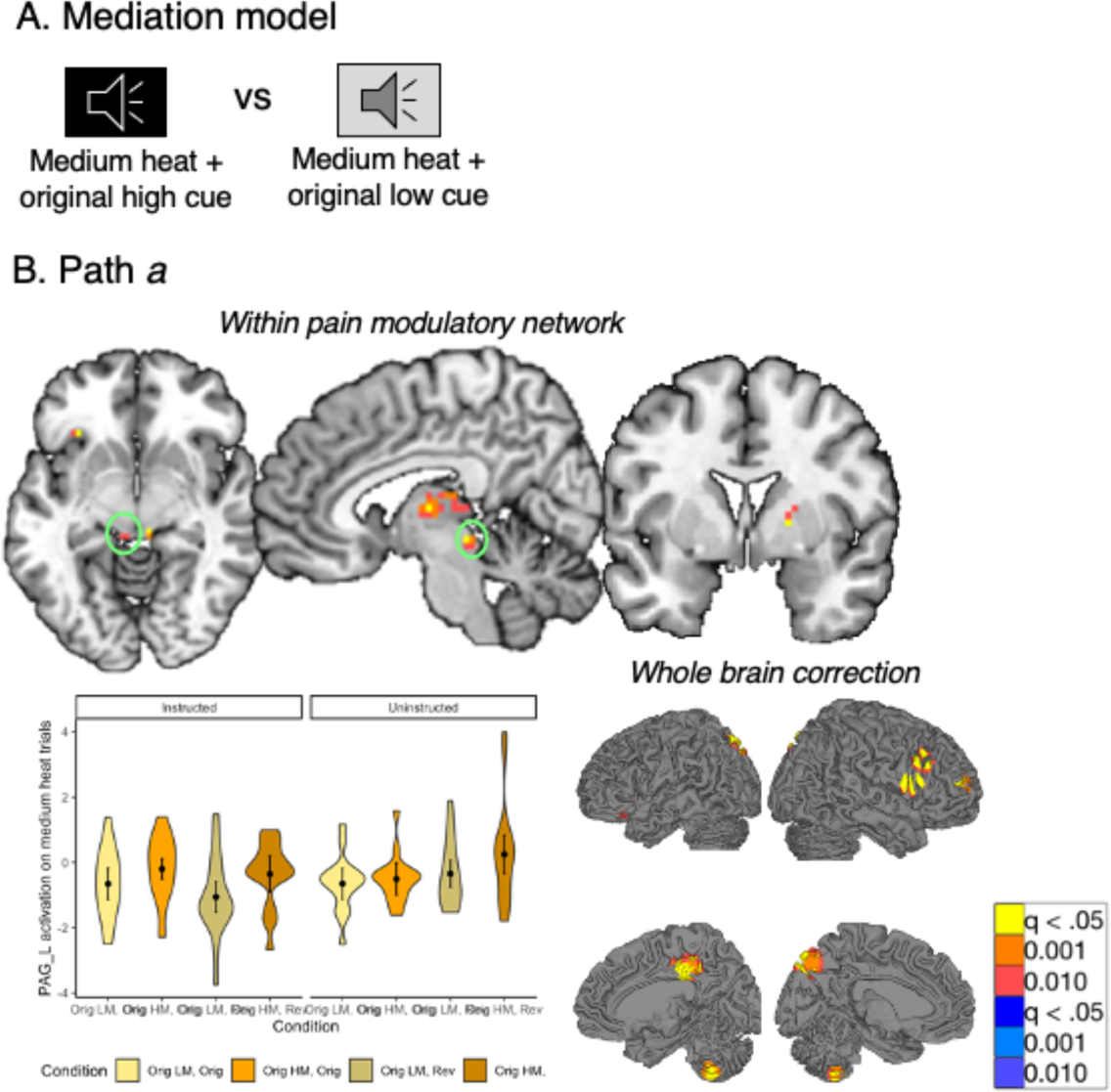
Original cue effects on medium heat pain. We conducted a second mediation analysis to isolate effects of original contingencies, controlling for current contingencies. A) Effects of original contingencies. The goal of our second mediation analysis was to specifically identify regions that continued to respond to the original contingencies across the entire task, regardless of reversals. B) Path a: Regions that show greater activation to original high pain contingencies despite reversals. Path a identified regions that showed greater activation to the Original High Cue across the entire task. Within pain modulatory regions (top), the periaqueductal gray, thalamus, and putamen all continued to show higher activation when medium heat was paired with the original high pain cue regardless of Phase. Extracting trial-by-trial responses from the PAG (bottom left) confirmed that this region showed greater heat-evoked activation with the Original High Cue during both original and reversed contingencies and that effects were present in both the Instructed Group and the Uninstructed Group. See Supplementary Figure S6 for means within other Path a regions. Whole brain correction identified additional effects in the right DLPFC, precuneus, and cerebellum (see Supplementary Figure S9 and Supplementary Table S6). Whole brain uncorrected results are presented in Supplementary Figure S10 and Supplementary Table S7.

**Table 4.**
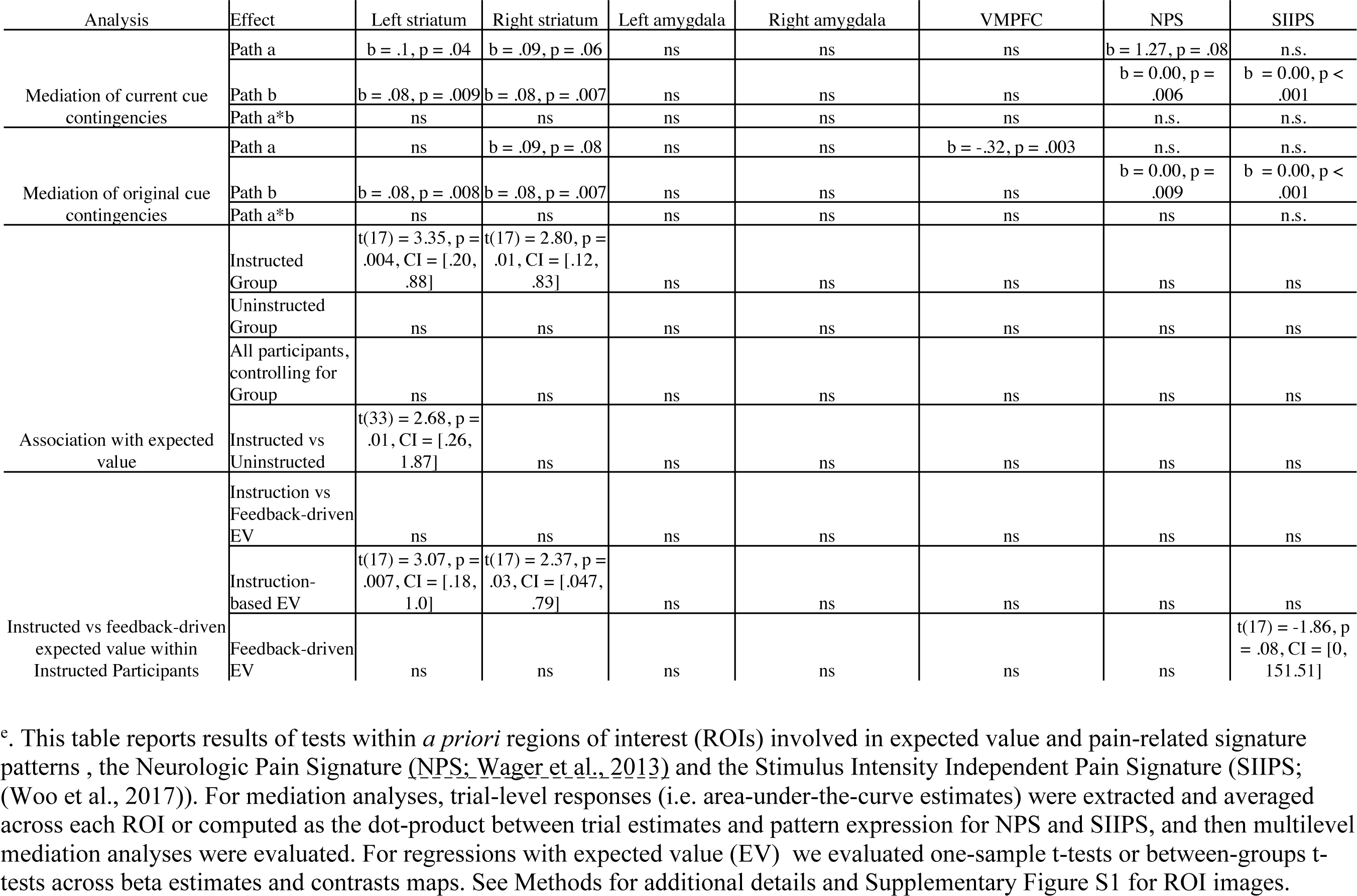
Effects of cues and learning on responses in value-related regions of interest and pain-related signature patterns.^e^

We focus in particular on effects of current contingencies (i.e. Cue x Phase interactions) on pain to capture the behavioral reversals we observed (Figure 4). Our main report focuses on results of small volume correction within pain-related regions or whole brain correction, as well as pain predictive signature patterns (see Methods). Exploratory uncorrected whole brain results are reported in Supplementary Figure S6 and Supplementary Table S5.

Path *a* identified regions that showed stronger activation in response to medium heat following high pain cues relative to low pain cues. Within pain modulatory regions, we observed significant positive Path *a* effects (current high cue > current low cue) in the left anterior and middle insula, left putamen, dACC, preSMA, and left VLPFC (see Figure 4B and Table S4).

Extracting trial-level responses confirmed that these regions showed greater activation when medium heat was preceded by the initial high pain cue relative to the initial low pain cue during the original contingences, whereas they showed greater activation when medium heat was paired with the initial low pain cue when contingencies were reversed, and these reversals were observed for both groups (see Figure 4B and Supplementary Figure S4). Whole brain FDR correction additionally indicated positive Path *a* effects in left M1 (see Figure 4B, Supplementary Figure S5, and Supplementary Table S4). Path *a* effects on NPS expression were marginal in the direction of stronger NPS expression when medium heat was preceded by high pain cues (see Table 4), driven by significant effects in the Uninstructed Group (see Supplementary Figure S4). There was no Path *a* effect on SIIPS expression (p > 0.4; see Table 4 and Supplementary Figure S4). Within value-related ROIs, we observed significant Path *a* effects on the left striatum and marginal positive associations in the right striatum (see Table 4); no other ROIs showed modulation by current cue contingencies.

We observed positive Path *b* effects (associations with pain, controlling for cue) within bilateral anterior insula, pregenual ACC, bilateral putamen, bilateral amygdala, and right SII (see Figure 4D and Supplementary Table S4). Whole brain FDR-correction also revealed positive Path *b* effects in the bilateral M1, right S1, bilateral superior parietal lobule, and other regions (see Figure 4D, Supplementary Figure S5, and Supplementary Table S4). No negative Path *b* effects survived correction within the pain modulatory mask or whole brain search. We observed significant Path *b* effects on responses to medium heat for both signature patterns, as well as the bilateral striatum (see Table 4).

Finally, we observed significant positive mediation of current cue effects on pain in the left middle insula (see Figure 4C and Supplementary Table S4). Mediation was primarily driven by the covariance between paths a and b, meaning that individuals who showed stronger path *a* effects also showed stronger Path *b* effects, although correlations between paths were marginal (r = 0.28, p = .098; see Figure 4C). We did not observe mediation by the NPS or SIIPS pattern or any value-related ROI (see Table 4) and no additional regions were identified in whole brain search at FDR-corrected thresholds. See Supplementary Figure S6 and Supplementary Table S5 for whole-brain uncorrected results.

Notably, we did not observe significant moderation by Group in any of the paths at FDR-corrected thresholds, both when restricted to pain modulatory regions or whole brain correction There was also no moderation of the SIIPS, NPS, or value-related ROIs in any path. Uncorrected results are reported in Supplementary Figure S7 and Supplementary Table S5, but evidence of moderation was minimal. This suggests that brain mechanisms of cue effects on subjective pain are similar whether individuals learn through instruction or experience, despite stronger influences of cues on pain within the Instructed Group.

### Responses in periaqueductal gray and thalamus maintain initial contingencies despite reversals

While the main mediation analysis isolated effects of current contingencies on medium heat trials including reversals, some regions may show sustained responses to initial contingencies. We therefore conducted a second mediation to identify regions that responded to original contingencies and did not reverse as contingencies changed (Figure 5). We searched for mediators of original cue effects and controlled for current contingencies by including current contingencies as a covariate. These effects are thus most likely to be driven by responses during the reversed runs. We were most interested in Path *a*, which identified regions that showed stronger activation in response to cues that were originally paired with high pain relative to cues that were originally paired with low pain (see Figure 5). Within pain modulatory regions, we observed significant positive Path *a* effects (original high cue > original low cue) controlling for current contingencies in the periacqueductal gray (PAG), bilateral medial thalamus, right putamen, right lateral prefrontal cortex, and the orbital part of the left inferior frontal gyrus (see Figure 5B and Supplementary Table S6). As visualized in Figure 5B and Supplementary Figure S8, extracting trial-level responses from these regions confirmed that they showed elevated activation when medium heat was paired with the original high pain cue relative to the original low cue, regardless of whether contingencies had reversed. Whole brain correction additionally revealed positive effects of original contingencies in the right DLPFC, right anterior PFC, and the bilateral superior parietal lobule (see Figure 5B, Supplementary Figure S9, and Supplementary Table S6). ROI-wise analyses indicated that only the VMPFC was significantly modulated by initial contingencies (see Table 4); this region was also identified in whole brain uncorrected results (see Supplementary Table S7 and Supplementary Figure S10). There were no effects of original cues on the NPS or SIIPS (all p’s > 0.2). No brain regions mediated original cue effects on pain based on corrected analyses, whether restricted to pain-related regions or whole brain correction. We observed significant positive Path *b* effects (associations with pain, controlling for cue) in the bilateral anterior insula, right dorsal posterior insula, and bilateral putamen within pain modulatory ROIs, and widespread pain-related activation with whole brain correction (see Supplementary Figure S9 and Supplementary Table S6). We also observed significant Path B effects on the NPS, SIIPS, and the bilateral striatum (see Table 4).

We did not identify any regions that mediated effects of original contingencies on pain, consistent with the fact that pain updated as contingencies changed. We also did not observe significant moderation by Group in any of the paths at FDR-corrected thresholds in any of our analyses. This is consistent with our mediation of current contingencies, and confirmed by similar effects shown by both groups when we extracted responses in pain modulatory regions that showed Path A effects in either mediation analysis (i.e. Figure 4B, Figure 5B, and Supplementary Figures S4 and S8). Uncorrected whole brain results are reported in Supplementary Table S7 and Supplementary Figure S10.

### Quantitative models reveal that instructed participants reverse expectations upon instructions and learning is faster in uninstructed participants

We observed no group differences in the effects of cues on brain responses to noxious stimuli, suggesting that pain is mediated similarly whether or not participants are instructed about contingencies. However, learning might still differ between groups, as our mediation models and analyses by Phase assume that expectations update completely upon reversal, either through instruction in the Instructed Group or when contingencies reverse in the Uninstructed Group. However, if learning proceeds more dynamically (i.e. continuously as a function of pairings between cues and temperatures), we would not capture this with categorical models that assume immediate changes upon reversal.

To formally examine these dynamics, we applied a quantitative model of instructed reversal learning (Atlas et al., 2016; Atlas and Phelps, 2018) which accounts for how expectations update dynamically as a function of both experience and instruction. The model includes two parameters: ɑ, a standard learning rate that captures the extent to which expected value (EV) updates in response to prediction errors, and ⍴, which guides whether and how EV reverses upon instruction (see Methods). Here, we extended this model to predict subjective pain on medium heat trials. This model accounted for variations in pain reports better than other plausible models, including a standard Rescorla-Wagner model without the ⍴ parameter and a hybrid model of adaptive learning modified to reverse upon instruction ((Atlas et al., 2019); see Supplemental Methods for complete information on goodness-of-fit and model comparison).

Consistent with our task manipulation, instructed reversal parameters (i.e. ⍴) varied as a function of Group (fit to individuals: t(38) = 3.013, p = .005; see Figure 6A), such that participants in the Instructed Group showed larger reversals at the time of verbal instruction (fit to individuals: Instructed: *M* = .62, *SD* = .35; Uninstructed: *M* = .31, *SD* = .31). This confirms our task manipulation (instructed reversals should only be seen in the group that was exposed to instructions) and validates the model’s application to subjective pain. Consistent with our previous work on instructed threat learning (Atlas et al., 2016), learning rates (i.e. ɑ) were close to zero in the Instructed Group (*M* = .065, *SD* = .22), indicating that there was little additional learning as a function of experience between instructed reversals. Learning rates were indeed higher in the Uninstructed Group (*M =* .28, *SD* = 0.34), and differed significantly between groups (fit to individuals: t(38) = -2.32, p = .026). Differences in ⍴ and ɑ were observed when models were fit to individuals, and when they were fit across the group using a jack-knife model fitting procedure (see Supplemental Results). Thus expected value updates primarily upon instruction in the Instructed Group with very little additional learning between reversals (consistent with the Cue x Phase interactions we modeled behaviorally), whereas individuals in the Uninstructed Group update expected value over time as a function of experience, i.e. pairings between cues and heat, as depicted in Figure 6B.

**Figure 6.**
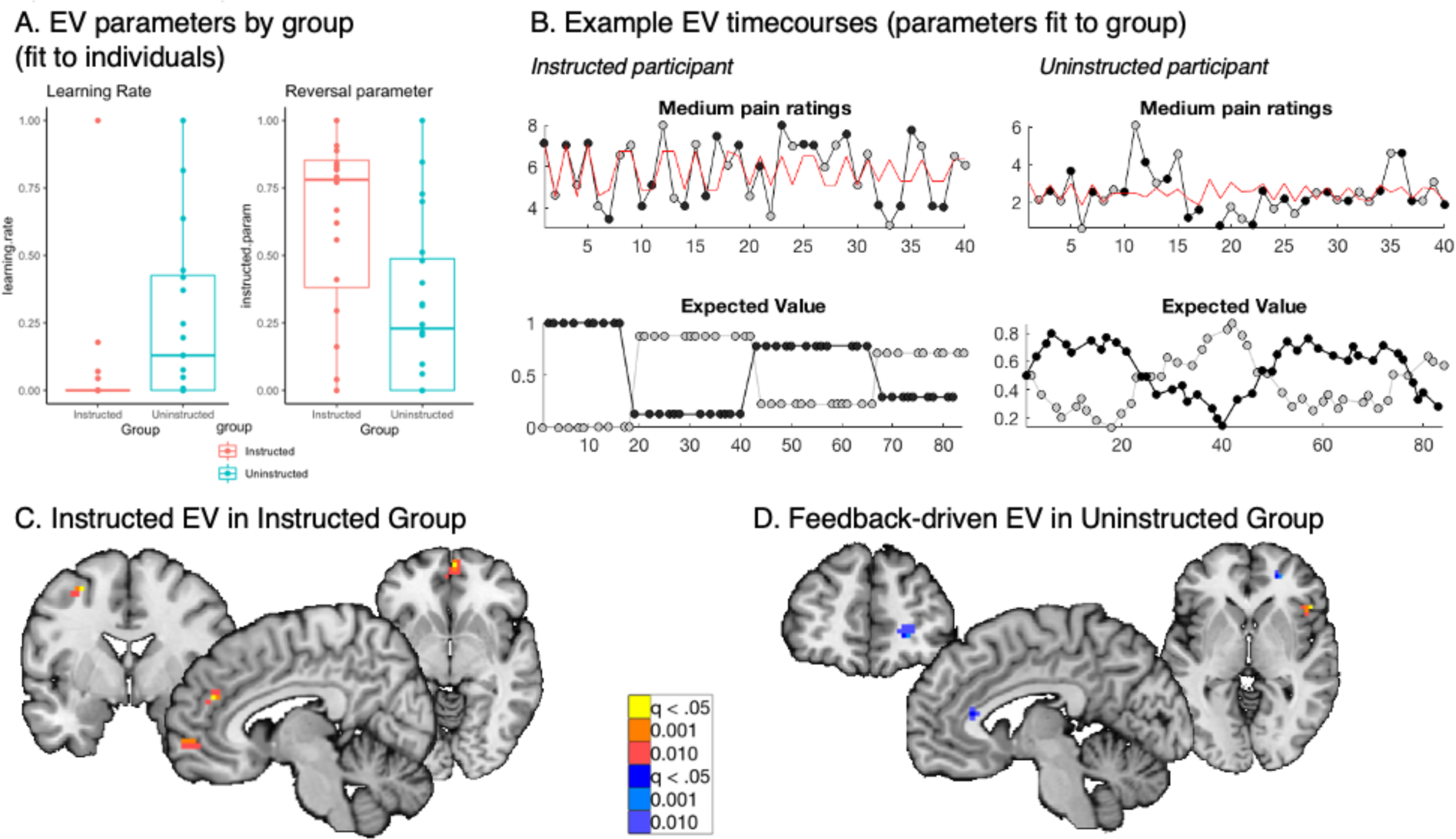
Instructed learning model fit to pain on medium heat trials. We fit a computational model of instructed reversal learning (Atlas et al., 2016) to pain reports on medium heat trials to isolate the dynamics of expected value and how expected value updates with instruction. A) Group differences in learning parameters. Fitting models to individuals revealed group differences in learning rate (α, left), such that participants in the Uninstructed Group (blue) showed stronger updates of expected value in response to prediction errors relative to the Instructed Group (red), whereas the Instructed Group showed stronger reversals at the time when instructions were delivered, based on the instructed reversal parameter (ρ, right). B) Predicted timecourse of expected value based on jack-knife model fits. We used model parameters from a jack-knife model fitting procedure (see Methods and Supplementary Results) to generate predicted timecourse of expected value (EV) for each group. Here we depict model predictions for an example participant in the Instructed Group (left) and the Uninstructed Group (right). As shown in the bottom row, EV reverses immediately upon instruction in the Instructed Group and reverses more gradually in the Uninstructed Group. C) Neural correlates of instructed EV. Across the Instructed Group, we observed positive associations between EV and responses to medium heat in the insula, MPFC, VMPFC, and DLPFC. No regions showed negative associations with EV. D) Neural correlates of feedback-driven EV. Within the Uninstructed Group, we observed positive associations between experience-based EV and right anterior insula responses to medium heat, and negative associations in the rACC and right PFC. See also Supplementary Figure S12 and Table S8.

### Expected value dynamically modulates responses to noxious stimulation, with differences between groups in the rostral anterior cortex

We next searched for neural correlates of dynamic expected value signals on medium heat trials. We used the learning time- course generated from fits to each group and searched for regions that correlated with expected value (EV), which updated with instruction and varied little between instructed reversals in the Instructed Group and depended more critically on trial-by-trial outcomes in the Uninstructed Group, which displayed higher learning rates. Figure 6B depicts example EV timecourses using the same parameters that were used to evaluate associations between EV and medium heat-evoked brain responses in each group.

We first examined correlations with expected value separately for each group. Within the Instructed Group, we observed significant positive associations with instruction-based EV in the right anterior insula within the pain modulation network, and observed positive associations in the VMPFC, DMPFC, and left DLPFC in whole-brain corrected results (see Figure 6C, Supplementary Figure S11, and Supplementary Table S8). Within the Uninstructed Group, we observed negative associations with feedback-driven EV within the pain modulation network in the rACC, and whole brain corrected analyses revealed additional negative associations in the right anterior PFC and precuneus, as well as positive associations in the right anterior insula/ inferior frontal gyrus (see Figure 6D, Supplementary Figure S11, and Supplementary Table S8). Whole brain uncorrected results for each group are presented in Supplementary Figure S12 and Supplementary Table S9. ROI-wise tests within the Instructed Group indicated significant associations with instructed EV in the bilateral striatum, but not the amygdala or VMPFC, whereas there were no associations between feedback-driven EV and activation in any value ROIs in the Uninstructed Group (see Table 4). There were no significant associations between EV and NPS or SIIPS expression for either group (see Table 4).

We next used robust regression to identify brain regions whose associations with EV differed between groups (see Figure 7). FDR correction within *a priori* pain modulatory regions revealed significant differences in rACC activation (see Figure 7A), such that there was a positive association with EV in the Instructed Group (CI =[0.2938, 1.3172], t(17) = 3.32, p < .001) and a negative association in the Uninstructed Group (CI =[-2.54, -0.96], t(16) = -4.69, p < .001). Whole brain FDR corrected comparisons between groups revealed positive differences (Instructed > Uninstructed) in the rACC, MPFC, left temporal pole, left TPJ, and right precuneus and negative differences (Uninstructed > Instructed) in the right VMPFC / mOFC, left DLPFC, left IPL, right cerebellum, left lateral PFC, and right DMPFC (see Figure 7B, Supplementary Figure S12, and Supplementary Table S8). ROI-wise tests within *a priori* value related regions indicated that groups differed in the left striatum, driven by positive associations in the Instructed Group (see Table 4).

**Figure 7.**
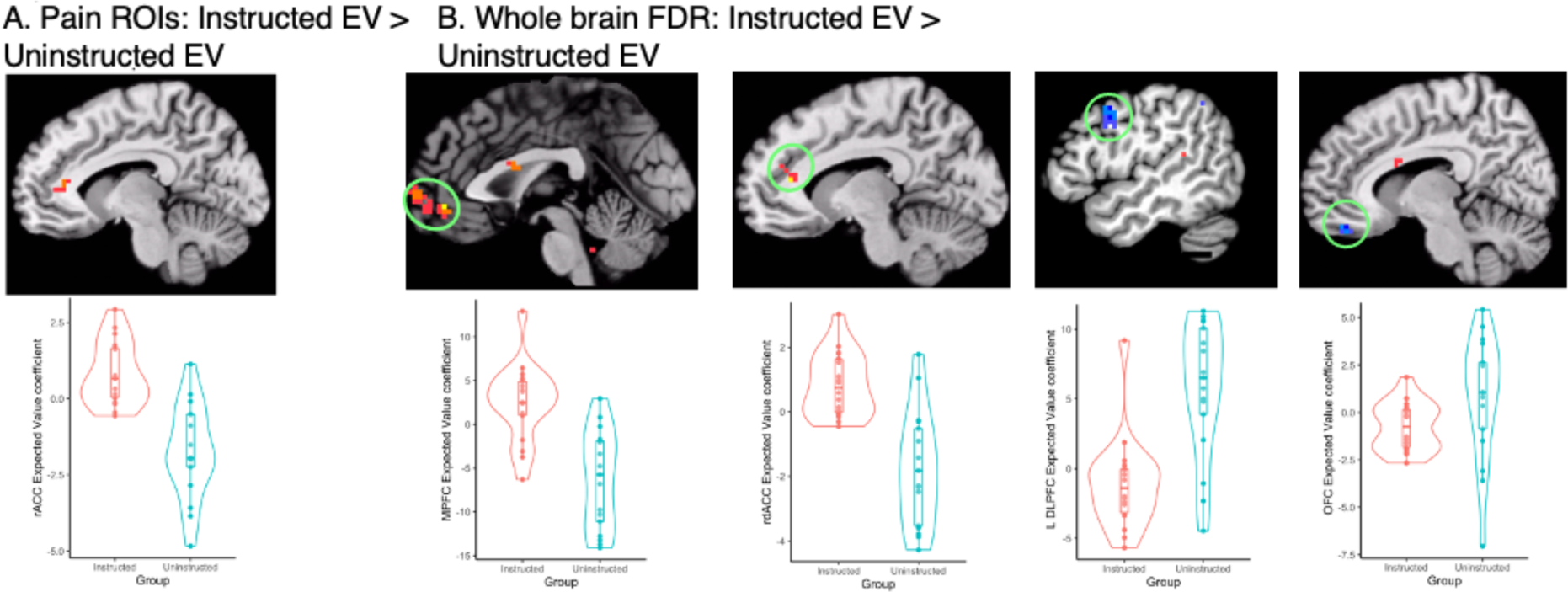
Group differences in associations with expected value on medium heat trials. We used the timecourse of expected value (EV) based on fitting computational models to pain reports from each group (see Figure 6) to isolate the neural correlates of instructed and uninstructed expected value during pain processing. We examined associations between brain responses to medium heat and the timecourse of EV and used robust regression (Wager et al., 2005) to compare groups. (A) Group differences in expected value within pain modulatory regions. The rostral anterior cingulate cortex (rACC) showed positive associations with EV within the Instructed Group (red) and negative associations within the Uninstructed Group (blue). B) Group differences in expected value based on whole brain search. Several regions showed significant differences in associations with EV, including the MPFC, rdACC, left DLPFC, and OFC/VMPFC. See also Supplementary Figure S12 and Supplementary Table S8.

Robust regression did not identify any significant associations across all participants (i.e. main effect, controlling for group) between EV and responses to medium heat within pain-modulatory regions. However, whole brain FDR correction revealed positive associations with EV across participants in the left DLPFC and negative associations in the right anterior PFC, left IPL, and left inferior frontal gyrus (see Supplementary Figure S13 and Supplementary Table S8). Additional results from uncorrected whole-brain exploratory analyses are reported in Supplementary Figure S13 and Supplementary Table S9.

Finally, we searched for correlates of instructed and feedback-driven EV signals within Instructed Group participants, to test whether brain responses were preferentially related to instructed or feedback-driven learning within participants exposed to both types of information. Controlling for uninstructed EV, instructed EV was positively associated with activation in the midbrain near the substantia nigra, and negatively associated with activation in the right precentral gyrus, based on whole brain correction (see Supplementary Figure S14 and Supplementary Table S10). No regions showed preferential associations with uninstructed EV or significant differences between instructed and uninstructed EV based on whole brain correction. For complete results, please see Supplementary Results.

## Discussion

We measured whether cue-based expectancy effects on pain and brain responses to noxious heat update dynamically as contingencies change, and whether these relationships vary as a function of whether individuals learn through instruction or experience. All participants demonstrated robust cue-based expectancy effects on pain, consistent with previous work from our group and others (Colloca et al., 2008a; Atlas et al., 2010; Wiech et al., 2014; Fazeli and Büchel, 2018; Jepma et al., 2018; Michalska et al., 2018; Koban et al., 2019; Abend et al., 2021). Here, we provide new evidence that these predictive cue-based expectancy effects on pain update as contingencies change, whether reversals are accompanied by instructions or learned through experience. Reinforcement learning models indicated that these effects emerge dynamically, consistent with error-driven learning. We observed dissociations in the associations between expected value and brain responses to heat in several brain regions, including the rostral anterior cingulate cortex (rACC), which was positively associated with expected value in the Instructed Group and negatively associated in the Uninstructed Group. Finally, several pain-related regions including the left anterior insula updated dynamically as contingencies changed regardless of group, whereas the periacqueductal gray (PAG) and thalamus responded to the original contingencies throughout the task. Here we discuss these findings and their implications for future work and our understanding of pain, predictive processing, and the interaction between learning and instructed knowledge.

Our study presents a novel examination of expectancy-based pain modulation during reversal learning. Pain reports reversed as contingencies changed, whether or not participants were instructed about contingencies. This suggests that individuals use both instructed knowledge and experience to generate cue-based expectations, which in turn modulate subjective pain. Dynamic cue effects on pain were mediated by the left anterior insula, a region that was previously found to mediate the effects of instructed cues on pain reports in the absence of reversals (Atlas et al., 2010). In addition, we observed significant reversals of cue effects in several pain-related regions, including the left insula, dACC, and putamen. In contrast, we observed sustained responses to initial contingencies in the PAG and thalamus, among other regions. Together, our findings build on prior work demonstrating that cue-based expectations modulate pain-related brain responses to noxious heat (Atlas et al., 2010; Fazeli and Büchel, 2018; Jepma et al., 2018; Sharvit et al., 2018; Koban et al., 2019) and further indicate a division within these systems. Insula, dACC, and putamen are flexible and update as contingencies change (like pain), whereas the periaqueductal gray and thalamus maintain initial contingencies despite reversals. We return to these dissociations later in the Discussion. Interestingly, cue effects on brain responses did not differ between groups, i.e. as a function of whether individuals were instructed about contingencies. This suggests that responses to contingency reversals and links with subjective pain are similar regardless of whether individuals learn through experience or instruction. Thus once an expectation or prediction is generated, it has similar effects on downstream responses regardless of how contingencies were established.

In contrast to the similar effects of predictive cues on brain responses, quantitative learning models revealed differences between groups in how pain and pain-related responses updated dynamically from trial to trial. Individuals who were exposed to instructed reversals updated pain immediately upon instruction without additional learning from intermittent reinforcement, whereas individuals who learned purely from experience had higher learning rates, meaning they updated expectations as a function of pairings between predictive cues and heat outcomes. This is consistent with prior work focusing on autonomic arousal during threat learning (Atlas et al., 2016) and confirmation bias in reward-related decision making (Doll et al., 2009). However, whereas our previous work showed that these factors were also associated with dissociations in value-based systems, such that the amygdala responded to feedback while the striatum and OFC updated with instruction (Atlas et al., 2016; Atlas, 2019), we observed somewhat different patterns in brain regions involved in value based learning in our pain paradigm. Consistent with findings in appetitive and aversive learning, ROI-wise analyses indicated that the striatum updated with instruction, along with the MPFC, DMPFC, and left DLPFC. The right insula tracked expected value, with greater activation when high pain was expected, whether individuals learned from experience or through instruction.

Perhaps most strikingly, several regions along the wall of the medial prefrontal cortex, including the rostral anterior cingulate cortex (rACC), DLPFC, MPFC, and medial OFC showed differential associations with expected value (EV) depending on group. Activation in rACC and MPFC was positively associated with EV in the Instructed Group (i.e. greater activation with high pain expectancy), and negatively associated with EV in the Uninstructed Group (i.e. greater activation with low pain expectancy). The rACC also showed preferential associations with instruction-based EV when we compared associations between instruction-based and experience-based EV within the Instructed Group. The rACC has been implicated in numerous studies of placebo analgesia and expectancy-based pain modulation (Petrovic, 2002; Bingel et al., 2006; Eippert et al., 2009; Geuter et al., 2013), and is a key component of the opioidergic endogenous pain modulation circuit (Zubieta, 2005; Wager et al., 2007; Navratilova et al., 2015). Our findings suggest that the rACC shows different dynamics and different patterns of responses to threat during reversal learning depending on whether individuals are exposed to contingency instructions. The rACC and PAG show increased functional connectivity under placebo (Bingel et al., 2006) and this connectivity is linked to endogenous opioid binding (Eippert et al., 2009).

We observed striking dissociations between these regions: the rACC responded to expected value and updated based on instructed or uninstructed learning, whereas the PAG maintained initial contingencies throughout the task. Future studies should examine the relationship between these regions and whether placebo-based connectivity differs in more dynamic environments or as a function of instruction.

We also observed differences between groups in the association between value-based processing in the left DLPFC, which was positively associated with EV in the Uninstructed Group and negatively associated with EV in the Instructed Group. Similar to the rACC, the left DLPFC has been implicated as playing a modulatory role in expectancy-based modulation in many previous studies (Lorenz et al., 2003; Wager, 2004; Atlas and Wager, 2014), consistent with its role in cognitive control and executive function (Miller and Cohen, 2001). Our mediation analyses indicate that it is dynamically modulated by pain-predictive cues, with greater activation when high pain is expected, but our computational models suggest that associations with dynamic expected value differ between groups. This builds on our previous work that indicated that DLPFC mediates cue effects on pain (Atlas et al., 2010), but that there are individual differences in the magnitude of these effects, such that some individuals activate DLPFC in response to high pain expectancy, while others show greater DLPFC activation in response to low pain cues. Future studies should use causal approaches such as TMS to better understand the contribution of DLPFC to expectancy-based pain modulation (Krummenacher et al., 2010).

Finally, we observed group differences between dynamic expected value and heat-evoked activation in the medial OFC/VMPFC driven by more negative associations in the Instructed Group and positive associations in the Uninstructed Group. The focus of activation in this region fell below our *a priori* ROI, which was selected based on previous studies of fear conditioning, and thus we did not see significant associations with EV or group differences in ROI-based analyses. Heat and aversive experiences usually elicit deactivation in the mOFC (Kong et al., 2010; Atlas et al., 2014), and thus our findings of negative associations with EV in the Instructed Group are consistent with previous work in expectancy-based pain modulation (Atlas et al., 2010). Findings in the Instructed Group also build on previous work indicating that this region updates upon instruction in both appetitive and aversive learning (Li et al., 2011a; Atlas et al., 2016) and that OFC value signals are sensitive to higher order knowledge across species (Wilson et al., 2014; Lucantonio et al., 2015; Schuck et al., 2018). We observed much greater variability in the associations between OFC activation and EV in the Uninstructed Group, however, such that some individuals showed negative associations with EV, whereas others showed positive associations. Learning rates varied widely across participants, which might explain this increased variability within the Uninstructed Group.

### Future directions and outstanding questions

This work highlights several promising avenues of inquiry that should be addressed in future work, in addition to those highlighted above. Comparisons between appetitive and aversive learning would reveal whether the differences observed here are driven by threat-specific processes or general differences in adaptive learning and flexibility. We did not include a group that underwent initial learning in the absence of instructions and then received instructed reversals, which would provide insights on whether or not learned associations can be reversed on the basis of higher order knowledge, consistent with dissociations observed in studies of placebo (Benedetti et al., 2003; Scott M Schafer et al., 2015). In the present analyses, we focused on within-subjects effects and did not examine how these differences vary across individuals as a function of factors such as anxiety, which has previously been shown to impact adaptive learning (Browning et al., 2015). However, we recently examined how instructed reversals impact pain expectations in youth with clinical anxiety and found that youth with and without anxiety showed similar responses to expectancy and instruction, although youth with anxiety showed greater autonomic arousal during pain anticipation (Abend et al., 2021). Future analyses may relate anticipatory responses to the cues themselves with the responses to noxious heat that we focused on here, as well as autonomic responses during anticipation and in response to heat.

### Conclusion

Together, these findings reveal that instructions and learning lead to both interactive and dissociable processes even within individuals. We view these findings in light of theories on the relationship between conditioning and expectancy (Rescorla, 1988; Kirsch, 1997; Kirsch et al., 2004) and long-standing debates about whether placebo effects depend on conditioning or expectancy. We suggest that considering the brain mechanisms that mediate dynamic expectancy-based pain modulation shines new light on these distinctions. The human brain contains parallel pain modulatory circuits that i) update as contingencies change (e.g. insula), ii) continue to respond to initial contingencies regardless of whether they were learned through instruction or experience (e.g. PAG), or iii) respond to experiential learning differentially as a function of whether or not individuals were exposed to instructions (e.g. rACC). These findings indicate that we gain new insights on clinically relevant outcomes from measuring how instructions and learning interact to shape outcomes, rather than assuming that circuits and processes are sensitive to either expectancy or conditioning. Understanding these processes in clinical populations may shed light directly on the mechanisms of therapeutic interventions, for example the interplay between instructed and exposure-based interventions in cognitive behavioral therapy for chronic pain and affective disorders.

## Supporting information

Supplemental Methods, Results, Tables, and Figures

## Acknowledgements

This research was funded by the Intramural Research Program of the National Center for Complementary and Integrative Health (Project ZIAAT-000030; PI Lauren Atlas) as well as NIMH Intramural Research Program Project ZIAMH-002782 (PI Daniel Pine). We thank Linda Ellison-Dejewski and Adebisi Ayodele for assistance with participant screening and clinical support. The authors declare no conflicts of interest.

