## Supplemental Methods, Results, Tables, and Figures for "Instructions and experiential learning have similar impacts on pain and pain-related brain responses but produce dissociations in value-based reversal learning"

10 Center Drive, Rm 4-1471

Bethesda, MD 20892

301-827-0214

Author note: This research was funded in part by the Intramural Research Program of the National Center for Complementary and Integrative Health (Project ZIAAT-000030; PI Lauren Atlas) as well as NIMH Intramural Research Program Project ZIAMH-002782 (PI Daniel Pine). Research was approved by the NIH’s Combined Neuroscience Institutional Review Board (Protocol 15-AT-0132, PI: Atlas; ClinicalTrials.gov identifier NCT02446262). The authors declare no conflicts of interest.

**Supplementary Methods**

**FMRI preprocessing and optimal combination.** We used AFNI’s “afniproc.py” program to preprocess functional data and combine across echoes prior to single trial estimation. We performed slice time correction using AFNI’s 3dTshift program then aligned anatomical and functional data using skull stripped images through AFNI’s align_epi_anat.py program. Anatomical data were then normalized to the Montreal Neurological Institute MNI152_T1_2009c template. We used AFNI’s 3dvolreg and 3dAllineate programs to perform motion correction by aligning echo 2 volumes to the minimum outlier and applying the same corrections to each echo. We made sure there was valid data for each echo and each TR prior to combining multi-echo data. We combined across the three echoes using AFNI’s @compute_OC_weights function to generate a weighted combination of the three echoes. These steps were achieved using the following command to afni_proc.py:

afni_proc.py -subj_id ${test_label}_$subj	\

-dsets_me_echo epi_run*_e01.nii \

-dsets_me_echo epi_run*_e02.nii \

-dsets_me_echo epi_run*_e03.nii \

-echo_times 11 22.72 34.44 \

-blocks tshift align tlrc volreg mask combine blur scale regress \

-tcat_remove_first_trs 4 \

-copy_anat anat_${subj}.nii	\

-anat_uniform_method unifize	\

-tcat_remove_first_trs 4	\

-tlrc_base MNI152_T1_2009c+tlrc	\

-volreg_align_to MIN_OUTLIER	\

-volreg_align_e2a	\

-volreg_tlrc_warp \

-mask_epi_anat yes \

-combine_method OC \

-blur_size 4	\

-regress_stim_times ${subj}_onsets_cue.txt ${subj}_heatonsets_HH.txt ${subj}_heatonsets_LL.txt ${subj}_heatonsets_currentHM.txt ${subj}_heatonsets_currentLM.txt ${subj}_onsets_scale.txt \

-regress_stim_labels cue HH LL HM LM scale	\

-regress_stim_types times times times times times times	\

-regress_basis_multi 'BLOCK(1,1)' 'BLOCK(8,1)' 'BLOCK(8,1)' 'BLOCK(8,1)' 'BLOCK(8,1)' 'BLOCK(1,1)'	\

-regress_opts_3dD -gltsym 'SYM: +HH -LL' -glt_label 1 'HvL'	\

-gltsym 'SYM: +HM - LM' -glt_label 2 'CurrentHMvLM' \

-gltsym 'SYM: +HH +HM +LM +LL' -glt_label 3 'HeatFX' -jobs 2\

-regress_motion_per_run	\

-regress_est_blur_epits	\

-regress_est_blur_errts	\

-regress_run_clustsim yes	\

-execute

**Bayesian models.** Bayesian linear mixed models were implemented using the *brms* package (Bürkner, 2017) in R (R Core Team, 1996). All coefficients were evaluated with a Gaussian prior centered on 0 with a 2.5 SD (i.e. a mildly informative conservative prior). For each outcome measure, we evaluated all potential models ranging from intercepts only to a maximal model (Barr et al., 2013) with random slopes for every factor and all potential interactions, then used “bayesfactor_models” from the *bayestestR* package (Makowski et al., 2019a) to identify the most likely model. We then used the “describe_posterior” function (Makowski et al., 2019a) to evaluate probability of direction (similar to a frequentist p-value with values > 95% indicating the effect is likely to exist), and the percentage of the posterior distribution within the region of partial equivalence (ROPE), which can delineate significance. Published guidelines suggest that when if <1% of the distribution is within the ROPE, one can reject the null hypothesis, whereas if >99% is in the ROPE, one can accept the null hypothesis (Makowski et al., 2019b). We also confirmed model specification using “bayesfactor_inclusion”.

We then evaluated the corresponding model using “lmer” from the *lme4* package (Bates et al., 2015 p.4) to provide frequentist statistics and using “lme” from the *nlme* package (Pinheiro et al., 2021) to incorporate autoregression. Results across approaches were largely consistent (see Tables 1-3).

**Instructed reversal learning model and model comparisons.**

We applied our instructed reversal learning model (Atlas et al., 2016) to pain reports on medium heat trials to estimate how quickly expectations update in response to unexpected outcomes (learning rate; ɑ) and how strongly expectations reverse in response to instructed reversals (instructed reversal parameter; ⍴). Our model is reported in detail in our previous work (Atlas et al., 2016; Atlas and Phelps, 2018). The model is a standard reinforcement learning model (Rescorla and Wagner, 1972) that describes how expected value (EV) updates in response to prediction errors in the environment, depending on learning rate. We include an additional parameter, ⍴, which guides whether cues’ expected values (*EV*) are exchanged when instructions are delivered. If ⍴ = 0, each cue’s EV remains stable (i.e. retains the same *EV* prior to instructions) whereas if ⍴ = 1, the EVs of the two respective cues are exchanged. Mathematically for two cues (a and b), this is implemented for trial (t) as:

EV_t+1_(x_b_) = ρ * V_t_(x_a_) + (1-ρ) * V_t_(x_b_)

For complete details, see (Atlas et al., 2016).

The present model fitting procedure differed in that we fit models to pain ratings on medium heat trials, rather than SCR on unreinforced trials. Consistent with our previous work (Atlas et al., 2016; Atlas and Phelps, 2018), the initial EV was set as 0.5 for Uninstructed Group participants, while we used asymmetric initial expected values for Instructed Participants who were informed about contingencies (1 for high pain cue; 0 for low pain cue). Models were fit at the level of the individual, with two free parameters (ɑ and ⍴), and we used 20 iterations with a random starting value for each parameter for each iteration. We minimized the deviance between expected value and pain on medium heat trials. EV was bounded between 0 and 1; however, we used MATLAB’s glmfit.m function to relate EV with pain on medium heat trials using the normal link function, which includes an intercept and therefore accounts for differences in scale between EV and pain reports. Consistent with other analyses, the first trial of every block was not included during model fitting due to novelty responses.

To evaluate goodness-of-fit, we compared the instructed Rescorla-Wagner model described above with three other variations of plausible learning models: a) an instructed Rescorla-Wagner model with initial value, ɑ, and ⍴ as free parameters; b) a traditional Rescorla-Wagner model without an instructed reversal parameter (initial value and ɑ as free parameters); and c) an instructed model of associability (i.e. a Hybrid model; (Li et al., 2011b)) with ⍴, η, and κ as free parameters (Atlas et al., 2019). We computed Akaike’s Information Criterion (Akaike, 1974) for each model and conducted formal model comparison across the four candidate models using Bayesian Model Selection (Stephan et al., 2009).

The instructed Rescorla-Wagner model with pre-determined initial values and free parameters for ɑ and ⍴ (i.e. the same model used in our previous work; (Atlas et al., 2016; Atlas and Phelps, 2018)) fit the data better than any of the alternatives, as indicated by Bayesian Model Selection (implemented in SPM’s spm_BMS.m program; Stephan, Penny, Daunizeau, Moran, & Friston, 2009), which gave this model a posterior probability of 87.95% relative to the other alternatives (Rescorla-Wagner with initial free parameters: 3.61%; Rescorla-Wagner without instruction parameter: 5.1%; Hybrid model: 3.35%). We therefore focus on this model in the main manuscript and used this model to generate regressors for use in fMRI analyses.

**Supplementary Results**

**Post-task ratings.**

We used ANOVAs to evaluate effects of Group and Cue on post-task affect ratings, and to measure effects of Group, Cue, and Phase for retrospective expectancy ratings (i.e. retrospective ratings of expected pain at the beginning and end of the task as a function of Cue). There were no differences in reported affect as a function of Group or Cue (all p’s > 0.09; see Supplementary Figure S15A). We observed a significant Group x Phase interaction on retrospective expectancy (F(1,36) = 4.386, p = .043) and a marginal Group x Cue x Phase interaction (F(1,36) = 3.635, p = .065). Post-hoc pairwise comparisons indicated that the Instructed Group reported differences in expected pain as a function of Cue at the beginning of the task (p_adjusted_ = .041), but not the end of the task (p > 0.9), whereas the Uninstructed Group did not report significant differences at any point (all p’s > 0.1). Group means are presented in Supplementary Figure S15.

**Model comparisons.**

*Pain ratings across temperatures.* Bayesian model comparison indicated that the best fitting model included fixed effects of Group, Temperature, Cue, and Phase, with all interactions and random intercepts and slopes (BF = 3.266*10^211, compared to a model with temperature only). The same model did not converge in the “lmer” function of *lme4* (Bates et al., 2015) unless we removed the covariance between random effects (i.e. specified that slopes and intercepts are uncorrelated using ||). Results of all approaches are reported in Table 1.

*Pain ratings on medium heat trials prior to reversal.* We first analyzed associations between predictors and pain on medium heat trials prior to the first instructed or experiential reversal (i.e. during acquisition). Bayesian model comparison using the function “loo_compare” (Vehtari et al., 2017, 2020) indicated that the best fitting model included fixed effects of Group, Cue, and Trial, with all interactions and random intercepts and slopes, although model comparison with bayesfactor_models indicated that the Trial factor was not necessary. Nonetheless we included Trial in our analyses to estimate a maximal model. The model with Trial included did not converge in the “lmer” function of *lme4* (Bates et al., 2015) unless we removed the covariance between random effects (i.e. specified that slopes and intercepts are uncorrelated using ||), and we used an optimizer in the “lme” function of *nlme* to achieve convergence (using the settings control = lmeControl(opt='optim')).

*Pain ratings on medium heat trials across the entire task.* We next evaluated predictions for medium pain across all trials, including reversals. Bayesian model comparison indicated that the best fitting model included fixed effects of Group, Cue, and Phase, with all interactions and random intercepts and slopes (BF = 1.459 *10^99, compared to a model with original cues only).

Although qqplots indicated that residuals were normally distributed with linear mixed models, and the various approaches produced similar results (see Table 3), posterior prediction plots for Bayesian models indicated that the pain rating data were skewed and were not well fit with a normal distribution. To validate the Bayesian model comparison, we performed posterior prediction to ensure models met assumptions using the “posterior_predict” function in the package *rstantools* (Gabry et al., 2020). We used the functions ppc_dens_overlay, ppc_hist, and ppc_stat from the *bayesplot* (Gabry et al., 2019; Gabry and Mahr, 2020) package to visualize the relationship between the observed data and the posterior. We therefore performed an additional model comparison using the best fitting model based on the normal distribution and comparing various families in brms. To facilitate comparisons across families, we normalized all responses by dividing by a factor of 10 (putting ratings on a 0-1 scale) and changed ratings of 0 to 0.1 (to facilitate log scaling). We compared the following families: normal; zero_inflated_beta, Beta, lognormal, skew_normal, student, and exGaussian. The function “loo_compare” in the package *‘loo’* (Vehtari et al., 2017, 2020) indicated that a Beta distribution provided the best fit for medium pain ratings. We therefore performed additional model comparisons to evaluate the best model based on the Beta distribution. Whereas assuming a normal distribution indicated that the best model included fixed effects of Group, Cue, and Phase, with all interactions and random intercepts and slopes, the Beta distribution supported the additional inclusion of a fixed factor for Trial.

Findings using the Beta family were consistent with models that assumed a normal distribution (i.e. the Cue, Phase, and Cue x Phase interactions all had >98% probability of being positive, and the Group x Cue x Phase interaction was marginal, with a 94.53% probability of being positive), although the practical significance of the Cue x Phase interaction was reduced (12% in ROPE). The additional main effect of time was associated with a 99.9% probability of being negative, such that pain decreased over time, but this finding was not practically significant (100% in ROPE). Notably, bayes factor comparisons using bayesfactor_models indicated that the model including time was not an improvement over the basic model identified using normal distribution (BF = 6*10^-83) and the Inclusion BF for Time was only 0.202, indicating that time explained very little variance in pain. Complete results are reported in Supplementary Table S2.

**Behavioral results restricted to fMRI participants (n=35).**

Whereas the main analyses focus on all 40 participants, we also analyzed temperature and cue effects on pain restricted to the 35 participants with useable fMRI data. We evaluated linear mixed models using the R packages LMER, NLME (to implement autoregression), and BRMS with weakly informative priors. See Methods for full details.

We first analyzed pain across all heat intensities. Consistent with results across all participants, Bayesian model comparison revealed that the best model included fixed effects of Temperature, Cue, Phase, and Group and all possible interactions, along with random intercepts and slopes. All models revealed significant effects of Temperature, Cue, Phase, Cue x Phase, and Temperature x Cue x Phase interactions across participants (see Supplementary Table S1). We also observed a significant Group x Temperature x Cue x Phase interaction; in contrast to the results across all participants, the Group x Cue x Phase interaction was non-significant or marginal (depending on the analysis approach) when restricted to fMRI participants (see Supplementary Table S1). Bayesian posterior estimates indicated that the effects of Temperature, Cue x Phase interactions, and Temperature x Cue x Phase interactions were practically significant with enough evidence to reject the null (<1% in ROPE), while the main effect of Phase supported the null (i.e. no influence of Phase; 99.8% in ROPE) despite being statistically significant. All other effects were of undecided significance (i.e. not enough evidence to accept or reject the null); complete results are reported in Supplementary Table S1.

**Validation of pain-related signature patterns.**

We tested whether the NPS and SIIPS could serve as pain-related brain signatures in the present dataset by testing the association between pain reports and signature pattern activation. We observed significant associations between subjective pain and both signatures across all trials (NPS: *B =* 4.25, SE = 0.54, p < .001; SIIPS: *B =* 440.32, SE = 60.1, p < .001), and both signatures were significantly associated with pain while controlling for heat intensity (NPS: *B =* 3.06, SE = 0.66, p < .001; SIIPS: *B =* 388.64, SE = 66.5, p < .001). There was a significant influence of Heat Intensity on the NPS while controlling for temperature (*B =* 3.53, SE = 1.32, p = .009), whereas there was no additional influence of Heat Intensity on SIIPS when accounting for pain (p > 0.2). These findings replicate previous work in other acute pain studies and validate the use of the NPS and SIIPS as pain-related brain signatures in the present study.

We also evaluated the association between neural signature activity and subjective pain on the critical medium heat trials, which were paired with predictive cues. NPS activation was significantly associated with pain ratings on medium heat trials (*B =* 5.69, SE = 1.46, p < .001) and effects did not differ between groups (all p’s > 0.1). Similarly, SIIPS activation was also significantly associated with pain on medium heat trials (*B =* 501.96, SE = 108.48, p < .001), and effects were similar across groups (all p’s > 0.1).

**Computational modeling and model comparison.**

*Model comparison for individual fits.* We fit several models to each individual’s pain ratings on medium heat trials and used Bayesian model comparison implemented with spm_BMS (Stephan et al., 2009) to compare between a) a Rescorla Wagner model that assumes initial expected value (1/0 for Instructed group, 0.5 for Uninstructed Group) and sets learning rate (ɑ) and instructed reversal parameters (⍴) as free parameters ; b) a Rescorla-Wagner model with a free parameter for initial expected value (EV), as well as free parameters for ɑ and ⍴; 3) a Rescorla-Wagner model with free parameters for ɑ and initial expected EV without a ⍴ parameter; and 4) a hybrid model of associability that includes parameters that describe how learning varies as a function of associability (kappa and eta) as well as an instructed reversal parameter (Li et al., 2011b; Atlas et al., 2019). Alpha values for the four respective models were as follows: [36.94; 1.5143; 2.14; 1.41], and exceedance probabilities were: [0.88; 0.04; 0.05; 0.03]. The first model (i.e. the Rescorla-Wagner model with fixed initial starting values and free parameters for ɑ and ⍴) was therefore determined to be the best fitting model and was used for statistical comparison as a function of group.

*Jack-knife model fitting.* We used fits to individual participants to determine the best-fitting model and perform group comparisons. We then used jack-knife estimation to generate group-level estimates that are less sensitive to individual noise and were used to generate fMRI regressors for each group. In particular, jack-knife estimation iteratively fits models to the entire group with one subject left out on each iteration, which provides a distribution of estimates that can be used for statistical comparison (Wu, 1986). See Atlas & Phelps (Atlas and Phelps, 2018) for complete details on jack-knife model fitting procedures.

Results of jack-knife estimation were consistent with results from individual subject fits. We observed a significant group difference in the learning rate ɑ (t(1,38) = 33.07, p < .001, CI = [.69, .78]), driven by higher ɑ values in the Uninstructed Group (*M =*.264, *SE = 0.028*) relative to the Instructed Group (*M =* 0, *SE* = 0), and a significant group difference in the instructed reversal parameter ⍴ (t(1,38) = -9.41, p < .001, CI = [-.32, -.21]) driven by higher ⍴ parameters in the Instructed Group (*M =* .875, *SE* = 0.003), relative to the Uninstructed Group (*M =* .139, *SE = 0.022*). Mean estimates for each group and each parameter were used to generate regressors that were used for fMRI analysis (see Figure 6B).

**Comparing instructed and feedback-driven signals within the Instructed Group.** Analyses across groups indicated that several regions showed differential associations with expected value between the two groups. For example, the rACC was positively associated with instructed EV in the Instructed Group and negatively associated with feedback-driven EV in the Uninstructed Group. The instructed EV timecourse identifies responses that update immediately with instruction and do not vary as a function of reinforcement between the instructed reversals, mimicking the timecourse of pain, to which EV signals were fit. However, it is possible that individual brain regions update with different timecourses, i.e. as a function of experience even if they do not lead to changes in pain, consistent with the dissociations we observed in our two mediation analyses (i.e. Figure 4 and Figure 5) and previous work on appetitive and aversive learning (Li et al., 2011a; Atlas et al., 2016). We therefore searched for correlates of instructed and feedback-driven EV signals within Instructed Group participants and contrasted the two sources of EV to determine whether any regions responded preferentially to one type of EV signal over the other. When we tested effects within *a priori* value-processing regions of interest, we observed significant associations with Instructed EV, controlling for uninstructed EV, in the bilateral striatum (see Table 4), although the difference between instructed and uninstructed EV was not significant in either ROI. There were no significant associations for the amygdala or VMPFC ROIs (see Table 4). Whole brain FDR-correction revealed significant positive associations with Instructed EV in the midbrain near the substantia nigra and negative associations in right M1/ precentral gyrus (M1; see Figure S14). However no regions survived whole brain correction for Uninstructed EV or the difference between Instructed and Uninstructed EV, and we did not observe significant associations with EV within *a priori* pain modulatory networks (see Table S10). Exploratory whole brain uncorrected results are presented in Supplementary Figure S15 and Supplementary Table S11.

Supplementary Figures

Supplementary Figure S1. A priori regions of interest.

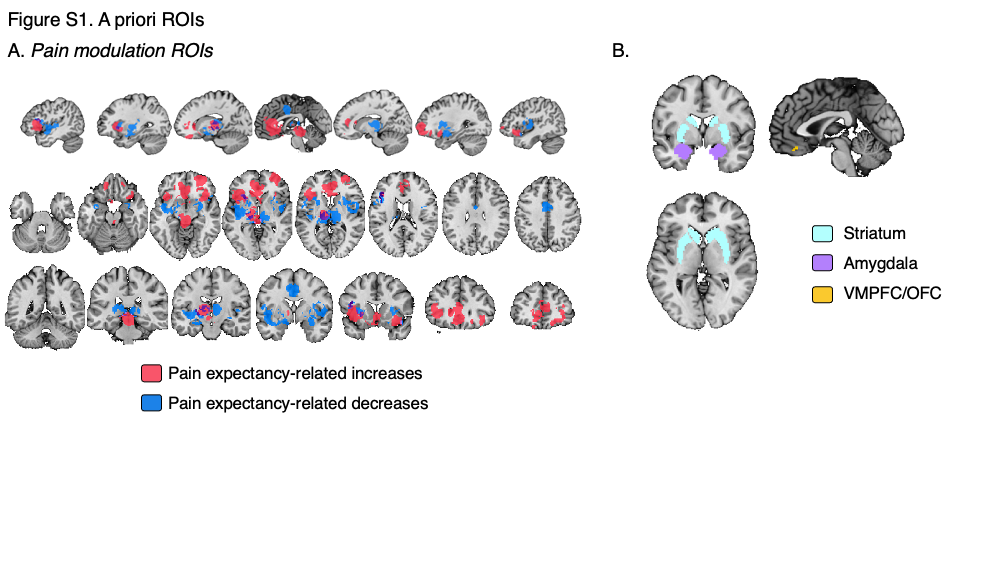

Figure S1. A priori regions of interest. We searched within two sets of a priori regions of interest (ROIs). A) Pain modulation ROIs were identified through a meta-analysis of placebo analgesia and expectancy-based pain modulation (Atlas and Wager, 2014). We created a mask combined across modulatory regions whose responses to noxious stimuli increase with expectations for reduced pain (red) and regions whose responses to noxious stimuli decrease with expectations for reduced pain (blue). B) We also examined responses within value-related ROIs (right), including the bilateral striatum (cyan), bilateral amygdala (violet), and the VMPFC/OFC (yellow). See Methods for details of ROI identification and ROI-wise analyses.

Supplementary Figure S2. Temperature effects on pain.

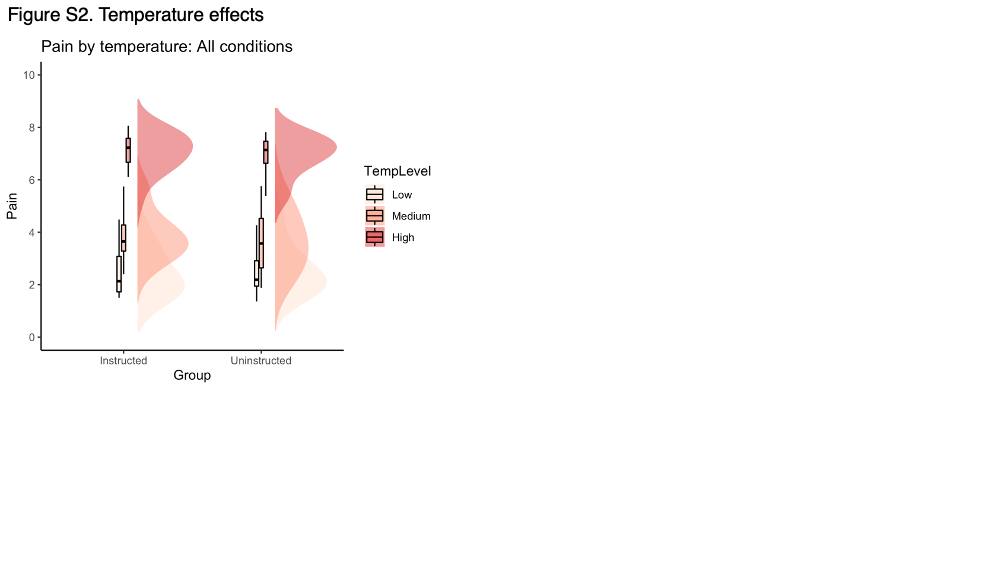

Figure S2. Temperature effects on pain. Associations between heat intensity and pain were similar across groups. See Table 1 and Supplementary Table S1 for complete results.

Supplementary Figure S3. Cue and Phase effects for individual participants

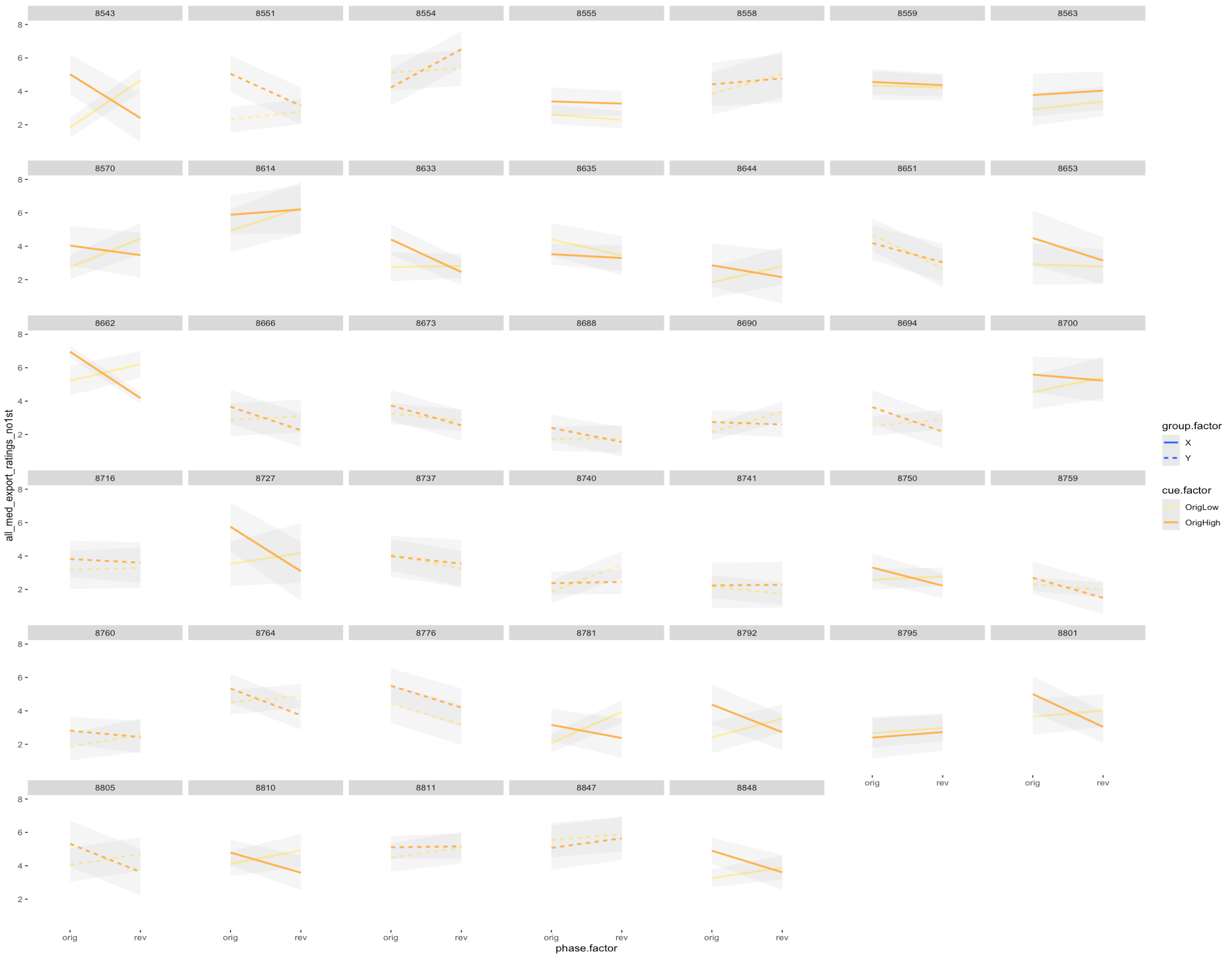

Figure S3. Cue and Phase effects for individual participants. This figure depicts effects of Cue (original low = yellow; original high =orange) and Phase (original = “orig”, reversed = “rev”) on subjective pain on medium heat trials for each participant. Instructed Group participants are displayed with solid lines and Uninstructed Participants are displayed in dashed lines.

Figure S4. Current contingency mediation: Effects of cue and phase within pain modulatory network.

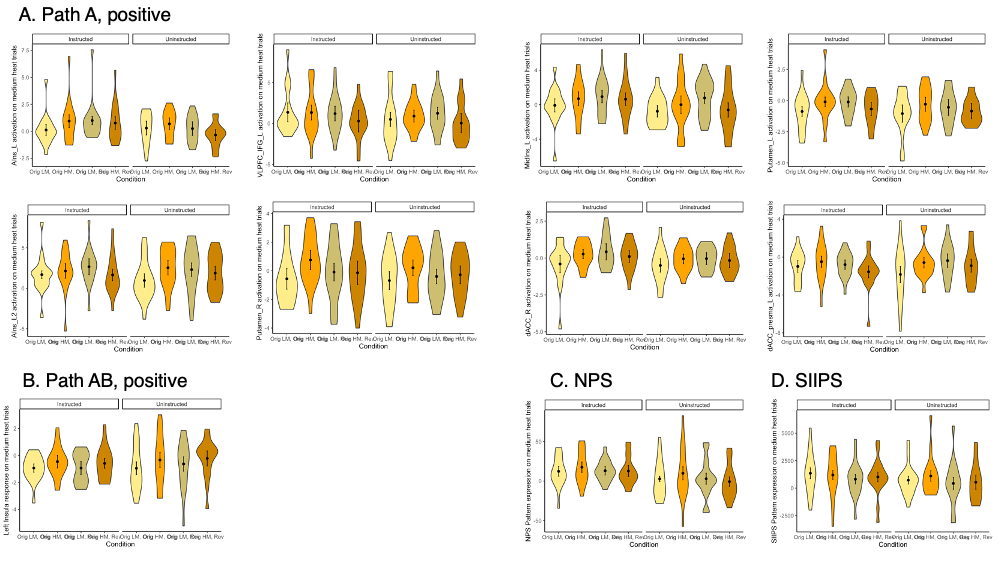

Figure S4. Current contingency mediation: Effects of cue and phase within pain modulatory network. This figure depicts extracted responses within pain modulatory regions for Path a and mediation of current contingency effects on medium heat pain. A) Responses in pain modulatory clusters identified in analyses of Path a, or regions identified as showing higher activation in response to current high pain cues relative to current low pain cues. We extracted responses and averaged as a function of Group (Instructed vs Uninstructed), Cue (Original low = “Orig LM”; Original high = “Orig HM”), Phase (Left two lighter bars = Original contingencies; Right two darker bars = reversed contingencies). This confirms that these regions update responses as contingencies change and do so similarly across both groups. B) Responses within the left insula mediator region. C. Pattern expression within the neurologic signature pattern (NPS; (Wager et al., 2013)). D) Pattern expression within the stimulus-intensity independent pain signature pattern (SIIPS; (Woo et al., 2017)).

Figure S5. Current contingency mediation: Whole brain FDR-correction.

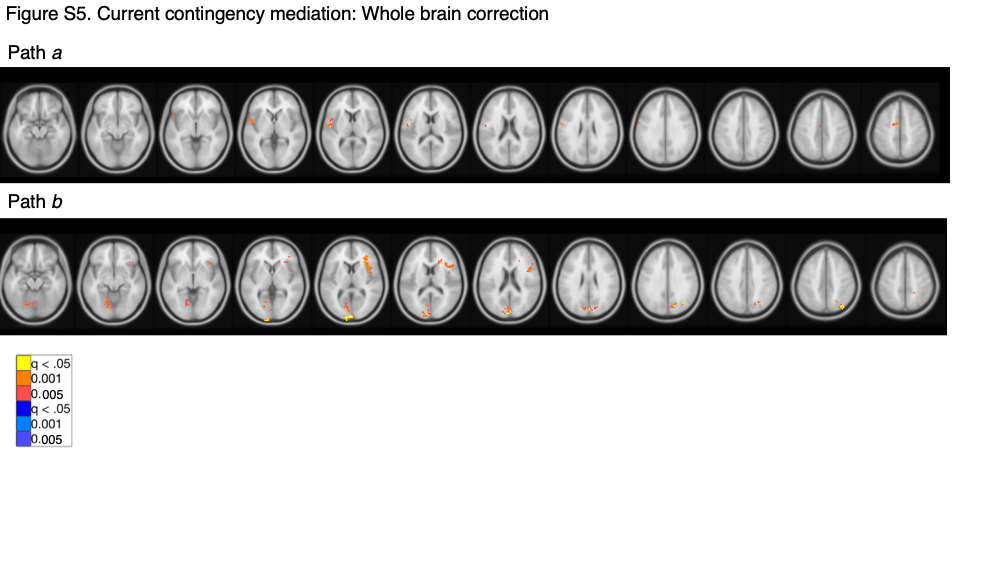

Figure S5. Current contingency mediation: Whole-brain FDR correction. Results are reported in Supplementary Table S4

Figure S6. Current contingency mediation: Whole brain uncorrected results

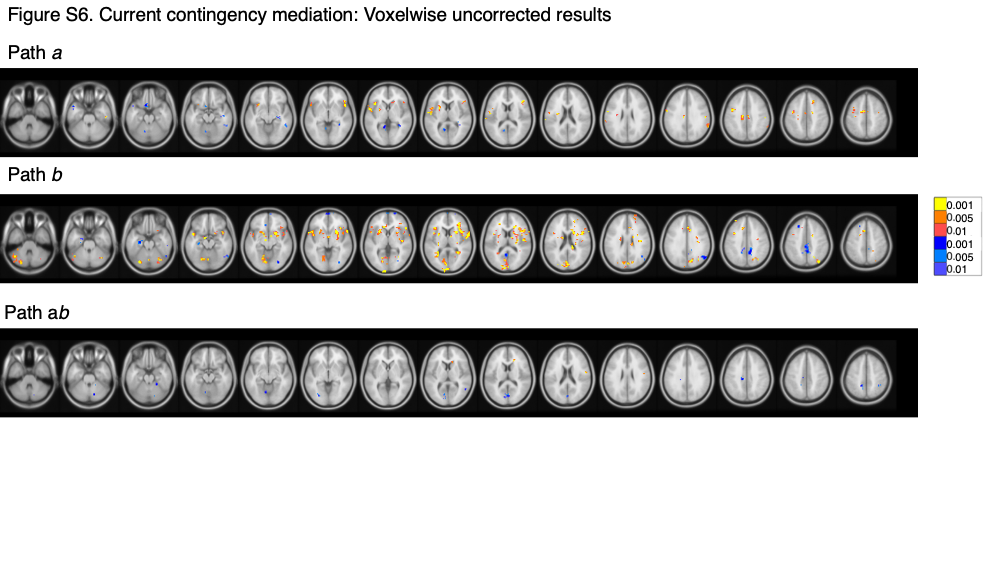

Figure S6. Current contingency mediation: Whole-brain uncorrected. Results are reported in Supplementary Table S5.

Figure S7. Current contingency mediation including Group as a moderator: Voxelwise uncorrected results.

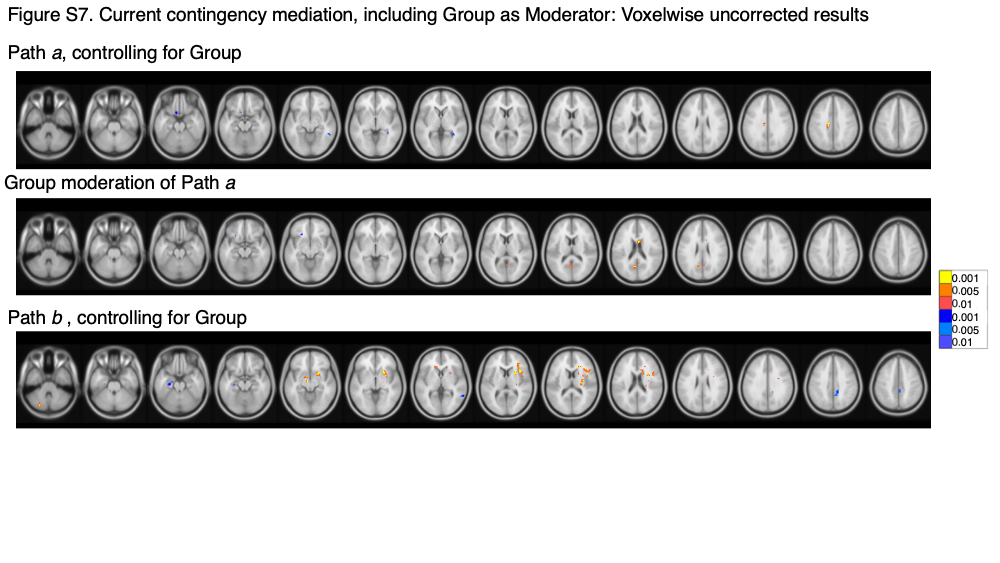

Figure S7. Current contingency mediation including Group as a moderator: Whole-brain unccorrected. Results are reported in Supplementary Table S5.

Figure S8. Original contingency mediation: Effects of cue and phase within pain modulatory network.

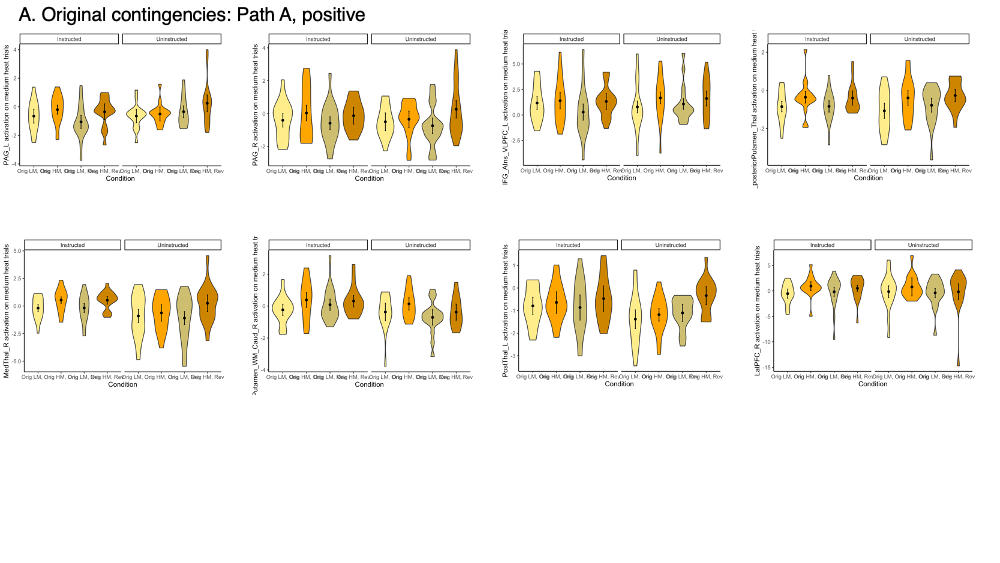

Figure S8. Original contingency mediation: Effects of cue and phase within pain modulatory network. This figure depicts extracted responses in pain modulatory clusters identified in analyses of Path a of the mediation based on original contingencies, i.e. regions identified as showing higher activation in response to original high pain cues relative to original low pain cues, regardless of phase. We extracted responses and averaged as a function of Group (Instructed vs Uninstructed), Cue (Original low = “Orig LM”; Original high = “Orig HM”), Phase (Left two lighter bars = Original contingencies; Right two darker bars = reversed contingencies). This confirms that these regions maintain initial responses despite contingency reversals and that responses are similar across both groups.

Figure S9. Original contingency mediation: Whole brain correction.

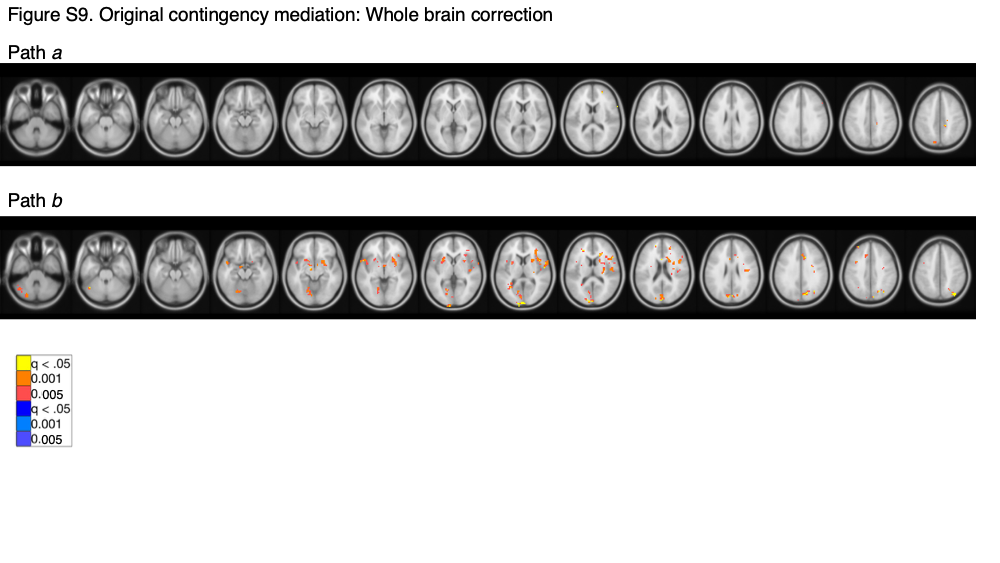

Figure S9. Original contingency mediation: Whole-brain FDR correction. Results are reported in Supplementary Table S6.

Figure S10. Original contingency mediation: Whole brain uncorrected results.
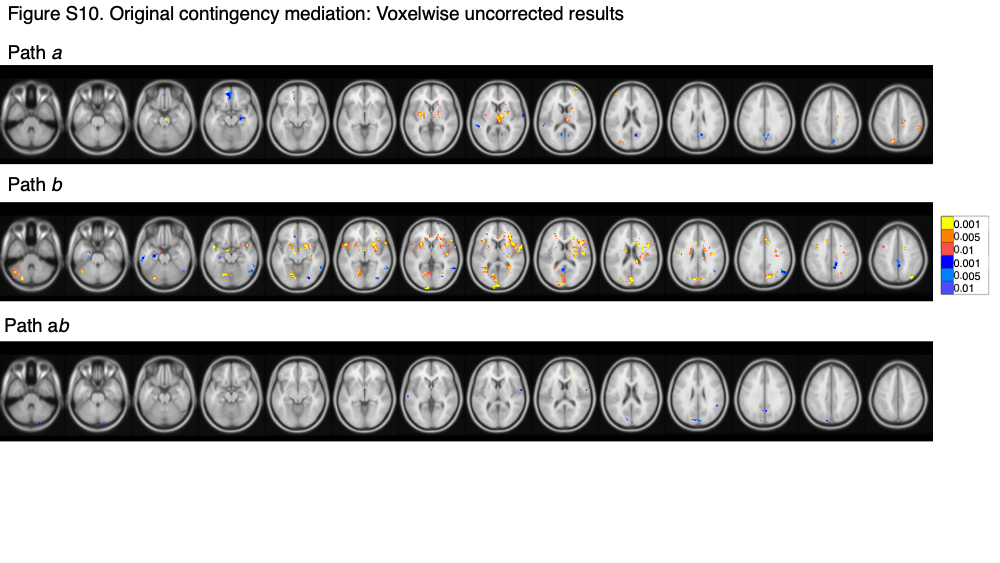

Figure S10. Original contingency mediation: Whole-brain uncorrected results. Results are reported in Supplementary Table S7.

Figure S11. Original contingency mediation, including Group as moderator: Voxelwise uncorrected results.

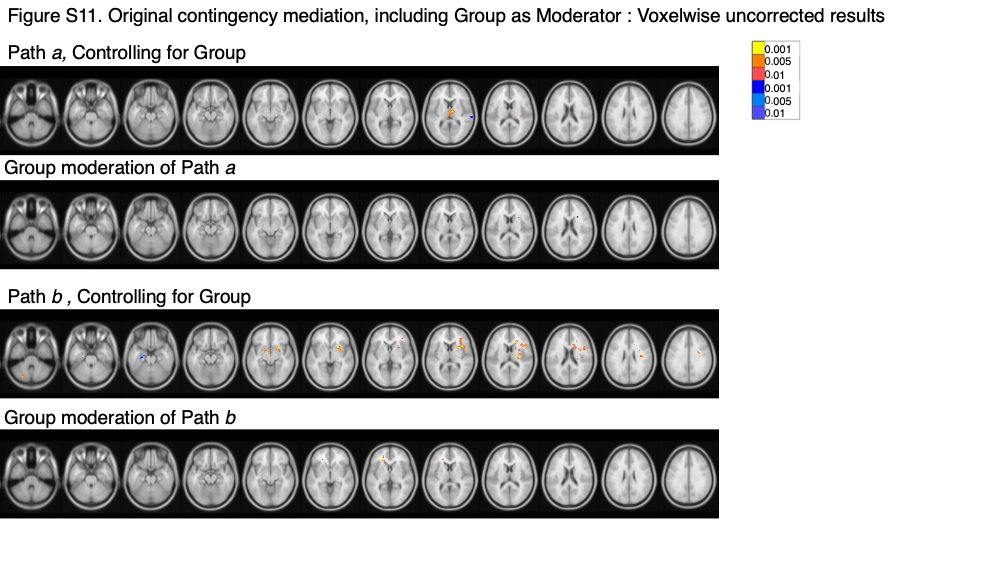

Figure S11. Original contingency mediation, including Group as a moderator: Whole-brain uncorrected results. Results are reported in Supplementary Table S7.

Figure S12. Expected Value: Whole brain correction.

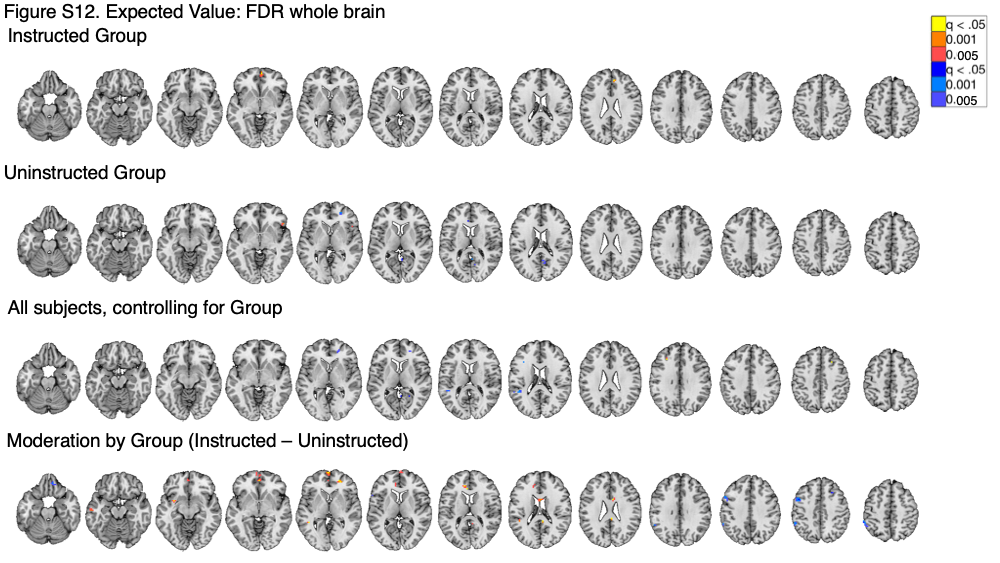

Figure S12. Expected value: Whole-brain correction. Results are reported in Supplementary Table S8.

Figure S13. Expected Value: Whole brain uncorrected results.

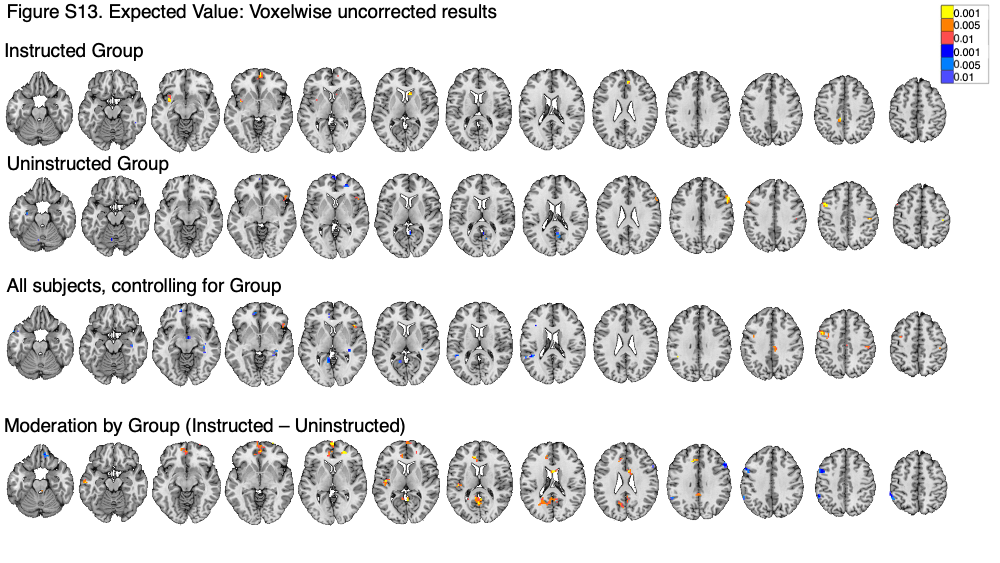

Figure S13. Expected value: Whole-brain uncorrected results. Results are reported in Supplementary Table S9.

Figure S14. Instructed vs Feedback driven Expected Value within Instructed Group: FDR whole brain

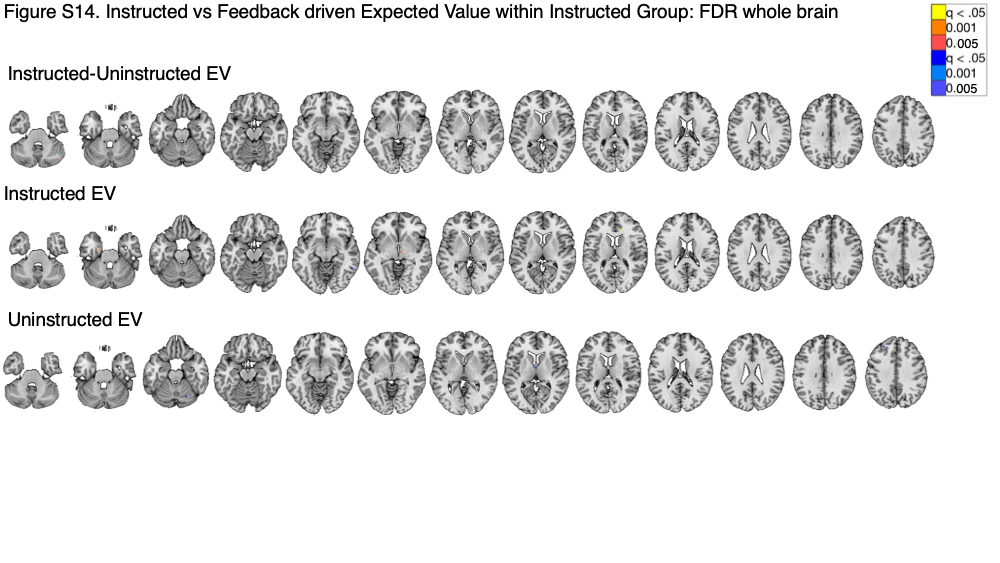

Figure S14. Instructed vs Feedback driven Expected Value within Instructed Group: Whole-brain corrected results. Results are reported in Supplementary Table S10.

Figure S15. Instructed vs Feedback driven Expected Value within Instructed Group: Uncorrected whole brain results.

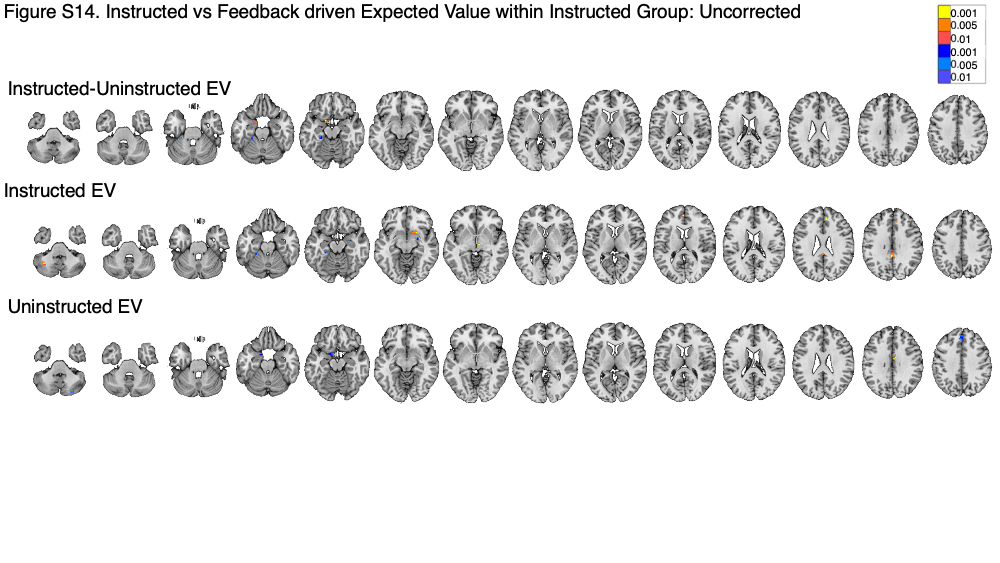

Figure S15. Instructed vs Feedback driven Expected Value within Instructed Group: Whole-brain uncorrected results. Results are reported in Supplementary Table S11.

Figure S16. Retrospective ratings.
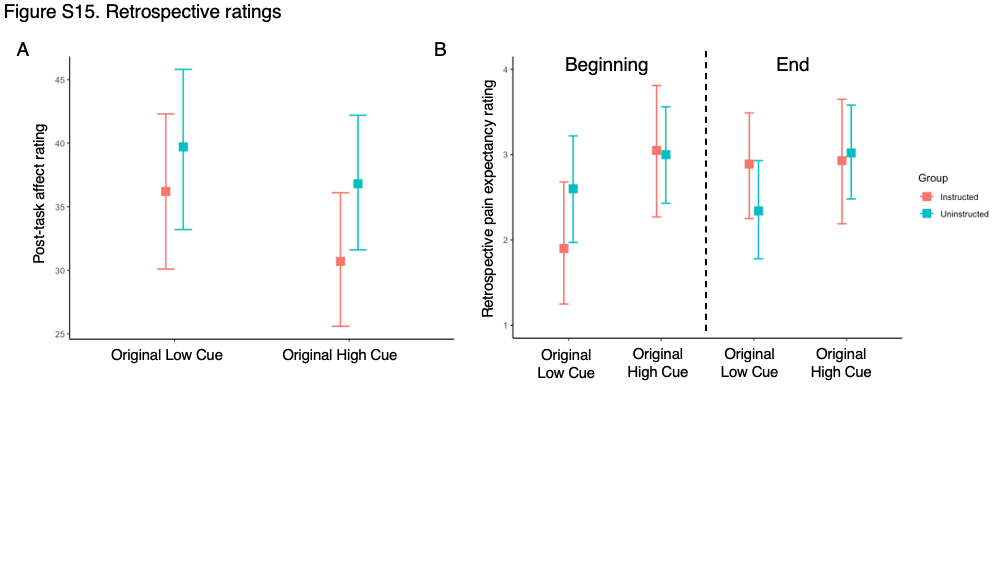

Figure S16. Retrospective ratings. Following the task, participants provided retrospective ratings of affect (A) and expected pain at the beginning and end of the task (B) as a function of Cue.

Supplementary Tables

Table S1. Heat intensity effects on pain in fMRI participants (n = 35).^a^

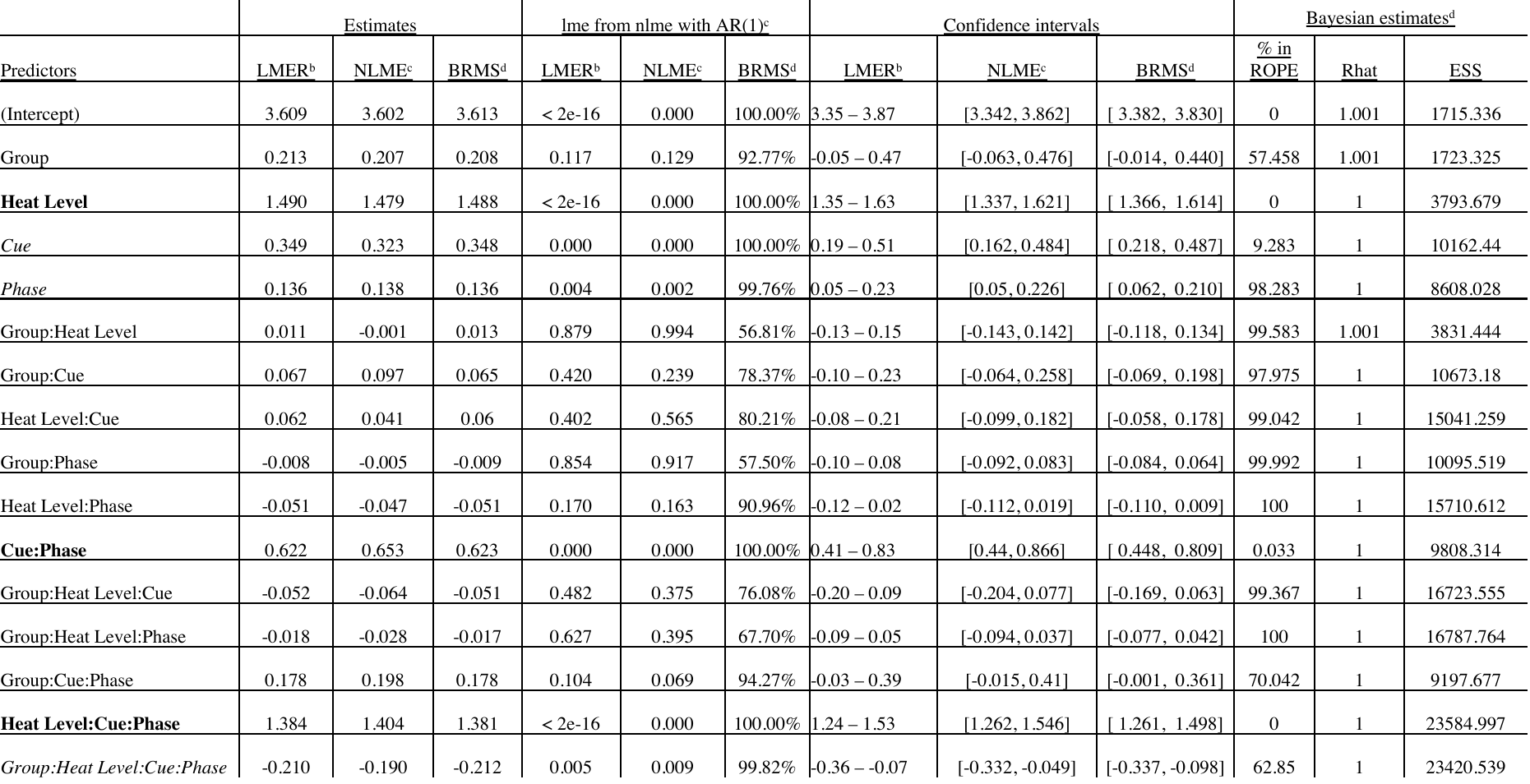

^a^. This table presents results of a linear mixed model predicting subjective pain as a function of Heat Level (High vs Medium vs Low), Group (Instructed vs Uninstructed), Cue (Original High vs Original Low), and Phase (Original vs Reversed) in subjects with useable fMRI data (n = 35). Model specification is identical to Table 1. Trials that were omitted from fMRI analyses (e.g. due to head motion or scanner artifacts) were also omitted for consistency with neuroimaging analyses.

^b^. Estimates based on a linear mixed effects model implemented in the “lmer” function of lme4 (Bates et al., 2015) using the following code: lmer(Pain~group*templevels*cue*phase+(1+templevels+cue*phase||subject)).

^c^. Estimates based on a linear mixed effects model implemented in the “lme” function of nlme (Pinheiro et al., 2021) including autoregression using the following code: lme(Pain~group*templevels*cue*phase, random=~1+templevels+cue*phase|subject, correlation=corAR1(), na.action=na.exclude).

^d^. Estimates based on Bayesian model linear mixed models using the “brms” function (Bürkner, 2017) using the following code: brm(Pain~group*templevels*cue*phase+(1+templevels+cue*phase|subject,prior=set_prior("normal(0,2.5)", class="b"), save_all_pars=TRUE, silent=TRUE, refresh=0, iter = 4000, warmup = 1000). Posterior estimates and the Region of Partial Equivalence were obtained using the “describe_posterior” function from the package BayesTestR (Makowski et al., 2019a) and interpreted as in (Makowski et al., 2019b). The Region of Partial Equivalence (ROPE) was defined as [-0.176, 0.176].

Table S2. Effects on medium heat pain within fMRI participants (n = 35).^a^

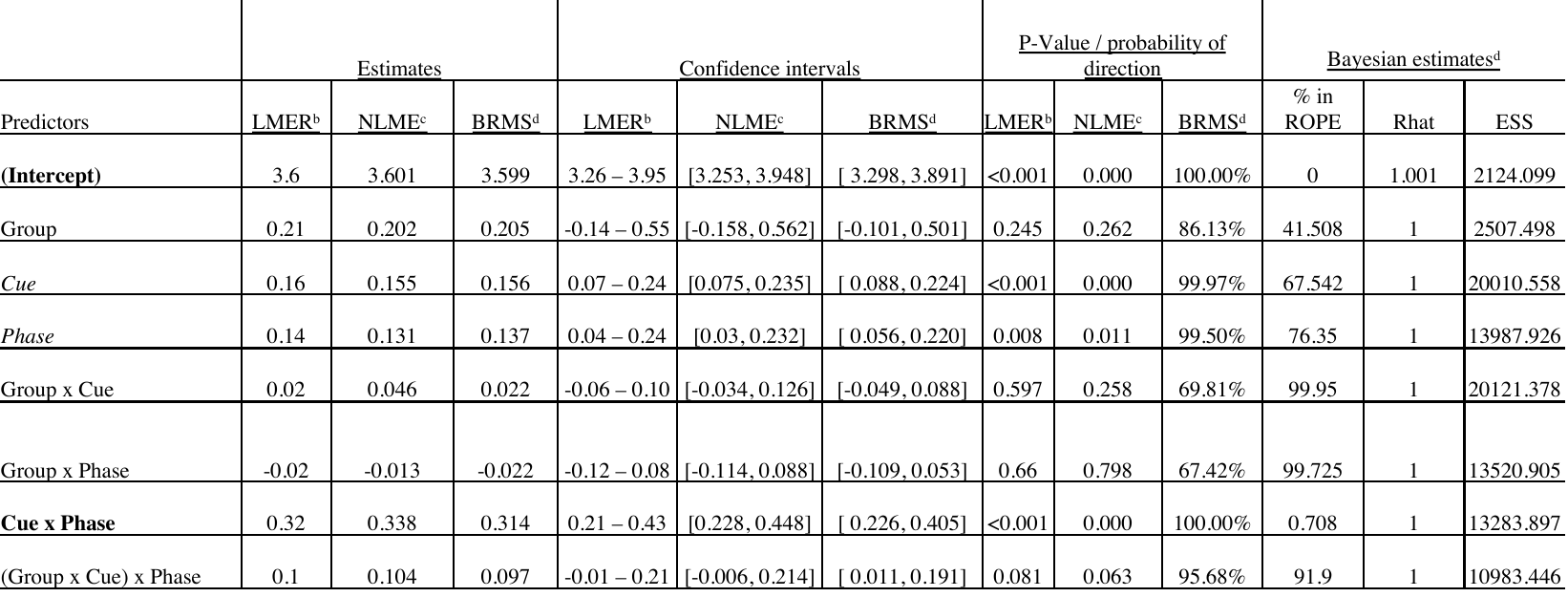
 ^a^. This table presents results of a linear mixed model predicting subjective pain on medium heat trials as a function of Group (Instructed vs Uninstructed), Cue (Original High vs Original Low), and Phase (Original vs Reversed) across all fMRI participants (n = 35). See Table 1 for additional information about model specification and presentation.

^b^. Estimates based on a linear mixed effects model implemented in the “lmer” function of lme4 (Bates et al., 2015) using the following code: lmer(Pain_Medium_~Group*Cue*Phase+(1+ Cue*Phase|Subject)).

^c^. Estimates based on a linear mixed effects model implemented in the “lme” function of nlme (Pinheiro et al., 2021) including autoregression using the following code: lme(Pain~Group *Cue*Phase, random=~1+Cue*Phase|Subject, correlation=corAR1(), na.action=na.exclude).

^d^. Estimates based on Bayesian model linear mixed models using the “brms” function (Bürkner, 2017) using the following code: brm(Pain~Group *Cue*Phase+(1+Cue*Phase|Subject,prior=set_prior("normal(0,2.5)", class="b"), save_all_pars=TRUE, silent=TRUE, refresh=0, iter = 4000, warmup = 1000). Posterior estimates and the Region of Partial Equivalence were obtained using the “describe_posterior” function from the package BayesTestR (Makowski et al., 2019a) and interpreted as in (Makowski et al., 2019b). The Region of Partial Equivalence (ROPE) was defined as [-0.175, 0.175].

Table S3. Bayesian multilevel model evaluating effects of Group, Cue, Phase, and Trial on medium heat pain using Beta family.^a^

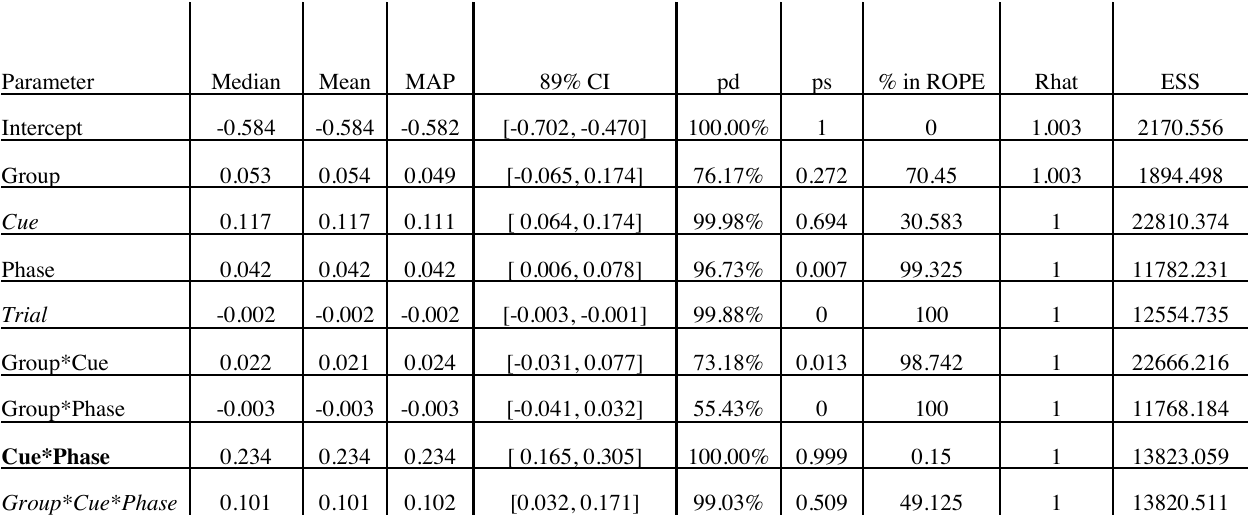

^a^. Estimates based on Bayesian model linear mixed models using the “brms” function (Bürkner, 2017) using the following code: brm(Pain~group*cue*phase+trial+(1+cue*phase|subject,prior=set_prior("normal(0,2.5)", class="b"), family = Beta(), save_all_pars=TRUE, silent=TRUE, refresh=0, iter = 4000, warmup = 1000). Posterior estimates and the Region of Partial Equivalence were obtained using the “describe_posterior” function from the package BayesTestR (Makowski et al., 2019a) and interpreted as in (Makowski et al., 2019b). The Region of Partial Equivalence (ROPE) was defined as [-0.236, 0.236].

Table S4. Mediation of current cue contingencies: Small-volumes corrected results.^e^

| **Correction** | **Effect** | **Anatomical label** | **x** | **y** | **z** | **# of voxels** | **Volume (mm^3^)** |
| --- | --- | --- | --- | --- | --- | --- | --- |
| Pain modulatory network | *Path a, positive* | L Insula Lobe | -32 | 16 | -8 | 9 | 243 |
|  |  | L IFG p. Triangularis (VLPFC) | -40 | 32 | -2 | 5 | 135 |
|  |  | L Rolandic Operculum / Middle insula | -50 | -2 | 4 | 22 | 594 |
|  |  | L Putamen | -26 | 8 | 8 | 17 | 459 |
|  |  | L Insula Lobe | -38 | 20 | 2 | 12 | 324 |
|  |  | R MCC / dACC | 8 | -2 | 32 | 5 | 135 |
|  |  | L dACC / Posterior-Medial Frontal | -8 | -4 | 50 | 14 | 378 |
|  | *Path a, negative* | *no voxels survive* |  |  |  |  |  |
|  | *Path b, positive* | L Hippocampus | -14 | -4 | -16 | 4 | 108 |
|  |  | L Superior Temporal Gyrus / Anterior insula | -44 | 4 | -14 | 19 | 513 |
|  |  | R Putamen / Amygdala | 26 | 8 | -4 | 69 | 1863 |
|  |  | L Ventral Striatum | -14 | 4 | -10 | 7 | 189 |
|  |  | L Putamen | -26 | 14 | 2 | 38 | 1026 |
|  |  | R IFG p. Orbitalis / Anterior insula | 38 | 26 | -8 | 28 | 756 |
|  |  | L Anterior Insula Lobe | -40 | 20 | 4 | 44 | 1188 |
|  |  | L pregenual ACC | -10 | 28 | 2 | 19 | 513 |
|  |  | R Rolandic Operculum / Area OP4 [PV] (dorsal posterior insula) | 52 | -8 | 10 | 25 | 675 |
|  |  | R IFG p. Opercularis / middle insula | 40 | 10 | 8 | 28 | 756 |
|  |  | L IFG p. Opercularis / anterior insula | -40 | 8 | 20 | 9 | 243 |
|  | *Path b, negative* | *no voxels survive* |  |  |  |  |  |
|  | *Path ab, positive* | L middle insula | -34 | -8 | 2 | 5 | 135 |
|  | *Path ab, negative* | *no voxels survive* |  |  |  |  |  |
| Whole brain correction | *Path a, Positive* | L Rolandic Operculum | -50 | 2 | 8 | 82 | 2214 |
|  |  | DLPFC / dACC | -20 | -8 | 62 | 95 | 2565 |
|  | *Path a, Negative* | *no voxels survive* |  |  |  |  |  |
|  | *Path b, Positive* | L Cuneus / Area hOc2 [V2] | -2 | -80 | 16 | 372 | 10044 |
|  |  | R IFG p. Orbitalis / Anterior insula | 38 | 26 | -8 | 18 | 486 |
|  |  | R IFG p. Triangularis / Anterior insula | 38 | 20 | 10 | 162 | 4374 |
|  |  | Area 2 | 22 | -44 | 58 | 32 | 864 |
|  |  | L Precuneus / Area 7A (SPL) | -8 | -70 | 58 | 12 | 324 |
|  |  | L Postcentral Gyrus / Area 4a | -34 | -28 | 68 | 32 | 864 |
|  |  | RPrecentral Gyrus | 22 | -28 | 64 | 14 | 378 |
|  |  | R Superior Parietal Lobule / Area 5L (SPL) | 20 | -50 | 74 | 8 | 216 |
|  |  | L Paracentral Lobule / Area 4a | -10 | -28 | 74 | 6 | 162 |
|  | *Path b, Negative* | *no voxels survive* |  |  |  |  |  |
|  | *Path ab, Positive* | *no voxels survive* |  |  |  |  |  |
|  | *Path ab, Negative* | *no voxels survive* |  |  |  |  |  |

^e^. This table presents small-volume corrected results from a voxelwise multilevel mediation analysis evaluating the effects of cues on subjective pain based on current contingencies (i.e. including reversals), regardless of Group. Top rows are FDR-corrected within pain modulatory networks (see Supplementary Figure S1) and bottom rows are whole-brain corrected. Both analyses are FDR-corrected at q < .05. See Methods for additional details.

Table S5. Mediation of current cue contingencies: Uncorrected results.^f^

| **Analysis** | **Effect** | **Anatomical label** | **x** | **y** | **z** | **# of voxels** | **Volume (mm^3^)** |
| --- | --- | --- | --- | --- | --- | --- | --- |
| All participants, regardless of Group | Path a, positive | L Cerebelum VIII / Lobule VIIb (Hem) | -34 | -62 | -56 | 18 | 486 |
|  |  | R Inferior Temporal Gyrus | 50 | -20 | -28 | 6 | 162 |
|  |  | L Anterior Insula Lobe | -32 | 16 | -10 | 13 | 351 |
|  |  | R IFG p. Triangularis / Area 45 | 50 | 22 | 2 | 65 | 1755 |
|  |  | L Rolandic Operculum / Middle insula | -50 | 2 | 8 | 104 | 2808 |
|  |  | R Caudate Nucleus | 14 | 22 | -2 | 9 | 243 |
|  |  | L Putamen | -26 | 4 | 4 | 23 | 621 |
|  |  | L Anterior Insula Lobe | -38 | 20 | 2 | 12 | 324 |
|  |  | L Superior Temporal Gyrus / Area PFop (IPL) | -64 | -28 | 20 | 10 | 270 |
|  |  | R Postcentral Gyrus / Area PFt (IPL) | 56 | -20 | 32 | 10 | 270 |
|  |  | R SupraMarginal Gyrus / Area PFm (IPL) | 62 | -46 | 34 | 22 | 594 |
|  |  | L Superior Frontal Gyrus | -20 | -8 | 56 | 206 | 5562 |
|  |  | R MCC | 4 | -20 | 38 | 11 | 297 |
|  |  | R Superior Frontal Gyrus | 16 | -8 | 44 | 5 | 135 |
|  |  | R Superior Frontal Gyrus (DMPFC) | 20 | 26 | 44 | 11 | 297 |
|  |  | L Middle Frontal Gyrus | -26 | -20 | 46 | 14 | 378 |
|  |  | RPrecentral Gyrus | 32 | -8 | 52 | 12 | 324 |
|  |  | R Postcentral Gyrus | 26 | -40 | 74 | 12 | 324 |
|  | Path a, negative | L Medial Temporal Pole | -46 | 10 | -26 | 14 | 378 |
|  |  | L Cerebelum VI / Lobule VI (Hem) | -14 | -62 | -22 | 13 | 351 |
|  |  | L Rectal Gyrus / Area Fo2 / subgenual ACC | -8 | 14 | -22 | 28 | 756 |
|  |  | R Inferior Temporal Gyrus | 52 | -44 | -14 | 19 | 513 |
|  |  | R Hippocampus / CA2 | 34 | -20 | -14 | 26 | 702 |
|  |  | L Calcarine Gyrus / Area hOc1 [V1] | -10 | -56 | 4 | 42 | 1134 |
|  | Path b, positive | R Cerebelum VIII / Lobule VIIb (Hem) | 34 | -64 | -52 | 20 | 540 |
|  |  | L Cerebelum VIII / Lobule VIIIa (Hem) | -32 | -56 | -50 | 35 | 945 |
|  |  | L Cerebelum Crus 2 / Lobule VIIa crusII (Hem) | -28 | -80 | -38 | 45 | 1215 |
|  |  | L Cerebelum VII / Lobule VI (Hem) | -32 | -38 | -40 | 22 | 594 |
|  |  | L Cerebelum Crus 1 / Lobule VIIa crusI (Hem) | -44 | -62 | -32 | 59 | 1593 |
|  |  | R Cerebelum VI / Lobule VIIa crusI (Hem) | 32 | -70 | -26 | 40 | 1080 |
|  |  | L Calcarine Gyrus / Area hOc2 [V2] | -2 | -80 | 14 | 509 | 13743 |
|  |  | R Insula Lobe | 38 | 10 | 8 | 694 | 18738 |
|  |  | L Putamen | -20 | 8 | -8 | 122 | 3294 |
|  |  | L Insula Lobe | -44 | 14 | -4 | 106 | 2862 |
|  |  | R Insula Lobe | 46 | 8 | -10 | 35 | 945 |
|  |  | R Hypothalamus (ventromedial) | 4 | -2 | -8 | 15 | 405 |
|  |  | R Calcarine Gyrus / Area hOc1 [V1] | 28 | -62 | 4 | 11 | 297 |
|  |  | L Putamen | -28 | -22 | 8 | 12 | 324 |
|  |  | L IFG p. Opercularis | -40 | 8 | 14 | 23 | 621 |
|  |  | Thal: Temporal | 10 | -4 | 10 | 15 | 405 |
|  |  | L Caudate Nucleus | -16 | -8 | 20 | 17 | 459 |
|  |  | R Middle Frontal Gyrus | 26 | 52 | 22 | 34 | 918 |
|  |  | L Middle Occipital Gyrus | -28 | -82 | 22 | 5 | 135 |
|  |  | L rdACC | -14 | 22 | 38 | 22 | 594 |
|  |  | L DLPFC | -32 | 46 | 38 | 6 | 162 |
|  |  | L Precentral Gyrus | -38 | 4 | 40 | 12 | 324 |
|  |  | Area 2 | 22 | -44 | 58 | 38 | 1026 |
|  |  | L Posterior-Medial Frontal | -2 | 16 | 56 | 66 | 1782 |
|  |  | L Superior Frontal Gyrus | -20 | 2 | 52 | 10 | 270 |
|  |  | L Precuneus / Area 7A (SPL) | -8 | -70 | 58 | 12 | 324 |
|  |  | R Superior Parietal Lobule / Area 7P (SPL) | 22 | -76 | 58 | 4 | 108 |
|  |  | L Postcentral Gyrus / Area 4a | -34 | -32 | 68 | 43 | 1161 |
|  |  | L Postcentral Gyrus / Area 3a | -20 | -32 | 58 | 5 | 135 |
|  |  | R Superior Frontal Gyrus | 16 | -16 | 62 | 9 | 243 |
|  |  | RPrecentral Gyrus | 22 | -28 | 68 | 26 | 702 |
|  |  | RPrecentral Gyrus | 34 | -14 | 64 | 7 | 189 |
|  |  | R Superior Parietal Lobule / Area 5L (SPL) | 20 | -50 | 74 | 14 | 378 |
|  |  | L Paracentral Lobule / Area 4a | -10 | -28 | 74 | 8 | 216 |
|  |  | R Posterior-Medial Frontal | 10 | -2 | 74 | 10 | 270 |
|  |  | L Superior Parietal Lobule / Area 5L (SPL) | -20 | -46 | 74 | 8 | 216 |
|  | Path b, Negative | L Hippocampus / DG | -26 | -16 | -22 | 45 | 1215 |
|  |  | R Inferior Temporal Gyrus | 52 | -58 | -20 | 22 | 594 |
|  |  | R Inferior Temporal Gyrus | 58 | -32 | -20 | 8 | 216 |
|  |  | R Inferior Occipital Gyrus / Area hOc4v [V4(v)] | 32 | -80 | -8 | 14 | 378 |
|  |  | L Mid Orbital Gyrus / Area s32 | -8 | 34 | -14 | 10 | 270 |
|  |  | L Mid Orbital Gyrus / Area Fp1 | -2 | 68 | -4 | 28 | 756 |
|  |  | R Superior Frontal Gyrus / Area Fp1 | 20 | 70 | 2 | 15 | 405 |
|  |  | L Precuneus | -4 | -56 | 14 | 18 | 486 |
|  |  | R Angular Gyrus / Area PGp (IPL) | 52 | -62 | 32 | 42 | 1134 |
|  |  | R Precuneus | 4 | -44 | 40 | 122 | 3294 |
|  |  | L Superior Frontal Gyrus | -22 | 28 | 46 | 13 | 351 |
|  | Path ab, Positive | R Cerebelum VIII / Lobule VIIa crusII (Hem) | 32 | -64 | -46 | 6 | 162 |
|  |  | R Postcentral Gyrus / Area 3a | 56 | -8 | 22 | 8 | 216 |
|  |  | L Middle Frontal Gyrus | -28 | 40 | 46 | 6 | 162 |
|  | Path ab, Negative | L Hippocampus / DG | -26 | -16 | -22 | 45 | 1215 |
|  |  | R Inferior Temporal Gyrus | 52 | -58 | -20 | 22 | 594 |
|  |  | R Inferior Temporal Gyrus | 58 | -32 | -20 | 8 | 216 |
|  |  | R Inferior Occipital Gyrus / Area hOc4v [V4(v)] | 32 | -80 | -8 | 14 | 378 |
|  |  | L Mid Orbital Gyrus / Area s32 | -8 | 34 | -14 | 10 | 270 |
|  |  | L Mid Orbital Gyrus / Area Fp1 | -2 | 68 | -4 | 28 | 756 |
|  |  | R Superior Frontal Gyrus / Area Fp1 | 20 | 70 | 2 | 15 | 405 |
|  |  | L Precuneus | -4 | -56 | 14 | 18 | 486 |
|  |  | R Angular Gyrus / Area PGp (IPL) | 52 | -62 | 32 | 42 | 1134 |
|  |  | R Precuneus | 4 | -44 | 40 | 122 | 3294 |
|  |  | L Superior Frontal Gyrus | -22 | 28 | 46 | 13 | 351 |
| Moderation analysis, controlling for Group | Path a, positive | L Middle / posterior cingulate cortex | -14 | -20 | 34 | 20 | 540 |
|  | Path a, negative | L Rectal Gyrus / Area Fo2 / subgenual ACC | -10 | 14 | -22 | 15 | 405 |
|  |  | R Inferior Temporal Gyrus | 52 | -44 | -14 | 9 | 243 |
|  | Path b, positive | L Cerebelum Crus 1 / Lobule VIIa crusI (Hem) | -26 | -76 | -34 | 21 | 567 |
|  |  | L amygdala, contiguous with midbrain near substantia nigra | -10 | -8 | -14 | 24 | 648 |
|  |  | R Putamen | 26 | 10 | 2 | 96 | 2592 |
|  |  | R IFG p. Opercularis | 40 | 16 | 10 | 96 | 2592 |
|  | Path b, Negative | L ParaHippocampal Gyrus / Subiculum | -28 | -20 | -22 | 26 | 702 |
|  |  | R Middle Temporal Gyrus | 58 | -52 | 2 | 10 | 270 |
|  |  | R MCC | 10 | -40 | 38 | 33 | 891 |
|  | Path ab, Positive | *no voxels survive* |  |  |  |  |  |
|  | Path ab, Negative | *no voxels survive* |  |  |  |  |  |
| Group moderation of Path effects | Group x Path a, positive | L Precuneus | -4 | -58 | 20 | 64 | 1728 |
|  | Group x Path a, negative | L IFG p. Orbitalis / Area Fo3 | -26 | 26 | -14 | 9 | 243 |
|  | Group x Path b, Positive | *no voxels survive* |  |  |  |  |  |
|  | Group x Path b, Negative | *no voxels survive* |  |  |  |  |  |
|  | Group x Path ab, Positive | *no voxels survive* |  |  |  |  |  |
|  | Group x Path ab, Negative | *no voxels survive* |  |  |  |  |  |

^f^. This table presents uncorrected results from two voxelwise multilevel mediation analyses evaluating the effects of cues on subjective pain based on current contingencies (i.e. including reversals). All clusters are identified at a voxel-wise p-value of p < .001 (3 voxels at lowest threshold), contiguous with voxels at .005 and .01. Top rows report uncorrected results of each path across all participants, regardless of Group. Middle rows report Path a effects while controlling for Group. Bottom rows report clusters that are significantly moderated by Group. See Methods for additional details.

Table S6. Mediation of original cue contingencies: Small-volumes corrected results.^g^

| **Correction** | **Effect** | **Anatomical label** | **x** | **y** | **z** | **# of voxels** | **Volume (mm^3^)** |
| --- | --- | --- | --- | --- | --- | --- | --- |
| Pain modulatory network | Path a, positive | L Periaqueductal Gray | -4 | -32 | -10 | 10 | 270 |
|  |  | R Periaqueductal Gray | 8 | -28 | -8 | 4 | 108 |
|  |  | L IFG p. Orbitalis | -32 | 26 | -8 | 3 | 81 |
|  |  | L Putamen | -26 | -8 | 2 | 35 | 945 |
|  |  | R Thalamus | 4 | -16 | 8 | 79 | 2133 |
|  |  | R Putamen / Caudate | 20 | 4 | 10 | 5 | 135 |
|  |  | Thal: Visual | -22 | -32 | 8 | 5 | 135 |
|  | Path a, negative | *no voxels survive* |  |  |  |  |  |
|  | Path b, positive | L Hippocampus / Amygdala | -14 | -2 | -14 | 11 | 297 |
|  |  | L Superior Temporal Gyrus / anterior insula | -44 | 4 | -14 | 16 | 432 |
|  |  | R Putamen, contiguous with amygdala | 28 | 4 | -4 | 78 | 2106 |
|  |  | L Putamen | -26 | 14 | 2 | 38 | 1026 |
|  |  | R Insula Lobe | 38 | 26 | -4 | 21 | 567 |
|  |  | L Insula Lobe | -40 | 20 | 2 | 37 | 999 |
|  |  | R Rolandic Operculum / Area OP4 [PV] | 52 | -8 | 10 | 33 | 891 |
|  |  | R IFG p. Opercularis | 40 | 10 | 8 | 25 | 675 |
|  | Path b, negative | *no voxels survive* |  |  |  |  |  |
|  | Path ab, positive | *no voxels survive* |  |  |  |  |  |
|  | Path ab, negative | *no voxels survive* |  |  |  |  |  |
| Whole brain correction | Path a, positive | R Cerebelum IX / Lobule IX (Hem) | 2 | -50 | -52 | 8 | 216 |
|  |  | L IFG p. Orbitalis | -32 | 26 | -8 | 3 | 81 |
|  |  | R Superior Frontal Gyrus / Lateral PFC | 26 | 52 | 14 | 3 | 81 |
|  |  | R IFG p. Opercularis / DLPFC | 56 | 22 | 34 | 4 | 108 |
|  |  | R MCC / Area 5Ci (SPL) | 14 | -32 | 40 | 13 | 351 |
|  |  | L Superior Parietal Lobule | -14 | -76 | 46 | 22 | 594 |
|  | Path a, negative |  |  |  |  |  |  |
|  | Path b, positive | L Cerebelum Crus 2 / Lobule VIIa crusII (Hem) | -26 | -80 | -38 | 35 | 945 |
|  |  | R Cerebelum Crus 2 / Lobule VIIa crusII (Hem) | 26 | -86 | -40 | 3 | 81 |
|  |  | L Cerebelum Crus 1 / Lobule VIIa crusI (Hem) | -44 | -62 | -32 | 35 | 945 |
|  |  |  | -10 | -8 | -14 | 32 | 864 |
|  |  | L Temporal Pole | -50 | 10 | -8 | 47 | 1269 |
|  |  | R Putamen | 28 | 14 | 8 | 333 | 8991 |
|  |  | L Cuneus / Area hOc2 [V2] | 2 | -80 | 16 | 353 | 9531 |
|  |  | R hypothalamus | 4 | -2 | -8 | 13 | 351 |
|  |  | L Putamen | -26 | 10 | -2 | 62 | 1674 |
|  |  | R Insula Lobe | 38 | 22 | -4 | 9 | 243 |
|  |  | R Rolandic Operculum / Area OP3 [VS] | 50 | -8 | 14 | 114 | 3078 |
|  |  | R Thalamus | 28 | -16 | 14 | 31 | 837 |
|  |  | L IFG p. Opercularis | -40 | 8 | 10 | 5 | 135 |
|  |  | L ACC / Area 33 | -4 | 16 | 22 | 5 | 135 |
|  |  | L Superior Occipital Gyrus | -10 | -86 | 40 | 4 | 108 |
|  |  | L DMPFC | -14 | 22 | 38 | 16 | 432 |
|  |  | L DLPFC | -32 | 46 | 38 | 6 | 162 |
|  |  | R Cuneus / Area 7M (SPL) | 8 | -82 | 38 | 3 | 81 |
|  |  | L Inferior Parietal Lobule / Area hIP3 (IPS) | -32 | -58 | 40 | 4 | 108 |
|  |  | L Precentral Gyrus | -38 | 4 | 40 | 11 | 297 |
|  |  | L Superior Parietal Lobule / Area 7A (SPL) | -22 | -70 | 50 | 5 | 135 |
|  |  | Area 2 | 22 | -44 | 58 | 45 | 1215 |
|  |  | R Superior Parietal Lobule / Area 7P (SPL) | 22 | -76 | 56 | 4 | 108 |
|  |  | L Precuneus / Area 7A (SPL) | -10 | -70 | 58 | 5 | 135 |
|  |  | R Precuneus / Area 7P (SPL) | 10 | -70 | 56 | 4 | 108 |
|  |  | L Postcentral Gyrus / Area 3b | -32 | -38 | 62 | 11 | 297 |
|  |  | L Precuneus | -2 | -70 | 58 | 3 | 81 |
|  |  | RPrecentral Gyrus / Area 4a | 26 | -28 | 68 | 24 | 648 |
|  |  | L Postcentral Gyrus / Area 4a | -34 | -28 | 70 | 21 | 567 |
|  |  | L Paracentral Lobule / Area 4a | -10 | -28 | 74 | 10 | 270 |
|  |  | R Superior Parietal Lobule | 20 | -50 | 76 | 7 | 189 |
|  |  | L Superior Parietal Lobule / Area 5L (SPL) | -20 | -46 | 74 | 9 | 243 |
|  |  | RPrecentral Gyrus | 26 | -16 | 74 | 6 | 162 |
|  |  | L Postcentral Gyrus | -14 | -38 | 76 | 4 | 108 |
|  | Path b, negative | *no voxels survive* |  |  |  |  |  |
|  | Path ab, positive | *no voxels survive* |  |  |  |  |  |
|  | Path ab, negative | *no voxels survive* |  |  |  |  |  |

^g^. This table presents small-volume corrected results from a voxelwise multilevel mediation analysis evaluating the effects of cues on subjective pain based on original contingencies while controlling for reversals, regardless of Group. Top rows are FDR-corrected within pain modulatory networks (see Supplementary Figure S1) and bottom rows are whole-brain corrected. Both analyses are FDR-corrected at q < .05. See Methods for additional details.

Table S7. Mediation of original cue contingencies: Uncorrected results.^h^

| **Analysis** | **Effect** | **Anatomical label** | **x** | **y** | **z** | **# of voxels** | **Volume (mm^3^)** |
| --- | --- | --- | --- | --- | --- | --- | --- |
| All participants, regardless of Group | Path a, positive | R Cerebellum IX | 2 | -50 | -52 | 9 | 243 |
|  |  | R Thal: Premotor | 20 | -10 | 2 | 17 | 459 |
|  |  | R Thalamus | 4 | -16 | 8 | 101 | 2727 |
|  |  | R Putamen | 22 | 20 | 2 | 8 | 216 |
|  |  | L Putamen | -26 | -8 | 2 | 44 | 1188 |
|  |  | R Putamen | 22 | 2 | 10 | 39 | 1053 |
|  |  | R Superior Frontal Gyrus | 26 | 52 | 14 | 3 | 81 |
|  |  | L Middle Occipital Gyrus | -32 | -82 | 16 | 12 | 324 |
|  |  | L Middle Frontal Gyrus | -40 | 46 | 22 | 11 | 297 |
|  |  | R MCC | 14 | -28 | 40 | 19 | 513 |
|  |  | L Superior Parietal Lobule | -14 | -76 | 46 | 31 | 837 |
|  |  | R SupraMarginal Gyrus / Area PFm (IPL) | 58 | -40 | 44 | 11 | 297 |
|  | Path a, negative | L Rectal Gyrus / Area Fp2 | -4 | 46 | -16 | 74 | 1998 |
|  |  | R Hippocampus / CA2 | 26 | -16 | -16 | 25 | 675 |
|  |  | R Precuneus | 8 | -64 | 28 | 73 | 1971 |
|  |  | R Superior Temporal Gyrus / Area TE 3 | 68 | -10 | 8 | 9 | 243 |
|  |  | L Precuneus | -8 | -58 | 10 | 12 | 324 |
|  |  | L Superior Temporal Gyrus | -52 | -38 | 10 | 15 | 405 |
|  | Path b, positive | R Cerebelum VIII | 34 | -68 | -56 | 25 | 675 |
|  |  | L Cerebelum VIII | -32 | -56 | -50 | 39 | 1053 |
|  |  | Lobule VIIIb Verm | 4 | -70 | -52 | 5 | 135 |
|  |  | L Cerebelum Crus 2 | -28 | -80 | -38 | 51 | 1377 |
|  |  | L Cerebelum VI | -32 | -40 | -38 | 30 | 810 |
|  |  | L Cerebelum Crus 1 | -44 | -62 | -32 | 56 | 1512 |
|  |  | L Cerebelum VI | -26 | -76 | -22 | 16 | 432 |
|  |  | L Calcarine Gyrus / Area hOc2 [V2] | -2 | -80 | 14 | 475 | 12825 |
|  |  | L amygdala / parahippocampus | -10 | -4 | -14 | 40 | 1080 |
|  |  | R amygdala, putamen, insula (contiguous) | 32 | 4 | 14 | 827 | 22329 |
|  |  | L Insula Lobe | -44 | 10 | -4 | 88 | 2376 |
|  |  | R Temporal Pole | 44 | 10 | -16 | 13 | 351 |
|  |  | L Putamen | -26 | 10 | -2 | 82 | 2214 |
|  |  | R hypothalamus | 4 | -2 | -8 | 17 | 459 |
|  |  | R Calcarine Gyrus / Area hOc1 [V1] | 28 | -62 | 4 | 12 | 324 |
|  |  | R Thalamus: Temporal | 10 | -4 | 10 | 12 | 324 |
|  |  | L Caudate Nucleus | -16 | -4 | 22 | 23 | 621 |
|  |  | L Middle Occipital Gyrus | -28 | -82 | 22 | 5 | 135 |
|  |  | L Rolandic Operculum | -44 | -16 | 22 | 7 | 189 |
|  |  | R caudate | 20 | -2 | 26 | 12 | 324 |
|  |  | L Precentral Gyrus | -38 | 8 | 40 | 16 | 432 |
|  |  | L Superior Parietal Lobule / Area 7A (SPL) | -22 | -70 | 50 | 5 | 135 |
|  |  | R Area 2 | 22 | -44 | 58 | 50 | 1350 |
|  |  | L Posterior-Medial Frontal | -2 | 16 | 58 | 65 | 1755 |
|  |  | L Postcentral Gyrus / Area 4a | -32 | -32 | 62 | 56 | 1512 |
|  |  | L Precuneus / Area 7A (SPL) | -8 | -70 | 58 | 11 | 297 |
|  |  | R Superior Frontal Gyrus | 16 | -16 | 62 | 12 | 324 |
|  |  | RPrecentral Gyrus / Area 4a | 26 | -28 | 68 | 26 | 702 |
|  |  | RPrecentral Gyrus | 34 | -14 | 64 | 8 | 216 |
|  |  | R Superior Parietal Lobule / Area 5L (SPL) | 20 | -50 | 74 | 9 | 243 |
|  |  | L Paracentral Lobule / Area 4a | -10 | -28 | 74 | 22 | 594 |
|  |  | R Posterior-Medial Frontal | 8 | -2 | 74 | 11 | 297 |
|  |  | L Superior Parietal Lobule / Area 5L (SPL) | -20 | -46 | 74 | 9 | 243 |
|  |  | RPrecentral Gyrus | 26 | -16 | 74 | 7 | 189 |
|  | Path b, negative | L Hippocampus / DG | -26 | -16 | -22 | 39 | 1053 |
|  |  | L Inferior Temporal Gyrus | -58 | -26 | -22 | 14 | 378 |
|  |  | R Inferior Temporal Gyrus | 52 | -58 | -20 | 27 | 729 |
|  |  | R Middle Temporal Gyrus | 62 | -50 | -4 | 47 | 1269 |
|  |  | R Area hOc4v [V4v] | 32 | -76 | -8 | 19 | 513 |
|  |  | L Mid Orbital Gyrus / Area s32 | -8 | 34 | -14 | 10 | 270 |
|  |  | R Superior Frontal Gyrus / Area Fp1 | 20 | 70 | 2 | 14 | 378 |
|  |  | L Precuneus | -4 | -56 | 14 | 11 | 297 |
|  |  | R Angular Gyrus / Area PGp (IPL) | 52 | -62 | 32 | 34 | 918 |
|  |  | R Precuneus | 10 | -44 | 40 | 61 | 1647 |
|  | Path ab, positive | *no voxels survive* |  |  |  |  |  |
|  | Path ab, negative | R Cerebelum Crus 2 | 20 | -92 | -32 | 7 | 189 |
|  |  | R Inferior Temporal Gyrus / Area FG2 | 50 | -62 | -20 | 9 | 243 |
|  |  | L Superior Temporal Gyrus | -58 | -20 | 4 | 9 | 243 |
|  |  | R Rolandic Operculum / Area OP4 [PV] | 62 | -4 | 10 | 7 | 189 |
|  |  | L Cuneus / Area hOc3d [V3d] | -8 | -86 | 28 | 45 | 1215 |
|  |  | R SupraMarginal Gyrus / Area PFm (IPL) | 52 | -46 | 26 | 5 | 135 |
|  |  | L Precuneus | 2 | -58 | 28 | 12 | 324 |
|  |  | R Precuneus | 8 | -70 | 52 | 5 | 135 |
|  |  | R Thalamus | 8 | -14 | 8 | 42 | 1134 |
|  | Path a, negative | R Superior Temporal Gyrus / Area TE 3 | 68 | -26 | 8 | 9 | 243 |
|  | Path b, positive | L Cerebelum Crus 1 | -26 | -76 | -34 | 18 | 486 |
|  |  | L extended amygdala | -14 | -2 | -10 | 17 | 459 |
|  |  | L midbrain (substantia nigra) | -8 | -14 | -10 | 8 | 216 |
|  |  | R Putamen | 26 | 10 | 2 | 96 | 2592 |
|  |  | R IFG p. Opercularis | 40 | 16 | 10 | 89 | 2403 |
|  |  | R caudate | 28 | -16 | 14 | 31 | 837 |
|  | Path b, negative | *No voxels survive* |  |  |  |  |  |
|  | Path ab, positive | *No voxels survive* |  |  |  |  |  |
|  | Path ab, negative | *No voxels survive* |  |  |  |  |  |
| Group moderation of Path effects | Group x Path a, positive | *No voxels survive* |  |  |  |  |  |
|  | Group x Path a, negative | *No voxels survive* | 26 | 14 | 16 | 9 | 243 |
|  | Group x Path b, positive | *No voxels survive* |  |  |  |  |  |
|  | Group x Path b, negative | *No voxels survive* |  |  |  |  |  |
|  | Group x Path ab, positive | Area 7A SPL | -22 | -70 | 64 | 5 | 135 |
|  | Group x Path ab, negative | *No voxels survive* |  |  |  |  |  |

^h^. This table presents uncorrected results from two voxelwise multilevel mediation analyses evaluating the effects of cues on subjective pain based on original contingencies while controlling for reversals. All clusters are identified at a voxel-wise p-value of p < .001 (3 voxels at lowest threshold), contiguous with voxels at .005 and .01. Top rows report uncorrected results of each path across all participants, regardless of Group. Middle rows report Path a effects while controlling for Group. Bottom rows report clusters that are significantly moderated by Group. See Methods for additional details.

Table S8. Associations with expected value (EV): Small-volumes corrected results.^i^

| **Correction** | **Analysis** | **Effect** | **Anatomical label** | **x** | **y** | **z** | **# of voxels** | **Volume (mm^3^)** |
| --- | --- | --- | --- | --- | --- | --- | --- | --- |
| Pain modulatory network | Instructed Group | Positive association with EV | Right anterior insula | 34 | 16 | -16 | 4 | 108 |
|  |  | Negative association with EV | *No voxels survive* |  |  |  |  |  |
|  | Uninstructed Group | Positive association with EV | *No voxels survive* |  |  |  |  |  |
|  |  | Negative association with EV | Left rostrodorsal ACC | -4 | 32 | 14 | 4 | 108 |
|  | Main effect of EV, controlling for Group | Positive association with EV | *No voxels survive* |  |  |  |  |  |
|  |  | Negative association with EV | *No voxels survive* |  |  |  |  |  |
|  | Group differences in EV (Instructed - Uninstructed) | Positive effect | Left rostrodorsal ACC | -8 | 40 | 2 | 39 | 1053 |
|  |  | Negative effect | *No voxels survive* |  |  |  |  |  |
|  | Instructed Group | Positive association with EV | R Mid Orbital Gyrus / Area Fp2 / VMPFC | 4 | 56 | -4 | 18 | 486 |
|  |  |  | R ACC / DMPFC | 8 | 44 | 26 | 7 | 189 |
|  |  |  | L Middle Frontal Gyrus / DLPFC | -32 | -2 | 52 | 6 | 162 |
|  |  | Negative association with EV | *No voxels survive* |  |  |  |  |  |
|  | Uninstructed Group | Positive association with EV | R IFG p. Triangularis / Area 45 / anterior insula | 50 | 22 | -2 | 8 | 216 |
|  |  | Negative association with EV | R anterior PFC | 22 | 46 | 2 | 8 | 216 |
|  |  |  | R Precuneus | 4 | -52 | 16 | 20 | 540 |
|  | Main effect of EV, controlling for Group | Positive association with EV | L Middle Frontal Gyrus / DLPFC | -28 | 28 | 32 | 3 | 81 |
|  |  | Negative association with EV | R anterior PFC | 20 | 46 | 2 | 13 | 351 |
|  |  |  | L Superior Temporal Gyrus / Area PFcm (IPL) | -46 | -40 | 16 | 16 | 432 |
|  |  |  | L inferior frontal gyrus | -34 | 22 | 20 | 5 | 135 |
|  | Group differences in EV (Instructed - Uninstructed) | Positive effect | L Medial Temporal Pole | -32 | 14 | -38 | 7 | 189 |
|  |  |  | L Middle Temporal Gyrus | -58 | -16 | -16 | 13 | 351 |
|  |  |  | R Mid Orbital Gyrus / Area Fp2 / MPFC | 2 | 58 | -2 | 60 | 1620 |
|  |  |  | L ACC | -10 | 38 | 14 | 18 | 486 |
|  |  |  | R Precuneus | 8 | -44 | 16 | 5 | 135 |
|  |  |  | L Superior Temporal Gyrus / Area PFcm (IPL) / TPJ | -46 | -40 | 16 | 6 | 162 |
|  |  |  | Middle cingulate cortex | 4 | 8 | 22 | 18 | 486 |
|  |  | Negative effect | R Cerebelum VIII | 34 | -70 | -52 | 3 | 81 |
|  |  |  | R Rectal Gyrus / Area Fo1/ mOFC | 10 | 44 | -20 | 15 | 405 |
|  |  |  | L IFG p. Opercularis / Area 44 | -56 | 14 | 10 | 5 | 135 |
|  |  |  | L Inferior Parietal Lobule / Area PF (IPL) | -56 | -46 | 44 | 29 | 783 |
|  |  |  | L Precentral Gyrus / Area 44 / DLPFC | -50 | 8 | 40 | 35 | 945 |
|  |  |  | R Superior Frontal Gyrus / DMPFC | 26 | 22 | 44 | 2 | 54 |

^i^. This table presents group results from voxelwise analyses of associations between expected value and brain activation on medium heat, as measured by AUC estimates (see Methods). Group results were analyzed using robust regression. Top rows are FDR-corrected within pain modulatory networks (see Supplementary Figure S1) and bottom rows are whole-brain corrected. Both analyses are FDR-corrected at q < .05. See Methods for additional details.

Table S9. Associations with expected value (EV): Uncorrected results.^j^

| **Analysis** | **Effect** | **Anatomical label** | **x** | **y** | **z** | **# of voxels** | **Volume (mm^3^)** |
| --- | --- | --- | --- | --- | --- | --- | --- |
| Instructed Group | Positive association with EV | L anterior insula contiguous with amygdala | -34 | 4 | -8 | 28 | 756 |
|  |  | R Mid Orbital Gyrus / Area Fp2 / MPFC | 4 | 58 | -4 | 28 | 756 |
|  |  | R Caudate Nucleus | 10 | 16 | 8 | 16 | 432 |
|  |  | R ACC / DMPFC | 8 | 44 | 26 | 9 | 243 |
|  |  | L MCC | -8 | -40 | 44 | 13 | 351 |
|  | Negative association with EV | R Inferior Temporal Gyrus | 50 | -44 | -14 | 8 | 216 |
| Uninstructed Group | Positive association with EV | L ParaHippocampal Gyrus / Subiculum | -28 | -14 | -26 | 7 | 189 |
|  |  | L Cerebelum VI | -8 | -68 | -20 | 7 | 189 |
|  |  | L Superior Medial Gyrus | -2 | 64 | 2 | 20 | 540 |
|  |  | R Anterior PFC | 26 | 46 | 2 | 12 | 324 |
|  |  | R Calcarine Gyrus | 4 | -56 | 14 | 30 | 810 |
|  | Negative association with EV | L ParaHippocampal Gyrus / Subiculum | -28 | -14 | -26 | 7 | 189 |
|  |  | L Cerebelum VI | -8 | -68 | -20 | 7 | 189 |
|  |  | L Superior Medial Gyrus / Area Fp2 | -2 | 64 | 2 | 20 | 540 |
|  |  | R Anterior PFC | 26 | 46 | 2 | 12 | 324 |
|  |  | R Calcarine Gyrus | 4 | -56 | 14 | 30 | 810 |
| Main effect of EV, controlling for Group | Positive association with EV | R IFG p. Triangularis | 46 | 22 | 2 | 16 | 432 |
|  |  | L SupraMarginal Gyrus | -44 | -44 | 28 | 4 | 108 |
|  |  | R MCC | 8 | -22 | 40 | 18 | 486 |
|  |  | L Precentral Gyrus / DLPFC | -44 | 4 | 44 | 39 | 1053 |
|  |  | R Postcentral Gyrus / Area 2 | 50 | -22 | 46 | 11 | 297 |
|  |  | L Supplementary Motor Area | -10 | -20 | 56 | 9 | 243 |
|  | Negative association with EV | R Rectal Gyrus / Area Fo1 | 10 | 44 | -20 | 19 | 513 |
|  |  | R IFG p. Opercularis / Area 45 / lateral PFC | 56 | 20 | 32 | 27 | 729 |
|  |  | L Inferior Parietal Lobule / Area PF (IPL) | -56 | -46 | 44 | 45 | 1215 |
|  |  | L Precentral Gyrus / Area 44 / DLPFC | -50 | 8 | 40 | 47 | 1269 |
| Group differences in EV (Instructed - Uninstructed) | Positive effect | R Periaqueductal Gray | 2 | -34 | -20 | 6 | 162 |
|  |  | L Middle Temporal Gyrus | -58 | -16 | -16 | 14 | 378 |
|  |  | L ACC / MPFC | -4 | 50 | 4 | 165 | 4455 |
|  |  | R Superior Orbital Gyrus / Area Fp1 | 28 | 64 | -8 | 5 | 135 |
|  |  | L Precuneus | -4 | -56 | 20 | 150 | 4050 |
|  |  | L Heschls Gyrus / Area Ig2 | -40 | -20 | 10 | 28 | 756 |
|  | Negative effect | R Rectal Gyrus / Area Fo1 / VMPFC | 10 | 44 | -20 | 19 | 513 |
|  |  | R IFG p. Opercularis / Area 45 | 56 | 20 | 32 | 27 | 729 |
|  |  | L Inferior Parietal Lobule / Area PF (IPL) | -56 | -46 | 44 | 45 | 1215 |
|  |  | L Precentral Gyrus / Area 44 / DLPFC | -50 | 8 | 40 | 47 | 1269 |

^j^. This table presents group results from voxelwise analyses of associations between expected value and brain activation on medium heat, as measured by AUC estimates (see Methods). Group results were analyzed using robust regression. All clusters are identified at a voxel-wise p-value of p < .001 (3 voxels at lowest threshold), contiguous with voxels at .005 and .01. See Methods for additional details.

Table S10. Comparing instructed and feedback-driven expected value (EV) within the Instructed Group: Small volumes corrected results.^k^

| **Correction** | **Analysis** | **Effect** | **Anatomical label** | **x** | **y** | **z** | **# of voxels** | **Volume (mm^3^)** |
| --- | --- | --- | --- | --- | --- | --- | --- | --- |
| Pain modulatory network | Instructed vs Uninstructed EV | Positive association (Instructed > Uninstructed) | *No voxels survive* |  |  |  |  |  |
|  |  | Negative association (Uninstructed > Instructed) | *No voxels survive* |  |  |  |  |  |
|  | Instructed EV, controlling for uninstructed | Positive association with EV | *No voxels survive* |  |  |  |  |  |
|  |  | Negative association with EV | *No voxels survive* |  |  |  |  |  |
|  | Uninstructed EV, controlling for instructed | Positive association with EV | *No voxels survive* |  |  |  |  |  |
|  |  | Negative association with EV | *No voxels survive* |  |  |  |  |  |
| Whole brain correction | Instructed vs Uninstructed EV | Positive association (Instructed > Uninstructed) | *No voxels survive* |  |  |  |  |  |
|  |  | Negative association (Uninstructed > Instructed) | *No voxels survive* |  |  |  |  |  |
|  | Instructed EV, controlling for Uninstructed EV | Positive association with EV | R Substantia Nigra | 8 | -16 | -8 | 9 | 243 |
|  |  | Negative association with EV | RPrecentral Gyrus | 46 | -22 | 62 | 4 | 108 |
|  | Uninstructed EV, controlling for instructed | Positive association with EV | *No voxels survive* |  |  |  |  |  |
|  |  | Negative association with EV | *No voxels survive* |  |  |  |  |  |

^k^. This table presents group results from voxelwise analyses of associations between expected value and brain activation on medium heat within the Instructed Group (see Methods). Group results were analyzed using robust regression. Top rows are FDR-corrected within pain modulatory networks (see Supplementary Figure S1) and bottom rows are whole-brain corrected. Both analyses are FDR-corrected at q < .05. See Methods for additional details.

Table S11. Comparing instructed and feedback-driven expected value (EV) within the Instructed Group: Uncorrected results.^l^

| **Analysis** | **Effect** | **Anatomical label** | **x** | **y** | **z** | **# of voxels** | **Volume (mm^3^)** |
| --- | --- | --- | --- | --- | --- | --- | --- |
| Instructed vs Uninstructed EV | Positive association (Instructed > Uninstructed) | L amygdala | -14 | 2 | -16 | 14 | 378 |
|  | Negative association (Uninstructed > Instructed) | L Fusiform Gyrus | -26 | -34 | -20 | 18 | 486 |
| Instructed EV, controlling for uninstructed | Positive association | L cerebellum | -34 | -56 | -40 | 10 | 270 |
|  |  | R Putamen / extended amygdala | 20 | 10 | -8 | 15 | 405 |
|  |  | R Midbrain (substantia nigra) | 8 | -16 | -8 | 9 | 243 |
|  |  | L rostral ACC / MPFC | -4 | 50 | 10 | 6 | 162 |
|  |  | L PCC | 2 | -40 | 28 | 17 | 459 |
|  |  | R rdACC | 10 | 40 | 28 | 10 | 270 |
|  | Negative association | RPrecentral Gyrus | 46 | -22 | 62 | 4 | 108 |
| Uninstructed EV, controlling for instructed | Positive association | R MCC | 8 | -2 | 32 | 8 | 216 |
|  |  | RPrecentral Gyrus | 50 | -10 | 50 | 3 | 81 |
|  | Negative association | R Cerebelum Crus 2 | 32 | -80 | -38 | 13 | 351 |
|  |  | L ParaHippocampal Gyrus / amygdala | -14 | 2 | -20 | 13 | 351 |
|  |  | L Superior Medial Gyrus | 2 | 40 | 38 | 21 | 567 |

^l^. This table presents group results from voxelwise analyses of associations between expected value and brain activation on medium heat within the Instructed Group (see Methods). Group results were analyzed using robust regression. All clusters are identified at a voxel-wise p-value of p < .001 (3 voxels at lowest threshold), contiguous with voxels at .005 and .01. See Methods for additional details.
